## Supplementary information for "A novel nematode species from the Siberian permafrost shares adaptive mechanisms for cryptobiotic survival with *C. elegans* dauer larva"

### Supplementary Tables

**Supplementary Table 1. Morphometrics of *Panagrolaimus n. sp.***

| **Characters** | **Holotype female** | **All the type female specimens** | | | | |
|---|---|---|---|---|---|---|
| | | **n** | **min–max** | **mean** | **SD** | **CV** |
| Body length, µm | 944 | 19 | 813–979 | 896 | 35.3 | 3.93 |
| Body diameter at the level of anterior sensilla, µm | 8 | 19 | 6.5–8 | 7.4 | 0.47 | 6.35 |
| Body diameter at the level of nerve ring, µm | 29 | 19 | 24–33 | 29 | 1.99 | 6.78 |
| Body diameter at the level of cardia, µm | 31 | 19 | 30–35 | 33 | 1.59 | 4.88 |
| Body diameter at the level of midbody, µm | 44 | 19 | 37–44 | 40 | 1.82 | 4.52 |
| Body diameter at the level of anus, µm | 24 | 19 | 16–27 | 23 | 2.25 | 9.66 |
| a | 21.5 | 19 | 21–26 | 22 | 1.26 | 5.69 |
| b | 6.84 | 18 | 5.66–6.84 | 6.18 | 0.31 | 5.02 |
| c | 17.8 | 16 | 16–21 | 18 | 1.52 | 8.45 |
| c’ | 2.18 | 16 | 1.78–2.58 | 2.12 | 0.19 | 8.96 |
| V. % | 61.3 | 18 | 57–68 | 61 | 2.22 | 3.67 |
| Procorpus length, µm | 98 | 18 | 85–103 | 93 | 5.71 | 6.12 |
| Terminal bulb length, µm | 26 | 19 | 23–33 | 26 | 2.48 | 9.51 |
| Terminal bulb width, µm | 20 | 19 | 17–21 | 18 | 1.13 | 6.11 |

Remarks: a – body length divided by maximum body diameter; b – body length divided by length of the pharynx; c – body length divided by tail length; c’ – tail length divided by anal body diameter; CV – coefficient of variation; min–max – range; n – number of individuals measured; SD – standard deviation; V – distance of vulva from anterior end as percentage of body length (%).

**Supplementary Table 2. All taxa used for inferring the species tree**. Accession numbers for each gene are displayed. Members of *Panagrolaimus* who form a monophyletic clade with definitive *Propanagrolaimus* taxa have been renamed, and are designated with "*".

| Taxon | 18S Accession number | 28S Accession number |
|---|---|---|
| *Ancestral panagrolaimus* | - | - |
| *Panagrolaimus sp. ES6* | FJ590961.1 | FJ590996.1 |
| *Panagrolaimus rigidus* | FJ590974.1 | FJ591009.1 |
| *Panagrolaimus sp. ES2* | FJ590962.1 | FJ590997.1 |
| *Panagrolaimus sp. ES1* | FJ590960.1 | FJ590995.1 |
| *Panagrolaimus superbus* | FJ590973.1 | FJ591008.1 |
| *Panagrolaimus sp. ES5* | FJ590972.1 | FJ591007.1 |
| *Panagrolaimus sp. PS443* | FJ590978.1 | FJ591013.1 |
| *Panagrolaimus sp. DL0180* | FJ590990.1 | FJ591025.1 |
| *Panagrolaimus sp. DL0128* | FJ590987.1 | FJ591022.1 |
| *Panagrolaimus sp. ES3* | FJ590963.1 | FJ590998.1 |
| *Panagrolaimus sp. DL0117* | FJ590986.1 | FJ591021.1 |
| *Panagrolaimus sp. PS1579* | FJ590976.1 | FJ591011.1 |
| *Panagrolaimus sp. PS3966* | FJ590983.1 | FJ591018.1 |
| *Panagrolaimus davidi* | FJ590981.1 | FJ591016.1 |
| *Panagrolaimus sp. PS1159* | FJ590977.1 | FJ591012.1 |
| *Panagrolaimus sp. DL0139* | FJ590989.1 | FJ591024.1 |
| *Panagrolaimus sp. DL0137* | FJ590988.1 | FJ591023.1 |
| *Panagrolaimus sp. DL0072* | FJ590985.1 | FJ591020.1 |
| *Panagrolaimus sp. DL0050* | FJ590984.1 | FJ591019.1 |
| *Panagrolaimus sp. JB051* | FJ590971.1 | FJ591006.1 |
| *Panagrolaimus sp. SN103* | FJ590970.1 | FJ591005.1 |
| *Panagrolaimus sp. JB115* | FJ590969.1 | FJ591004.1 |
| *Panagrolaimus sp. JB131* | FJ590968.1 | FJ591003.1 |
| *Panagrolaimus sp. PS1162* | FJ590958.1 | FJ590993.1 |
| *Panagrolaimus sp. PS1806* | FJ590957.1 | FJ590992.1 |
| *Propanagrolaimus sp. PS1732** | FJ590959.1 | FJ590994.1 |
| *Propanagrolaimus sp. JU765** | FJ590956.1 | FJ590991.1* |
| *Propanagrolaimus sp. WTM1* | KJ434176.1 | KJ434176.1 |
| *Propanagrolaimus sp. LC91* | KJ434175.1 | KJ434175.1 |
| *Propanagrolaimus detritophagus** | FJ590980.1 | FJ591015.1 |
| *Propanagrolaimus paetzoldi** | FJ590979.1 | FJ591014.1 |
| *Panagrellus redivivus* | AF083007.1 | KU180687.1 |
| *Halicephalobus gingivalis* | MK087058.1 | DQ145685.1 |
| *Halicephalobus sp. AA4* | MF470238.1 | MG051261.1 |
| *Halicephalobus sp. FTL-5A-2* | MF470221.1 | MG051244.1 |
| *Halicephalobus sp. G228* | MF470235.1 | MG051259.1 |
| *Halicephalobus sp. KW-1A-7* | MF470228.1 | MG051253.1 |
| *Halicephalobus sp. SRC-14A* | MF470211.1 | MG051231.1 |

| | | |
|---|---|---|
| *Halicephalobus mephisto* | Mined from genome | Mined from genome |
| *Halicephalobus sp. LD6* | FJ590965.1 | FJ591000.1 |
| *Halicephalobus sp. LD8* | FJ590967.1 | FJ591002.1 |
| *Halicephalobus sp. LD7* | FJ590966.1 | FJ591001.1 |
| *Halicephalobus sp. LD3* | FJ590964.1 | FJ590999.1 |
| *Caenorhabditis briggsae* | U13929.1 | FJ591017.1 |

**Supplementary Table 3: Estimated repeat content of sequences from *Panagrolaimus n. sp.* genome assembly.** A total of 22.3% of the *Panagrolaimus n. sp.* genome assembly consists of repetitive sequences. Simple repeats comprise the largest portion of the repetitive genome.

| Repeat class | % of the assembly |
|---|---|
| SINE | 0.11% |
| LINE | 0.23% |
| LTR | 0.73% |
| DNA | 0.97% |
| Simple repeats | 14.30% |
| Unclassified repeats | 7.23% |
| TOTAL | 22.29% |

**Supplementary Table 4.** All 60 genes used for phylogenetic analyses and their best-fit models of sequence evolution.

| **Gene** | **Best fit model** |
|---|---|
| ahcy-1 | TPM2+F+G4 |
| alh-8 | TIM2e+I+G4 |
| aps-3 | TIM3+F+R5 |
| arx-2 | GTR+F+I+G4 |
| atg-3 | TIM3e+I+G4 |
| bud-31 | SYM+I+G4 |
| calm-1 | SYM+I+G4 |
| ccb-1 | TIM2+F+I+G4 |
| cni-1 | TIM2+F+I+G4 |
| csq-1 | TIM3+F+I+G4 |
| dlc-1 | TIM3e+I+G4 |
| dnj-30 | TIM3+F+G4 |
| eif-6 | TN+F+G4 |
| fbp-1 | GTR+F+I+G4 |
| gpa-12 | GTR+F+G4 |
| hint-1 | GTR+F+I+G4 |
| inf-1 | TIM2+F+I+G4 |
| let-522 | K2P+G4 |
| let-60 | TNe+I+G4 |
| mdh-1 | TIM2+F+I+G4 |
| mlt-8 | TPM2u+F+I+G4 |
| ndk-1 | TIM3+F+R4 |
| nuo-1 | TVMe+R4 |
| oig-2 | GTR+F+I+G4 |
| oig-3 | TVMe+I+G4 |
| pas-4 | TNe+I+G4 |
| pbs-3 | TIM2+F+R5 |
| pbs-5 | TPM2u+F+I+G4 |
| plpr-1 | TIM2+F+R4 |
| rab-14 | TIM2e+I+G4 |
| rab-7 | TIM2e+G4 |
| ran-1 | TIM3+F+G4 |
| rap-2 | HKY+F+I+G4 |
| ric-4 | TIM2e+R4 |
| rpb-10 | GTR+F+R5 |
| rpl-12 | TIM2+F+I+G4 |
| rpl-15 | SYM+I+G4 |
| rpl-2 | TNe+I+G4 |
| rpl-26 | GTR+F+I+G4 |
| rpl-31 | TIM3e+I+G4 |
| rpl-9 | HKY+F+I+G4 |
| rps-13 | TPM2+F+I+G4 |
| rps-15 | TIM2+F+I+G4 |

| | |
|---|---|
| rps-20 | TIM2e+I+G4 |
| rps-3 | GTR+F+I+G4 |
| rps-9 | TIM2+F+I+G4 |
| sar-1 | TIM2e+I+G4 |
| sftb-1 | TNe+G4 |
| snr-7 | GTR+F+R5 |
| snx-3 | TIM2+F+R4 |
| tin-10 | TIM2+F+I+G4 |
| tpi-1 | HKY+F+G4 |
| uba-5 | GTR+F+R5 |
| ufc-1 | TN+F+I+G4 |
| ufm-1 | TIM2+F+I+G4 |
| unc-69 | TIM3+F+I+G4 |
| usp-46 | GTR+F+R4 |
| vha-10 | TPM2+F+I+G4 |
| vha-14 | TIM2+F+I+G4 |
| vha-4 | SYM+R4 |

**Supplementary Table 5.** All genomes used for phylogenetic analysis, their class, and the number of genes used for each species are displayed.

| | | |
|---|---|---|
| *Acanthocheilonema viteae* | Nematoda | 33 |
| *Acrobeloides nanus* | Nematoda | 55 |
| *Ancylostoma caninum* | Nematoda | 68 |
| *Ancylostoma ceylanicum* | Nematoda | 87 |
| *Ancylostoma duodenale* | Nematoda | 57 |
| *Angiostrongylus cantonensis* | Nematoda | 65 |
| *Angiostrongylus costaricensis* | Nematoda | 78 |
| *Anisakis simplex* | Nematoda | 44 |
| *Ascaris lumbricoides* | Nematoda | 45 |
| *Ascaris suum* | Nematoda | 46 |
| *Brugia malayi* | Nematoda | 92 |
| *Brugia pahangi* | Nematoda | 40 |
| *Brugia timori* | Nematoda | 13 |
| *Bursaphelenchus xylophilus* | Nematoda | 57 |
| *Caenorhabditis elegans* | Nematoda | 133 |
| *Caenorhabditis remanei* | Nematoda | 129 |
| *Caenorhabditis sp34* | Nematoda | 131 |
| *Dictyocaulus viviparus* | Nematoda | 76 |
| *Diploscapter coronatus* | Nematoda | 104 |
| *Diploscapter pachys* | Nematoda | 91 |
| *Dirofilaria immitis* | Nematoda | 37 |
| *Ditylenchus destructor* | Nematoda | 44 |
| *Ditylenchus dipsaci* | Nematoda | 30 |
| *Dracunculus medinensis* | Nematoda | 34 |
| *Elaeophora elaphi* | Nematoda | 37 |
| *Enterobius vermicularis* | Nematoda | 42 |
| *Globodera pallida* | Nematoda | 27 |
| *Globodera rostochiensis* | Nematoda | 37 |
| *Gongylonema pulchrum* | Nematoda | 10 |
| *Haemonchus contortus* | Nematoda | 103 |
| *Haemonchus placei* | Nematoda | 80 |
| *Halicephalobus mephisto* | Nematoda | 63 |
| *Heligmosomoides polygyrus* | Nematoda | 77 |
| *Heterodera glycines* | Nematoda | 37 |
| *Heterorhabditis bacteriophora* | Nematoda | 17 |
| *Litomosoides sigmodontis* | Nematoda | 42 |
| *Loa loa* | Nematoda | 129 |
| *Meloidogyne arenaria* | Nematoda | 41 |
| *Meloidogyne enterolobii* | Nematoda | 24 |
| *Meloidogyne floridensis* | Nematoda | 11 |
| *Meloidogyne graminicola* | Nematoda | 20 |
| *Meloidogyne hapla* | Nematoda | 24 |
| *Meloidogyne incognita* | Nematoda | 37 |
| *Meloidogyne javanica* | Nematoda | 39 |

| | | |
|---|---|---|
| *Mesorhabditis belari* | Nematoda | 91 |
| *Micoletzkya japonica* | Nematoda | 65 |
| *Necator americanus* | Nematoda | 130 |
| *Nippostrongylus brasiliensis* | Nematoda | 74 |
| *Oesophagostomum dentatum* | Nematoda | 42 |
| *Onchocerca flexuosa* | Nematoda | 20 |
| *Onchocerca ochengi* | Nematoda | 33 |
| *Onchocerca volvulus* | Nematoda | 46 |
| *Oscheius tipulae* | Nematoda | 97 |
| *Panagrellus redivivus* | Nematoda | 65 |
| *Panagrolaimus superbus* | Nematoda | 37 |
| *Panagrolaimus sp. DL0137* | Nematoda | 59 |
| *Panagrolaimus sp. ES5* | Nematoda | 55 |
| *Panagrolaimus sp. PS1159* | Nematoda | 54 |
| *Panagrolaimus sp. PS1579* | Nematoda | 52 |
| *Parapristionchus giblindavisi* | Nematoda | 62 |
| *Parascaris univalens* | Nematoda | 59 |
| *Parastrongyloides trichosuri* | Nematoda | 47 |
| *Plectus sambesii* | Nematoda | 49 |
| *Pristionchus arcanus* | Nematoda | 69 |
| *Pristionchus entomophagus* | Nematoda | 65 |
| *Pristionchus exspectatus* | Nematoda | 68 |
| *Pristionchus fissidentatus* | Nematoda | 72 |
| *Pristionchus japonicus* | Nematoda | 74 |
| *Pristionchus maxplancki* | Nematoda | 70 |
| *Pristionchus mayeri* | Nematoda | 65 |
| *Pristionchus pacificus* | Nematoda | 74 |
| *Propanagrolaimus sp. JU765* | Nematoda | 68 |
| *Romanomermis culicivorax* | Nematoda | 13 |
| *Soboliphyme baturini* | Nematoda | 11 |
| *Steinernema carpocapsae* | Nematoda | 66 |
| *Steinernema feltiae* | Nematoda | 69 |
| *Steinernema glaseri* | Nematoda | 67 |
| *Steinernema monticolum* | Nematoda | 61 |
| *Steinernema scapterisci* | Nematoda | 73 |
| *Strongyloides papillosus* | Nematoda | 46 |
| *Strongyloides ratti* | Nematoda | 44 |
| *Strongyloides stercoralis* | Nematoda | 38 |
| *Strongyloides venezuelensis* | Nematoda | 43 |
| *Strongylus vulgaris* | Nematoda | 20 |
| *Syphacia muris* | Nematoda | 39 |
| *Teladorsagia circumcincta* | Nematoda | 43 |
| *Thelazia callipaeda* | Nematoda | 40 |
| *Toxocara canis* | Nematoda | 49 |
| *Trichinella britovi* | Nematoda | 13 |
| *Trichinella murrelli* | Nematoda | 12 |
| *Trichinella nativa* | Nematoda | 11 |
| *Trichinella nelsoni* | Nematoda | 13 |

| | | |
|---|---|---|
| *Trichinella papuae* | Nematoda | 13 |
| *Trichinella patagoniensis* | Nematoda | 14 |
| *Trichinella spiralis* | Nematoda | 103 |
| *Trichinella zimbabwensis* | Nematoda | 11 |
| *Trichuris muris* | Nematoda | 17 |
| *Trichuris suis* | Nematoda | 11 |
| *Trichuris trichiura* | Nematoda | 15 |
| *Wuchereria bancrofti* | Nematoda | 29 |
| *Macrostomum lignano* | Platyhelminthes | 11 |

### Supplementary methods

#### Sampling site

The sampling site was in the Kolyma Lowland of northeast Siberia, Russia, in the continuous permafrost zone (Fig.1a). Sample P-1320 was collected by Dr.S.Gubin in summer 2002 from the frozen wall of the outcrop Duvanny Yar located on the right bank of the Kolyma River (68° 37′ 739″ N, 159° 11′ 678″ E). The outcrop Duvanny Yar is a cliff about 9 km long and about 55 m high above the river water level and is described in detail as a key section of late Pleistocene deposits (Ice complex) in the northeastern Arctic (Fig.1b)[1,2] (Strauss, 2010;) These deposits accumulated syngenetically, i.e., simultaneously freezing from below and never thawed since the moment of formation. Frozen silty sediments include sandy alluvial layers, peat lenses, buried paleosols, Pleistocene rodent burrows and wide variety of plant and vertebrate fossils (Fig.1c)[3,4]. The permafrost deposits have high ice content (35–80%) and mean annual temperature varies from -3°C to -11°C depending on the depth[5]. Maximal depth of seasonal thawing of permafrost-affected soils at the sites does not exceed 70 cm, while permafrost thickness is over 500 meters[2].

#### Fossil rodent burrows

Studied burrow P-1320 was taken from the frozen outcrop wall at a depth of about 40 m below the surface and about 11 m above river water level in undisturbed and never thawed late Pleistocene permafrost deposits. The fossil burrow left by arctic gophers *Citellus (Urocitellus) parrii* Richardson, 1825, consists of entrance tunnel and a large nesting chamber up to 25 cm in diameter (Fig.1d)[6]. The material of fossil burrows represented by the well-preserved remnants of herbaceous plants, included seeds of higher plants, rodent bones and coprolites, insects, feathers, hairs of large animals and other material brought from the surface, and are part of the biogeocenoses formed in the immediate vicinity of the rodent's dwelling. Biological material in the ancient burrows is quite well preserved due to fastest burial and freezing into aggrading permafrost sediment[4].

#### Sampling procedure

Sampling, storage and transportation of the frozen samples under sterile conditions, as well as contamination testing were performed using protocols that have previously been described for microbiological purposes[7,8,9,10]. Samples were taken from the frozen outcrops wall. Melting parts of burrows were removed until the unthawed layer exposed. The whole ice-cemented chamber was removed from the frozen sediment, placed in the plastic bag and kept frozen during field and transportation to the laboratory in Pushchino. In the laboratory surface of frozen chamber was cleaned, the internal part was aseptically separated and divided into pieces inside a laminar airflow cabinet, placed in sterile plastic bags and stored at -20°C until analyses.

**Age of *Panagrolaimus n. sp.***

According the radiocarbon dating, stratigraphic and spore-pollen analysis the permafrost deposits of Duvanny Yar were accumulated under tundra-steppe landscape during the marine isotopic stages (MIS) 2 and 3 (15.5 - 55 kya BP)[2]. Earlier, more than 30 similar burrows of fossil rodents were described in the layer of permafrost sediments at depths of 10-20 m above river level [3,4]. Series of radiocarbon dating of animal and plant material from permafrost sediments, buried borrows and paleosols from this layer were obtained previously, amounting between about 30 and up to 50 kyr BP depending on the depth[11,10,2,12]. In the current study the Accelerator Mass Spectrometry (AMS) radiocarbon dating of plant material obtained from the burrow P-1320 was provided in the Laboratory for Radiocarbon Dating and Electron Microscopy, Institute of Geography, RAS, and determined a direct $^{14}C$ age of 44,315±405 BP ($IGAN_{AMS}$ 9137) and calibrated age range is 45,839 – 47,769 cal BP (95.4% probability) (Fig.S1).

It was shown that below the seasonally thawing layer, which is no more than 0.7 m in the study sites, any migration or thermal diffusion of bacterial cells with unfrozen water films within the stratum is precluded[13,14,8]. Permafrost deposits contained burrows were firmly cemented by ice and showed no signs of degradation or thawing of permafrost in the past. Thus, the vertical movement of modern microorganisms and contamination syncryogenic strata is impossible and age of biota in corresponds to the age of permafrost sediments[15]. Natural melting of permafrost leads to the destruction of the outcrops wall during the warm season at a rate of about a meter per month, exposing the new frozen deposits and continually revealing new buried paleontological objects, for example well-preserved mammal and other ancient animal mummies.

**Nematode isolation and culturing**

Revitalization of cultivable ancient nematodes was observed during enrichment cultivation with *E. coli* strain OP50 as a food source. Samples of about 1cm³ were taken from the inner part of the frozen burrows under sterile conditions (inside laminar air flow cabinet) and resuspended in mineral liquid Prescott–James medium (PJM) into Petri dishes[16]. To avoid drying and cross-contamination, the plates were wrapped with Parafilm and cultivated at 20ºC for several weeks. The soil suspensions were observed every 2-3 days in closed plates using an inverted light microscope Nikon Eclipse E100 to finding revived organisms until motile nematodes were detected. Clonal strains were established from individual nematodes which were picked from the soil suspension, transferred to fresh liquid PJM plates and grown at 20ºC using *E. coli* strain OP50 as a food source. Strain *E. coli* was grown in the tryptic soy broth agar (1/2 TSB) overnight at +35°C. Agar medium with lawn of *E.coli* was cut into pieces of about 1cm$^2$ in size and placed in a liquid PJM plates with nematode cultures. Futher strains of *Panagrolaimus n. sp.* are maintained on liquid PJM and Nematode Growth Medium (NGM) agar plates at +18°C[17].The stages of growing ancient nematodes from a frozen sample were repeated in the control lines. Laboratory bulk cultures and 10 strains (lineages) derived from isolates are reseeded every 2 months and maintained under laboratory conditions for more than 4 years until now. Morphological and phylogenetical investigations carried out used Pn2-1 strain.

**Light and scanning electron microscopy**

Nematodes were taken from a long-term culture *Pn2-1* strain. The specimens were collected from NGM plates and fixed in 4% formalin solution for about four hours and after that were picked into a Syracuse glass filled with mixture of glycerin, 95% ethanol, and distilled water (1:29:70). The glass were placed in a drying oven set to 40°C for 1–2 days until completely dehydrated, as in the glycerin-ethanol method[18]. Specimens were identified, measured, photographed, and drawn under a Leica DM5000 light microscope equipped with Leica Application Suite Version 3.8.0 software and a Leica DFC 425 C digital camera. All the measured sizes are given in µm. For scanning electron microscopy, formalin-fixed specimens were dehydrated in a series: 40% ethanol, 70% ethanol, 95% ethanol, 95% ethanol + acetone (50:50), acetone I, and acetone II, and then critical point dried. Once dried, specimens were mounted on a stub to be coated with platinum–palladium alloy and examined with Cam Scan S-2.

**Phylogenomic Analyses**

To explore the phylogenetic position of the novel species 18S and 28S rRNA gene sequences from 43 taxa across the *Propanagrolaimus*, *Panagrolaimus*, *Panagrellus* and *Halicephalobus* genera were downloaded from Genbank[19], with sequences from *Caenorhabditis briggsae* used as an outgroup (Supplementary Table 2). Both genes were identified in *Halicephalobus mephisto* (BioProject PRJNA528747) using BLAST[20] and a closely related reference (*Halicephalobus sp. LD6*). Nucleotide sequence alignments were generated using MAFFT L-INS-I[21] (v7.475). Multiple sequence alignments were manually curated and concatenated into a supermatrix[22], with a species tree inferred using maximum likelihood via IQTREE[23]. This supermatrix was partitioned using best-fit model of sequence evolution per gene inferred using ModelFinder[24] with 1000 bootstrap pseudo replicates used to estimate nodal support.

To further expand on the phylogenetic representation, 60 genes present in the novel species that had representative clusters shared with *Brugia malayi*, *Caenorhabditis elegans*, *Caenorhabditis remanei*, *Loa loa*, *Necator americanus* and *Trichinella spiralis* on OrthoDB[25] ( minimum of 1 gene per species for the cluster to have been considered, minimum of 10 genes per species needed for inclusion) were used for phylogenetic analysis (Supplementary Table 4). Genome assemblies for an additional 88 nematode species and an outgroup platyhelminth (*Macrostomum lignano*) were downloaded from the WormBase Parasite[26] database (Supplementary Table 5). Additional assemblies the *Panagrolaimus* genera[27], *Panagrolaimus superbus* (BioProject: PRJEB32708), *Panagrolaimus* sp. DL0137 (BioSample: SAMN06329917), *Panagrolaimus* sp. ES5 (BioSample: SAMN06329914), *Panagrolaimus* sp. PS1159 (BioSample: SAMN06329915), *Panagrolaimus* sp. PS1579 (BioSample: SAMN06329916) and the species *Propanagrolaimus* sp. JU765 (BioSample: SAMN06329918) were also included (n=100 nematode species). As these genomes were not present in the OrthoDB clusters, possible orthologs were identified by mapping *C. elegans* sequences for all genes to each genome, with a minimum threshold of 70% sequence identity and 75% gene coverage for a hit to be considered a putative ortholog. Given the potential for sequence quality negatively impacting sequence alignment, the frame-shift-aware alignment software MACSE[28] was used to generate multiple sequence alignments using the '*alignSequences*' function. The phylogeny was inferred using the supertree method with IQTREE as described above.

To estimate the rate of amino acid sequence diversity, the number of substitutions per site between the novel species, *Panagrolaimus* sp. PS1159 and *Panagrolaimus* sp. ES5, gene trees for each orthogroup were inferred using IQTREE. Distances between branches for all orthogroups were determined using the R package '*ape*'[29].
