## Supplementary material for "A novel nematode species from the Siberian permafrost shares adaptive mechanisms for cryptobiotic survival with *C. elegans* dauer larva": Orthology analysis

**Supplementary file – ORTHOLOGY_ANALYSIS**

All phylogenies were generated with IQtree2 with 1000 bootstraps (-bb option). The scale bar corresponds to 0.1 estimated amino acid substitutions per site.

*Anaplectus granulosus* – ANAGRA, *Caenorhabditis elegans* – CAEELE, *Diploscapter coronatus* – DIPCOR, *Diploscapter pachys* – DIPPAC, *Halicephalobus mephisto* – HALMEP, *Neocamacolaimus parasiticus* – NEOPAR, *Panagrellus redivivus* – PANRED, *Panagrolaimus davidi* – PANDAV, *Panagrolaimus kolymaensis* – HLNpanKol1, *Panagrolaimus* sp. ES5 – PANES5, *Panagrolaimus* sp. PS1159 – PANPS1159, *Panagrolaimus superbus* – PANSUP, *Plectus murrayi* – PLEMUR, *Plectus sambesii* – PLESAM, *Plectus* sp. (from Permafrost) – HLNpleKol1, *Pristionchus pacificus* – PRIPAC, *Propanagrolaimus* sp. JU765 – PROJU765, *Stephanolaimus elegans* – STEELE.

pp. 2-3: Trehalose synthesis genes

pp. 4-20: TCA cycle

p. 21: Glyoxylate shunt

pp. 22-38: Glycolysis / Gluconeogenesis

pp. 39-40: Polyamine synthesis

pp. 41-62: Dauer genes

**Trehalose synthesis**

TPS-1 / TPS-2

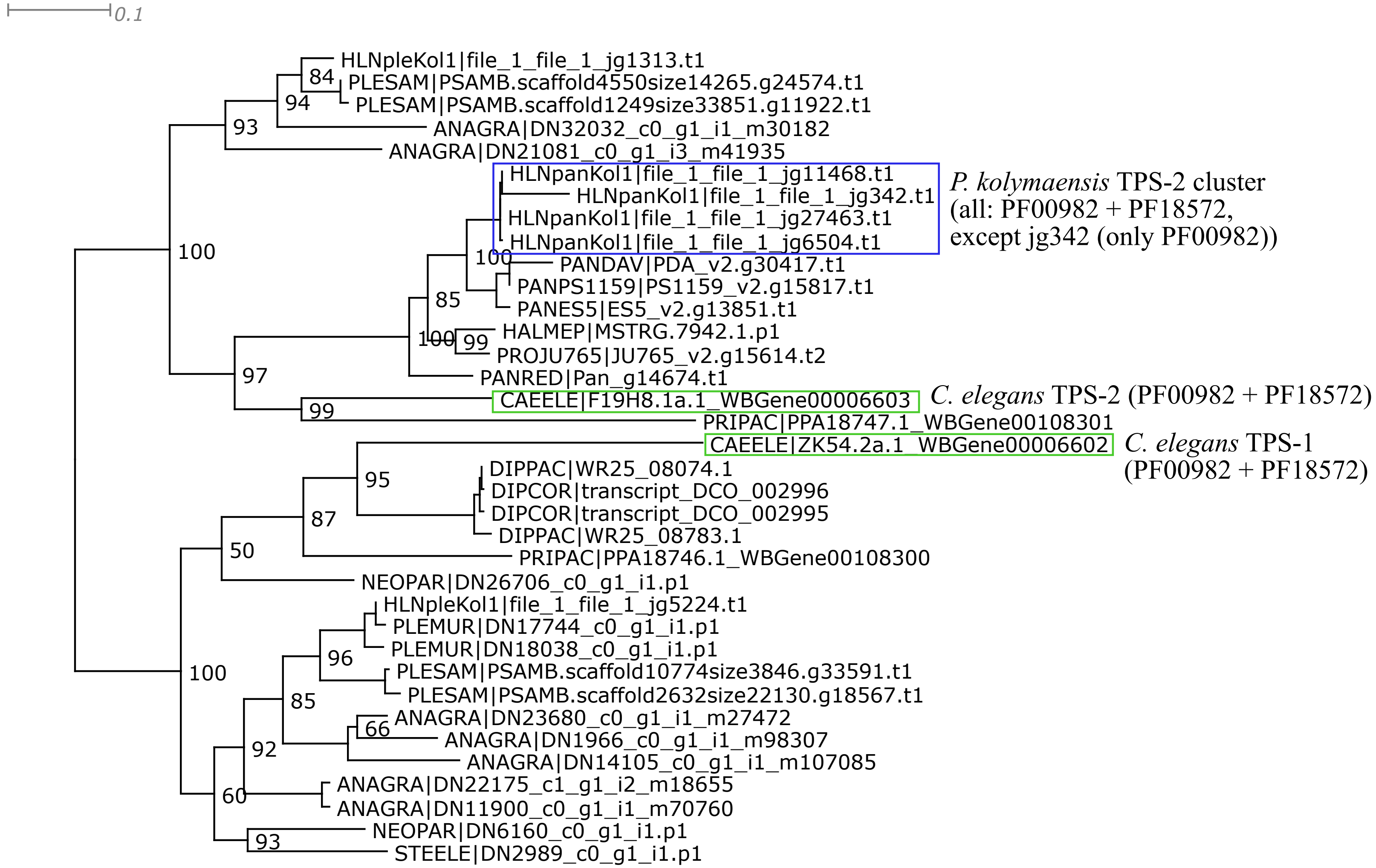


Trimal -automated1 function; one long-branched sequence (*D. pachys*) manually removed afterwards;
IQtree2 ML phylogeny best-fit model according to BIC: LG+I+G4
Almost all plectid sequences that cluster with *C. elegans* TPS-1 do have large gaps in the alignment and were removed at first;
Then the clustering of the one remaining plectid sequence not so clear, therefore, the short sequences were left in the phylogeny.
Panagrolaimids appear to encode only TPS-2.

GOB-1

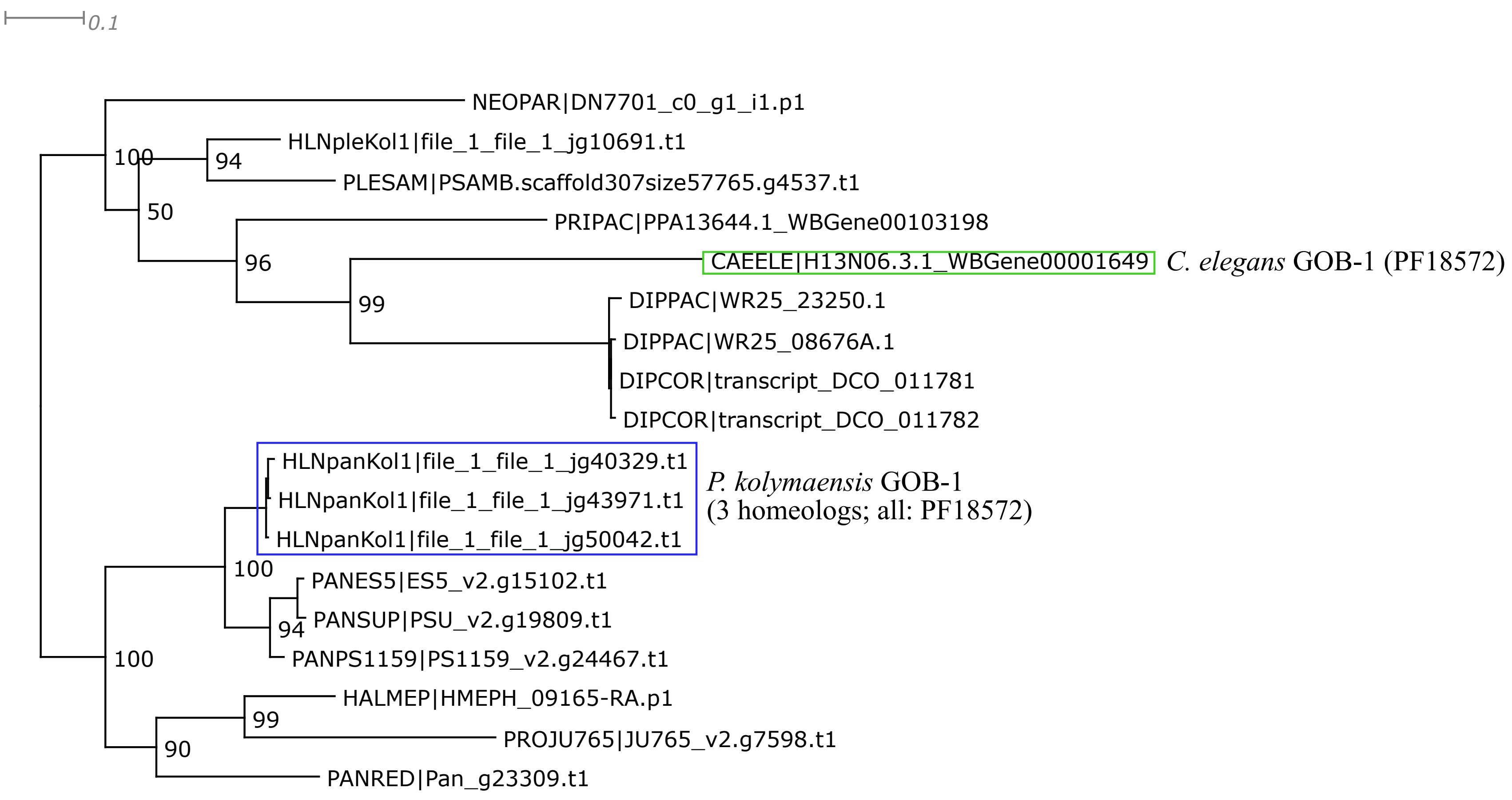


Trimal -automated1 function; short or spurious sequences manually removed afterwards;
IQtree2 ML phylogeny best-fit model according to BIC: LG+G4

**TCA Cycle**

CTS-1

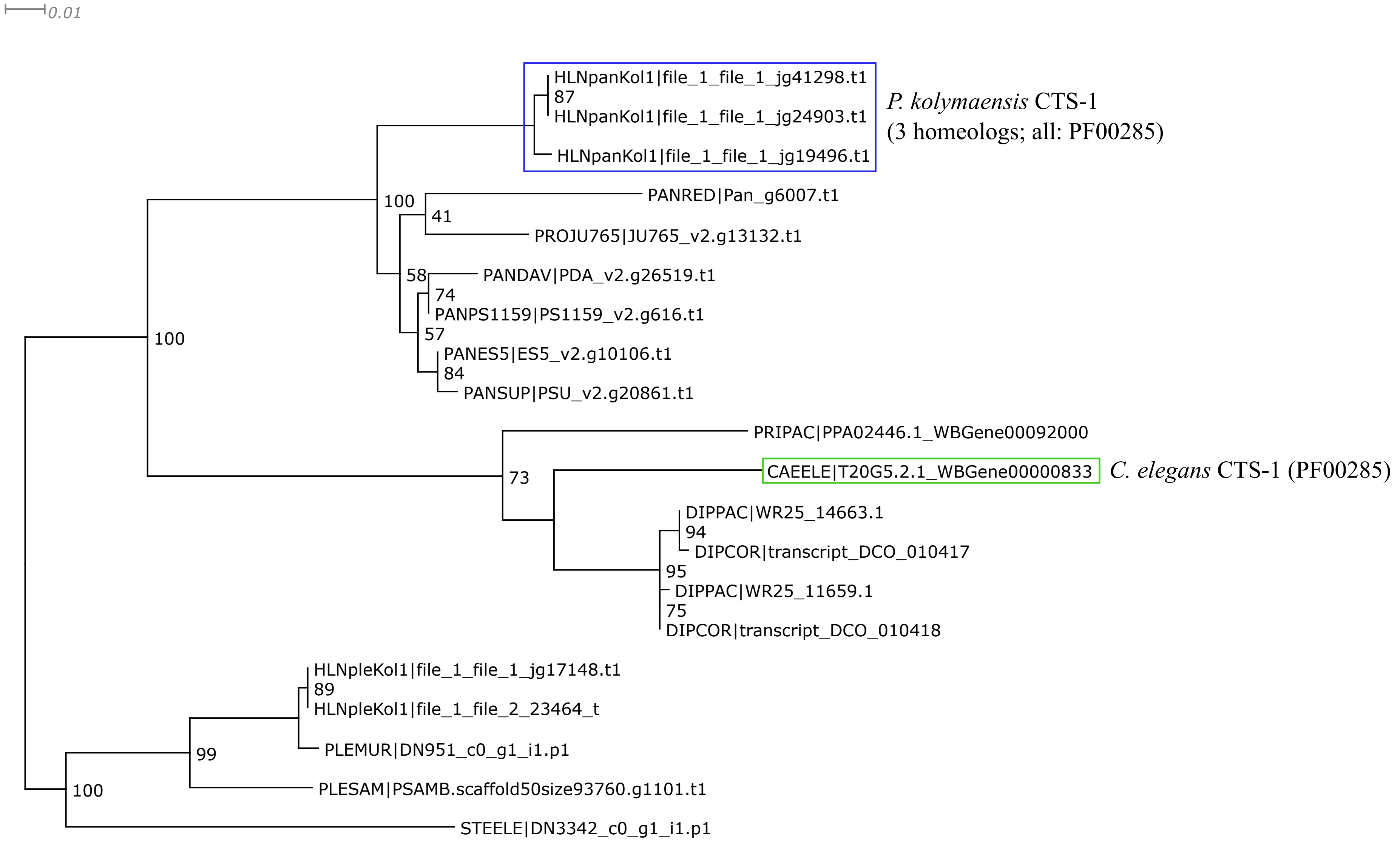


Trimal -automated1 function; short or spurious sequences manually removed afterwards;
IQtree2 ML phylogeny best-fit model according to BIC: LG+G4

ACO-1

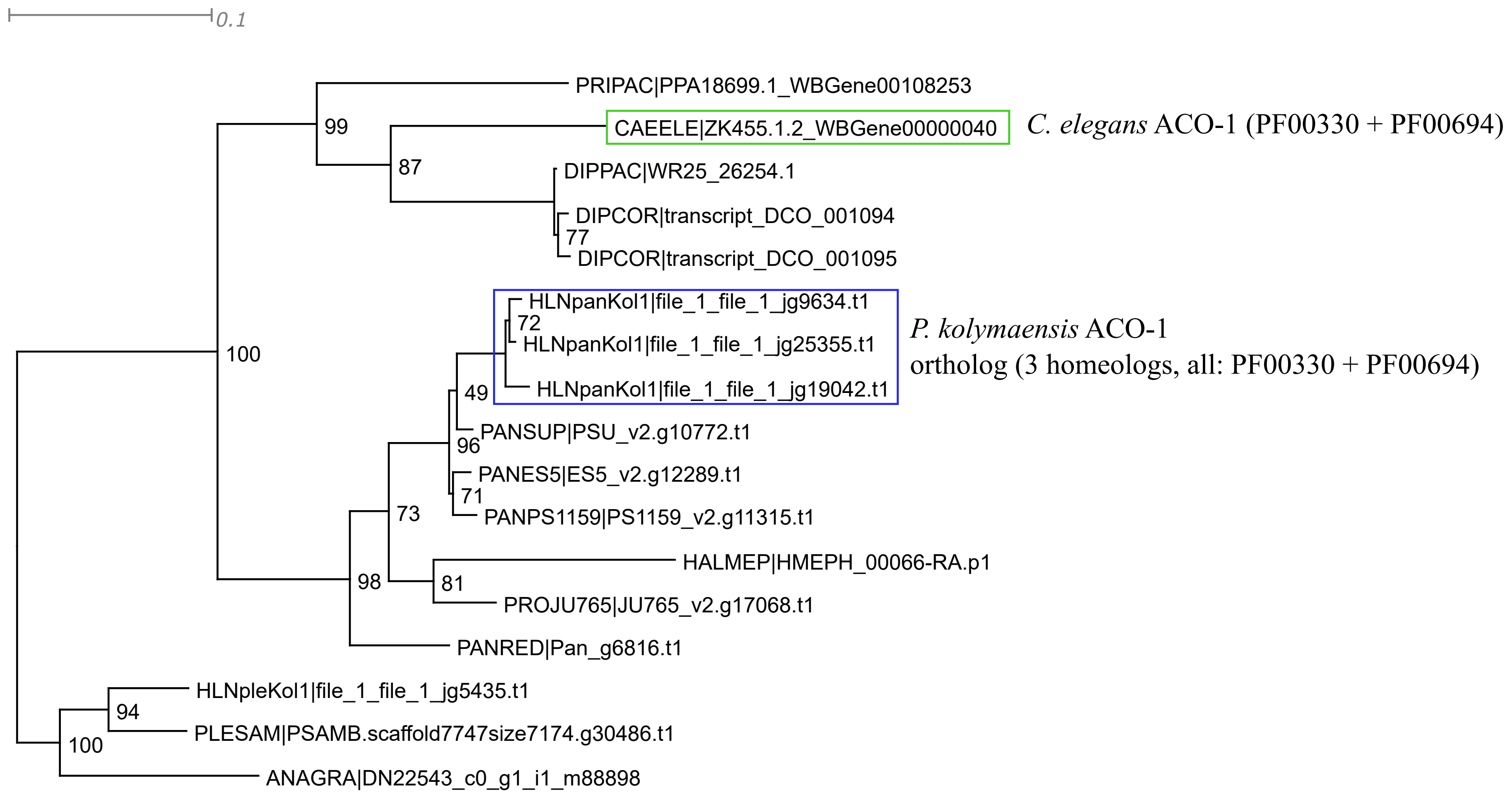


Trimal -automated1 function; short or spurious sequences manually removed afterwards;
IQtree2 ML phylogeny best-fit model according to BIC: LG+I+G4

ACO-2

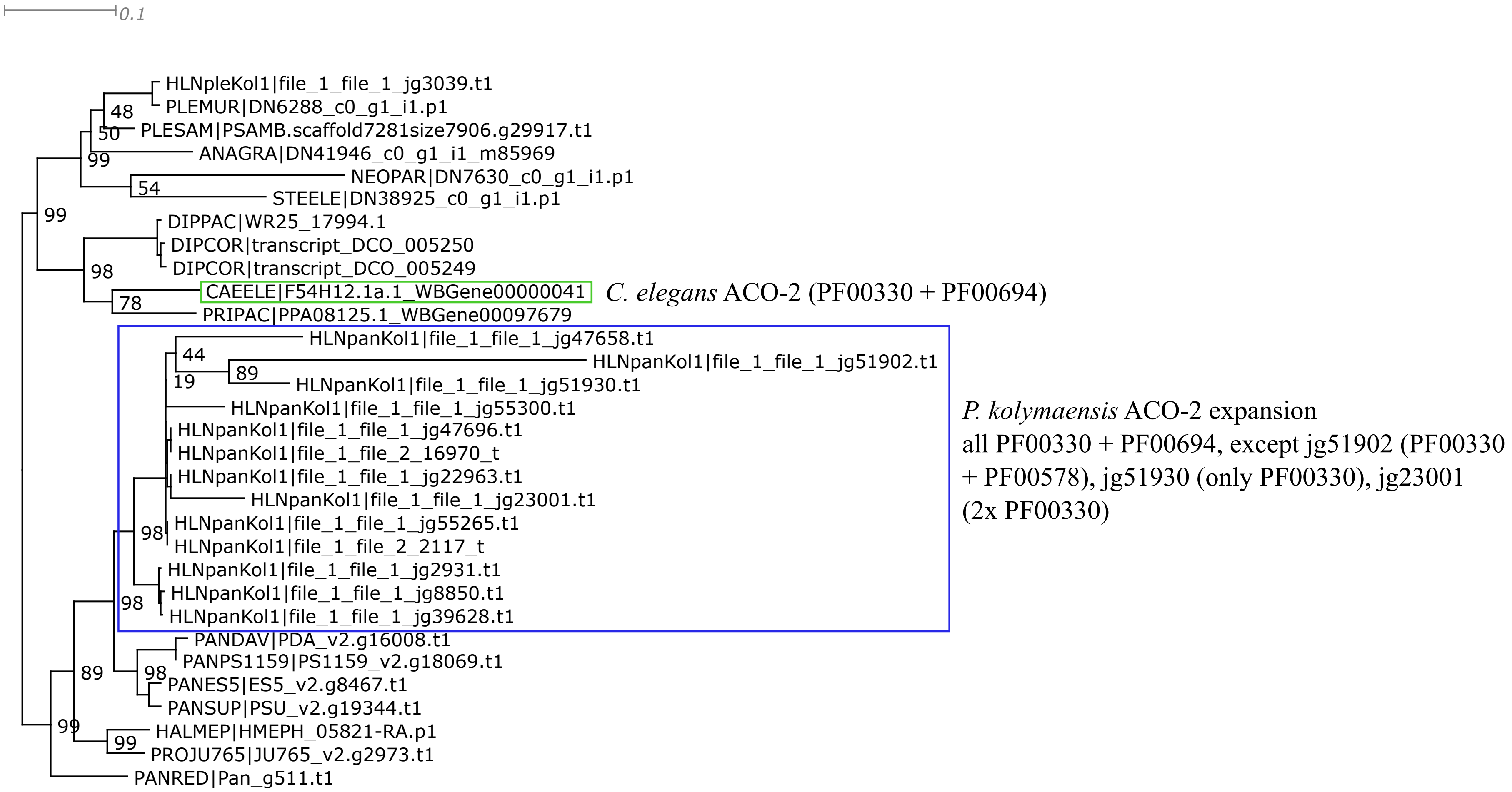


Trimal -automated1 function; short or spurious sequences manually removed afterwards;
IQtree2 ML phylogeny best-fit model according to BIC: WAG+G4

IDH-1 / IDH-2

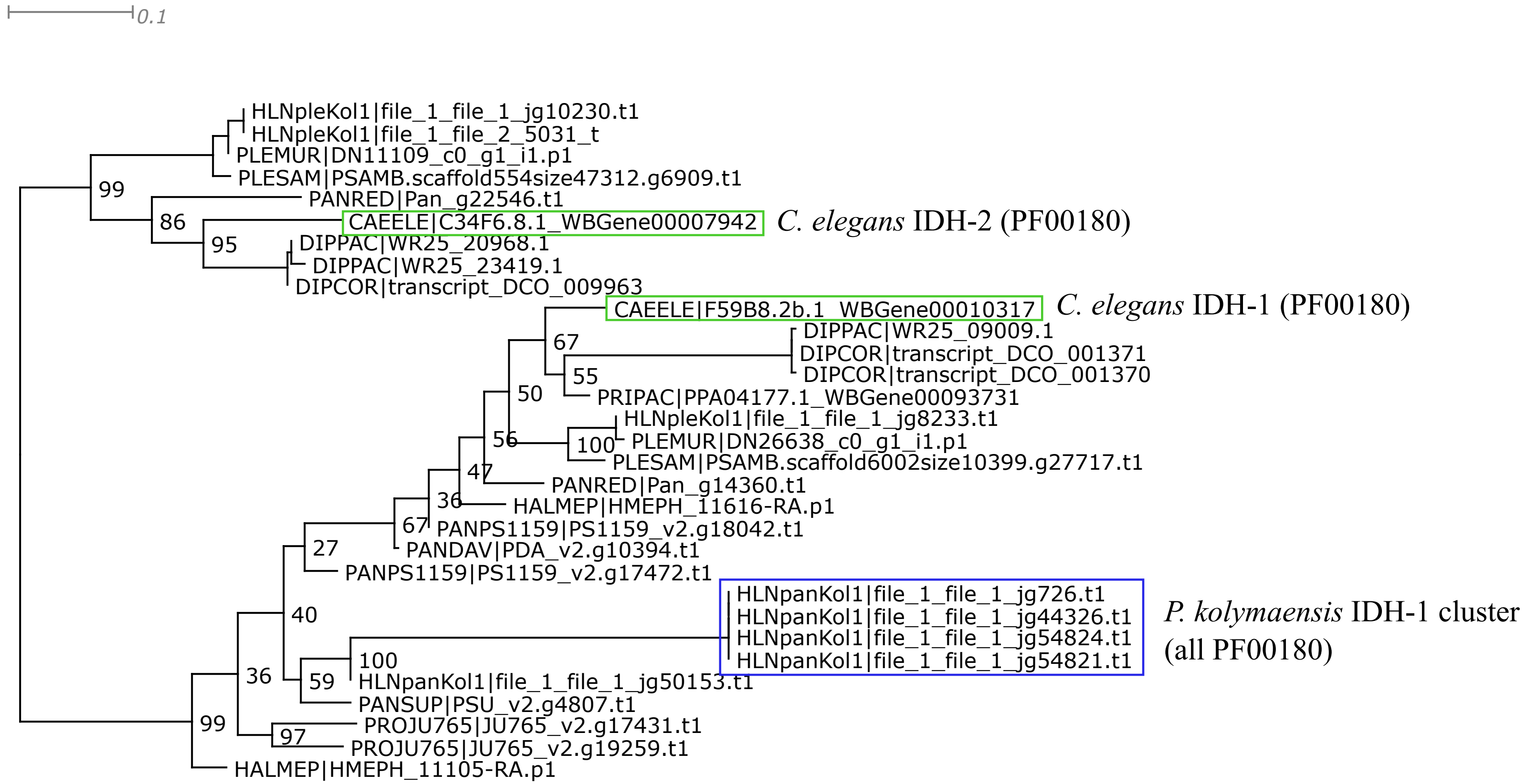


Trimal -automated1 function; short or spurious sequences manually removed afterwards;
IQtree2 ML phylogeny best-fit model according to BIC: WAG+I+G4

IDHA-1

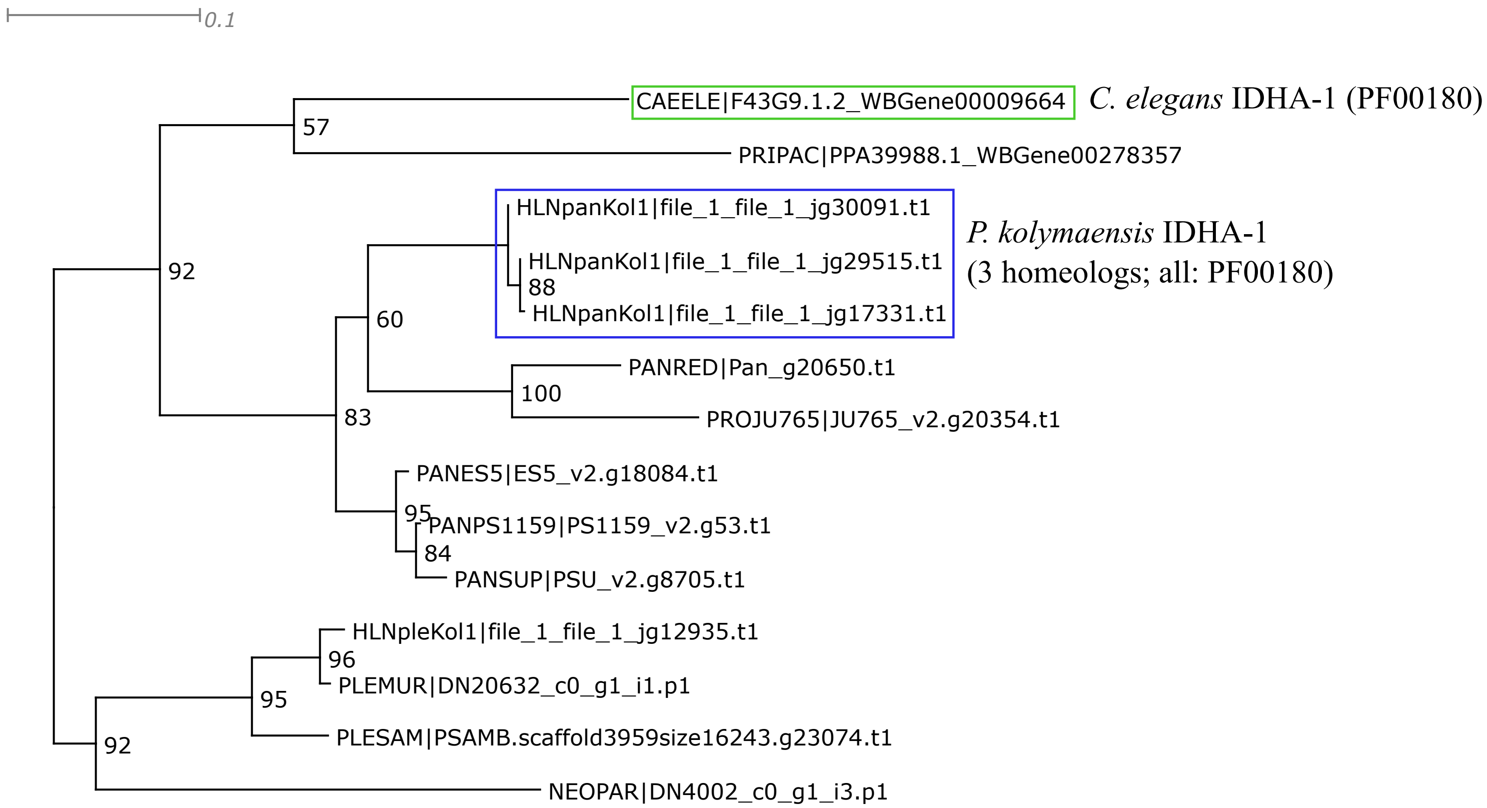


Trimal: 1. -resoverlap 0.75 -seqoverlap 80 functions; 2. -automated1 function; 3. short or spurious sequences manually removed afterwards; IQtree2 ML phylogeny best-fit model according to BIC: LG+I+G4

IDHB-1

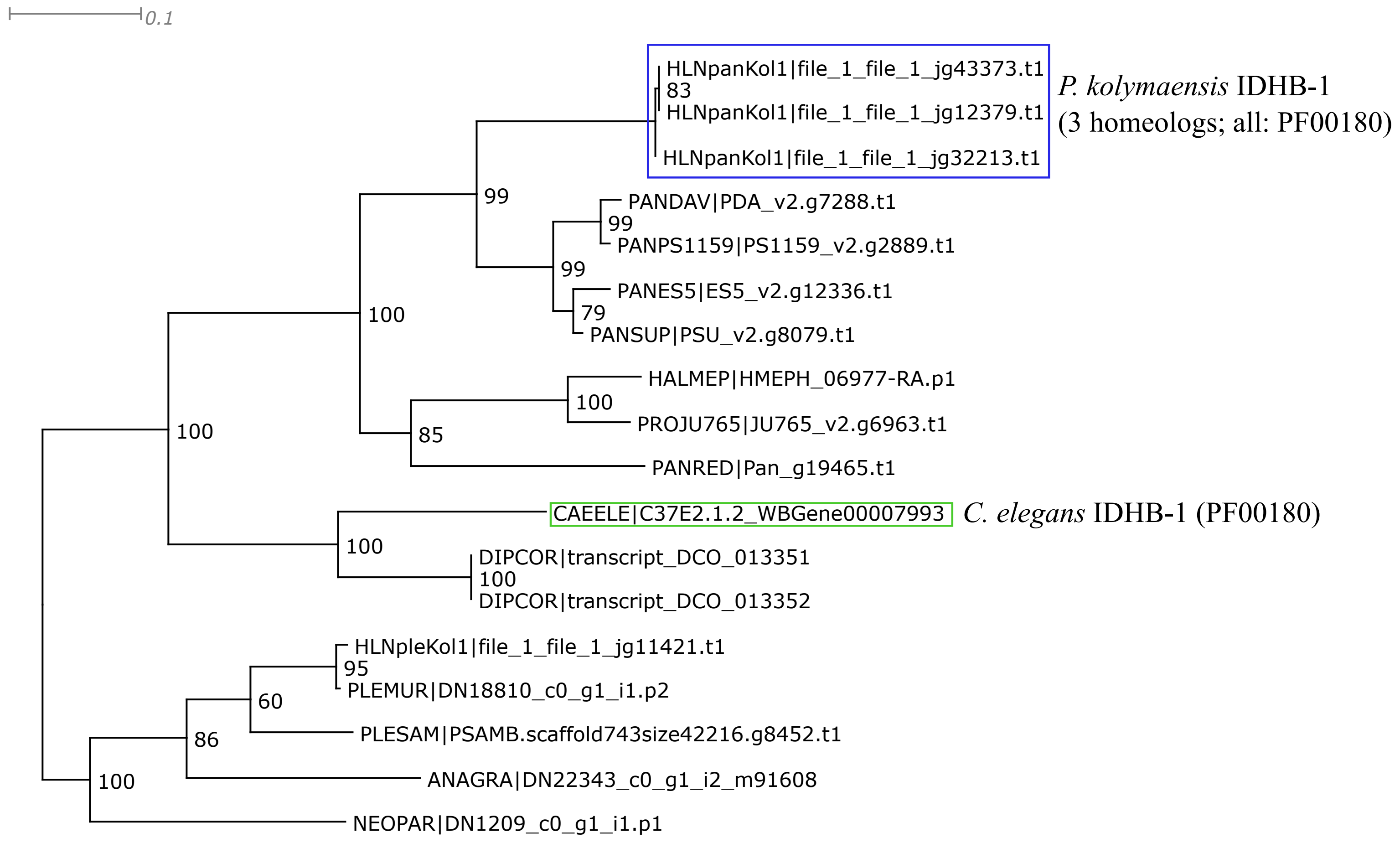


Trimal: 1. -resoverlap 0.75 -seqoverlap 80 functions; 2. -automated1 function; 3. short or spurious sequences manually removed afterwards; IQtree2 ML phylogeny best-fit model according to BIC: LG+I+G4

IDHG-1 / IDHG-2

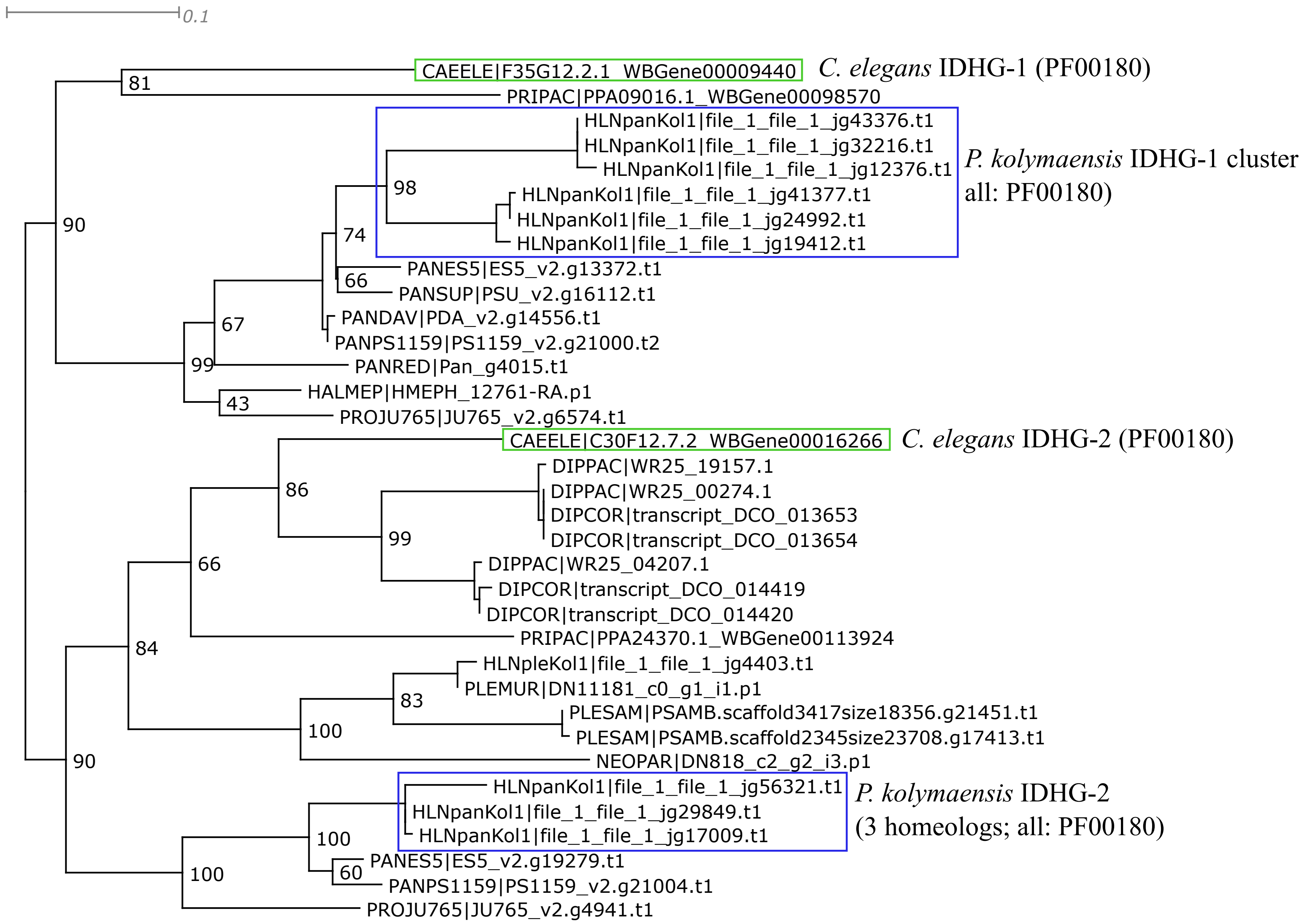


Trimal -automated1 function; short or spurious sequences manually removed afterwards;
IQtree2 ML phylogeny best-fit model according to BIC: LG+G4

MDH-1

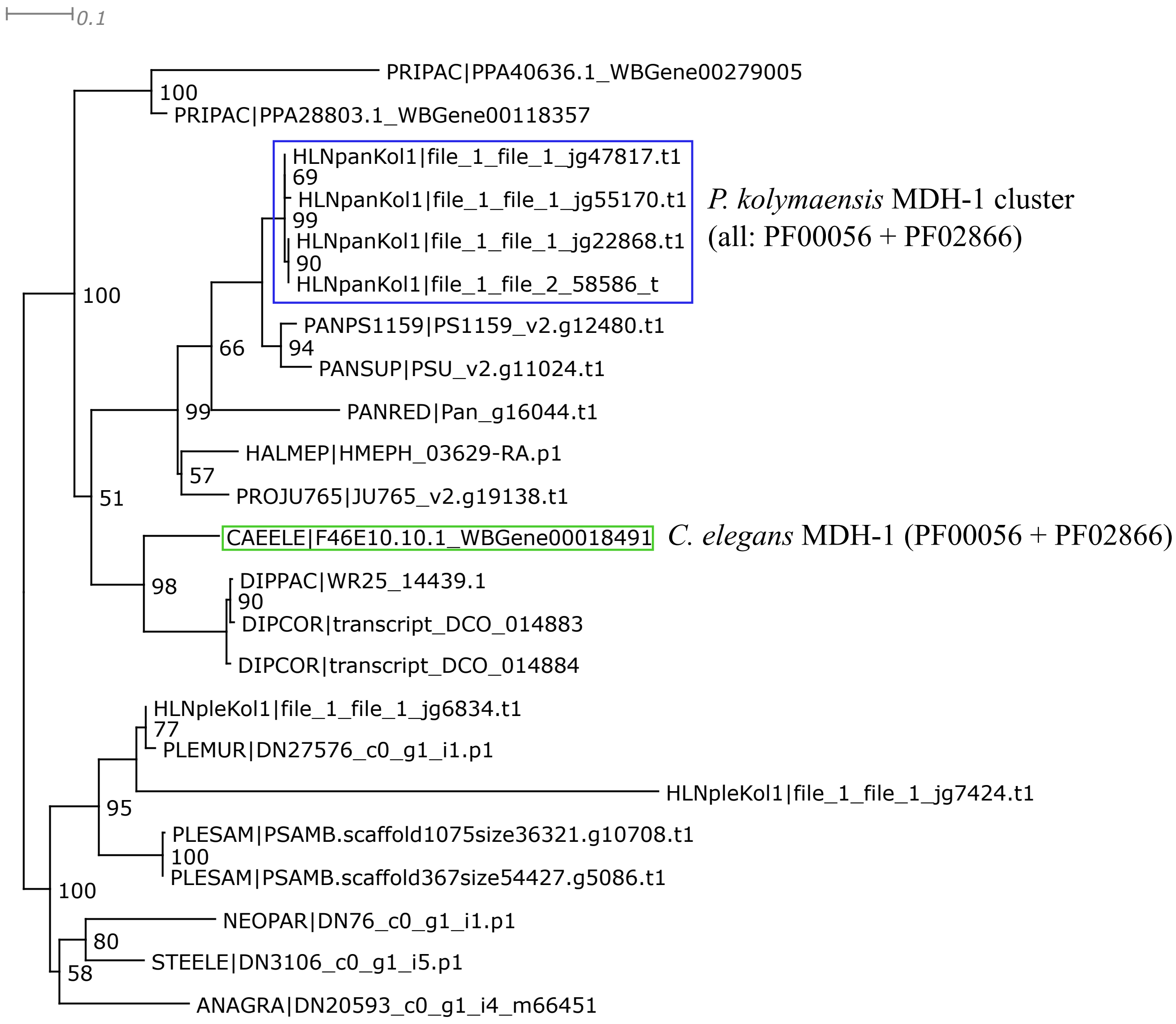


Trimal -automated1 function; short or spurious sequences manually removed afterwards;
IQtree2 ML phylogeny best-fit model according to BIC: LG+G4

MDH-2

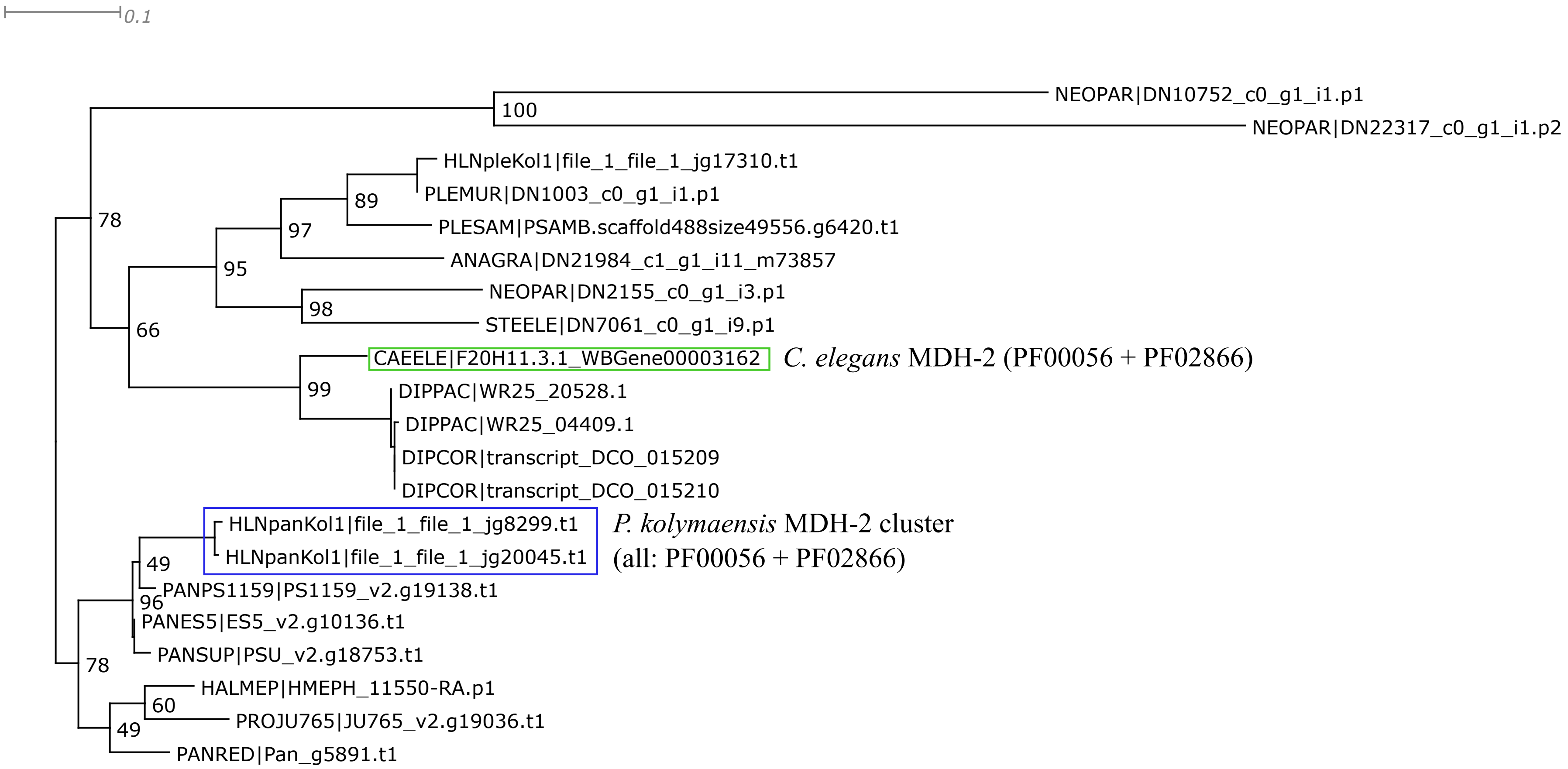


Trimal -automated1 function; short or spurious sequences manually removed afterwards;
IQtree2 ML phylogeny best-fit model according to BIC: LG+G4

OGDH-1 / OGDH-2

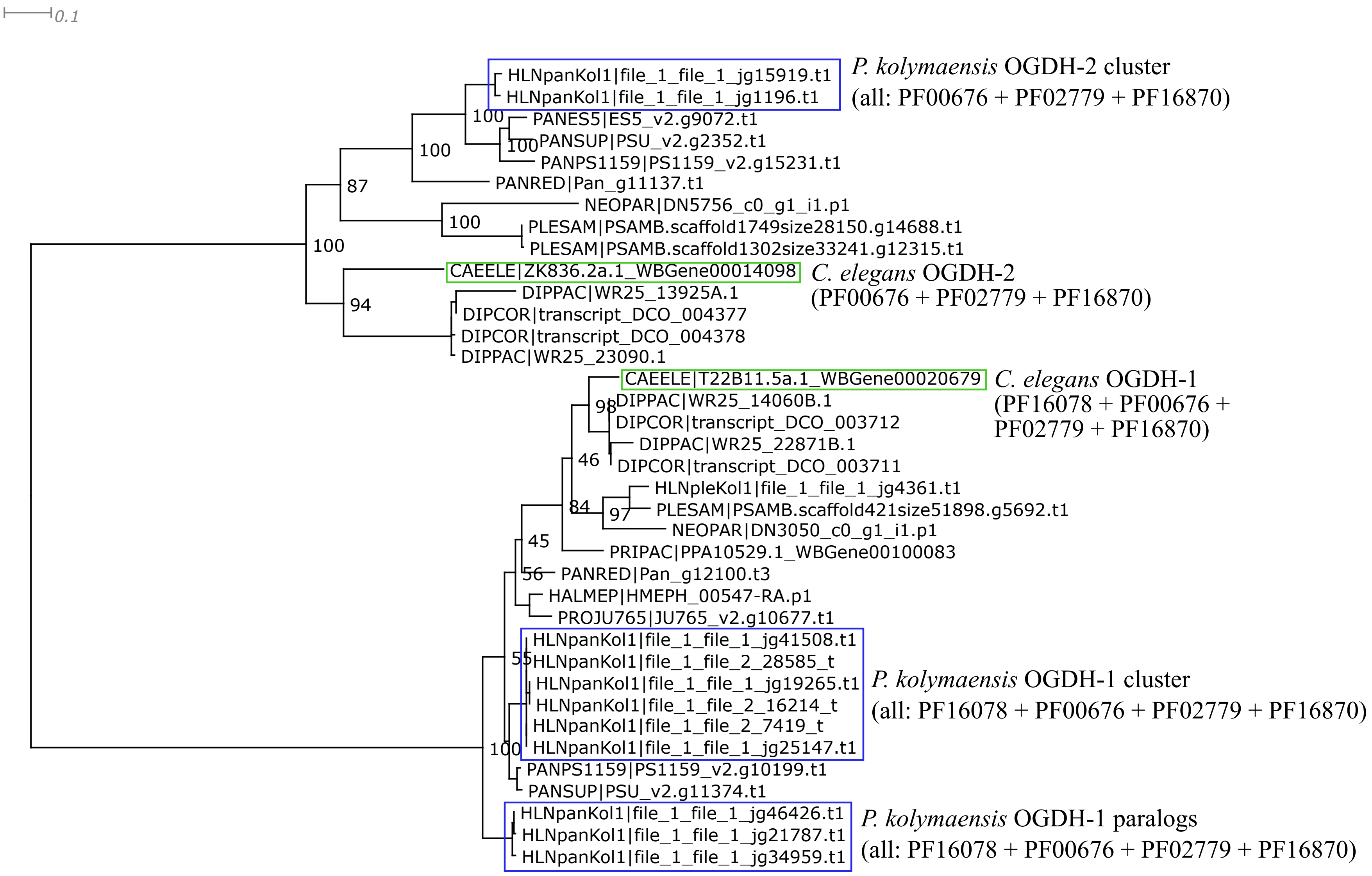


Trimal -automated1 function; short or spurious sequences manually removed afterwards;
IQtree2 ML phylogeny best-fit model according to BIC: LG+G4

SUCA-1

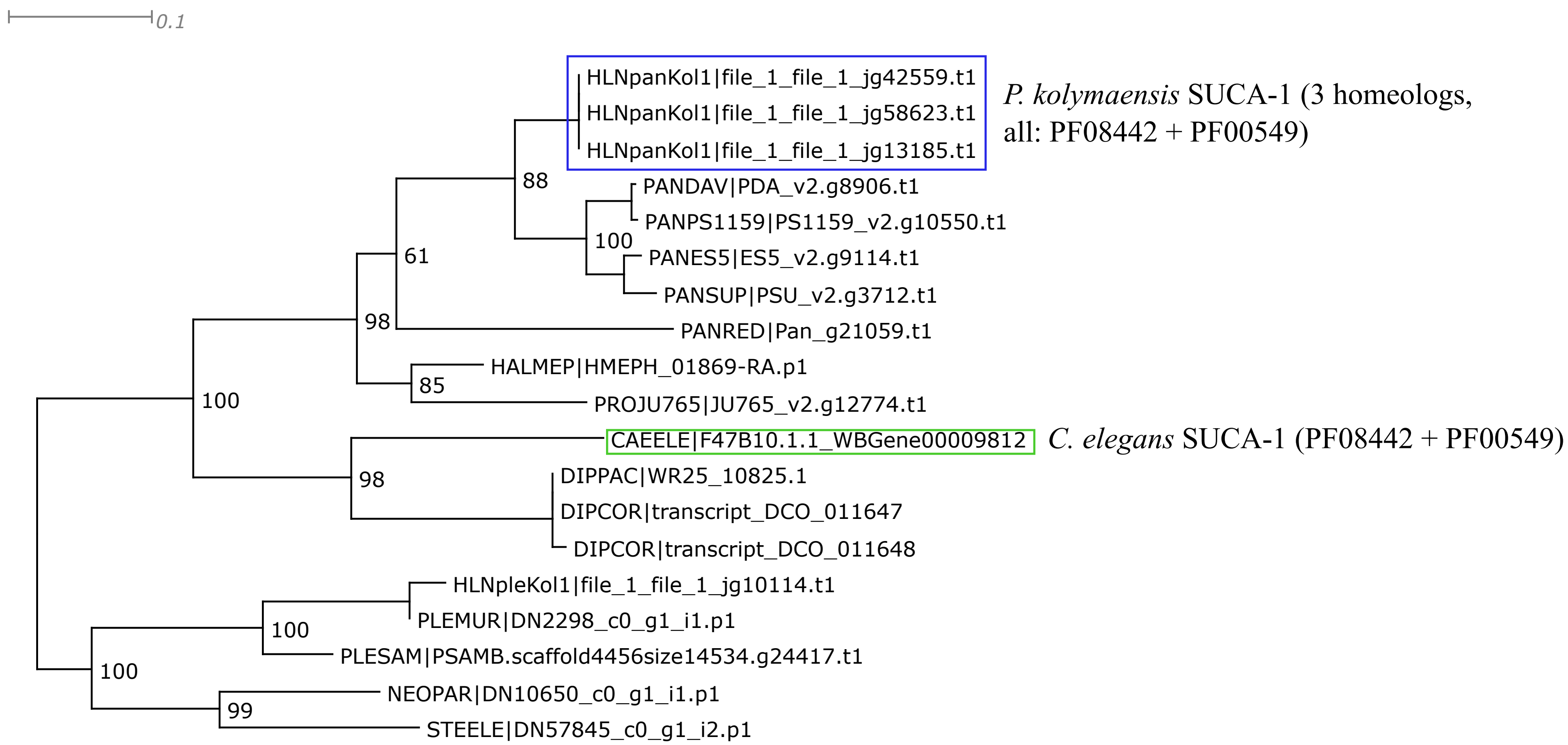


Trimal -automated1 function; short or spurious sequences manually removed afterwards;
IQtree2 ML phylogeny best-fit model according to BIC: LG+I+G4

SUCG-1

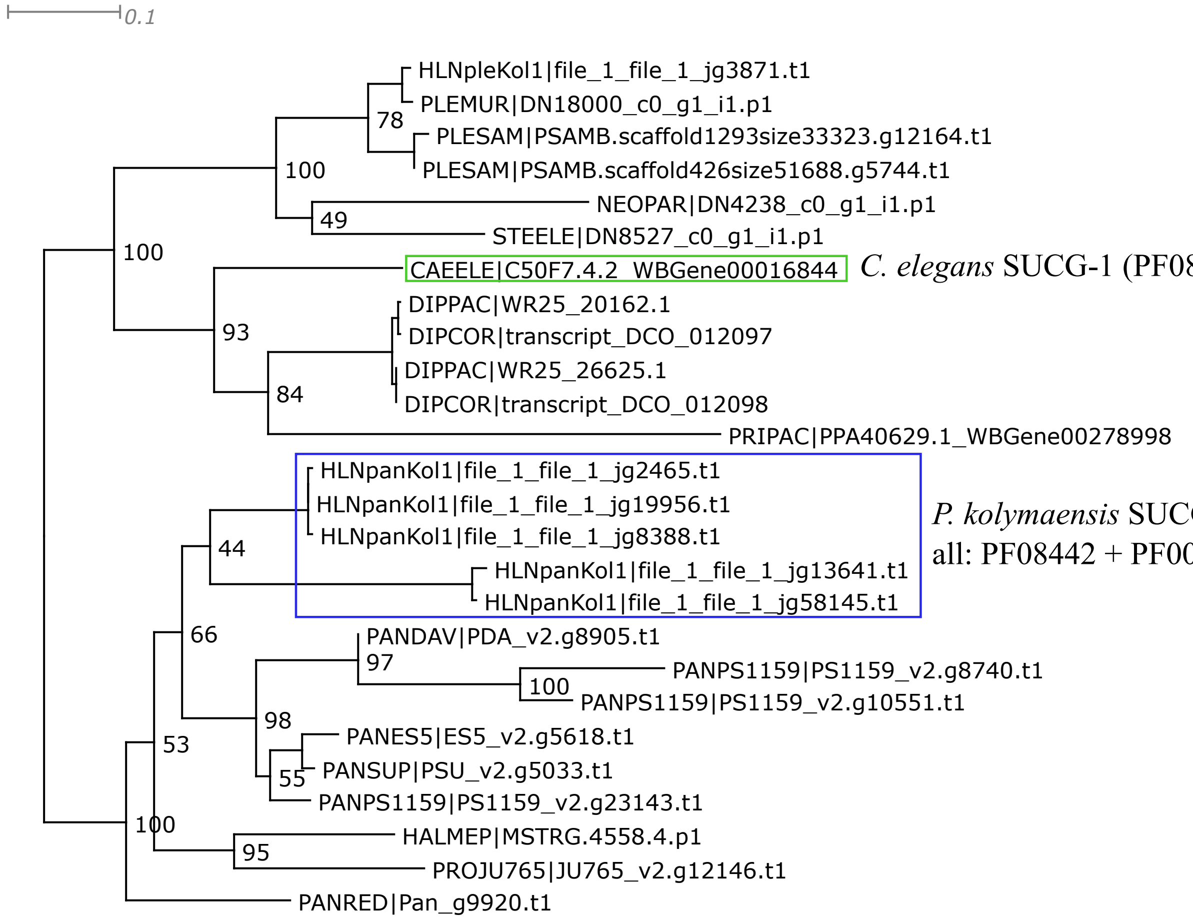


*C. elegans* SUCG-1 (PF08442 + PF00549)

*P. kolymaensis* SUCG-1 cluster; all: PF08442 + PF00549)

Trimal -automated1 function; short or spurious sequences manually removed afterwards;
IQtree2 ML phylogeny best-fit model according to BIC: LG+I+G4

SUCL-1 / SUCL-2

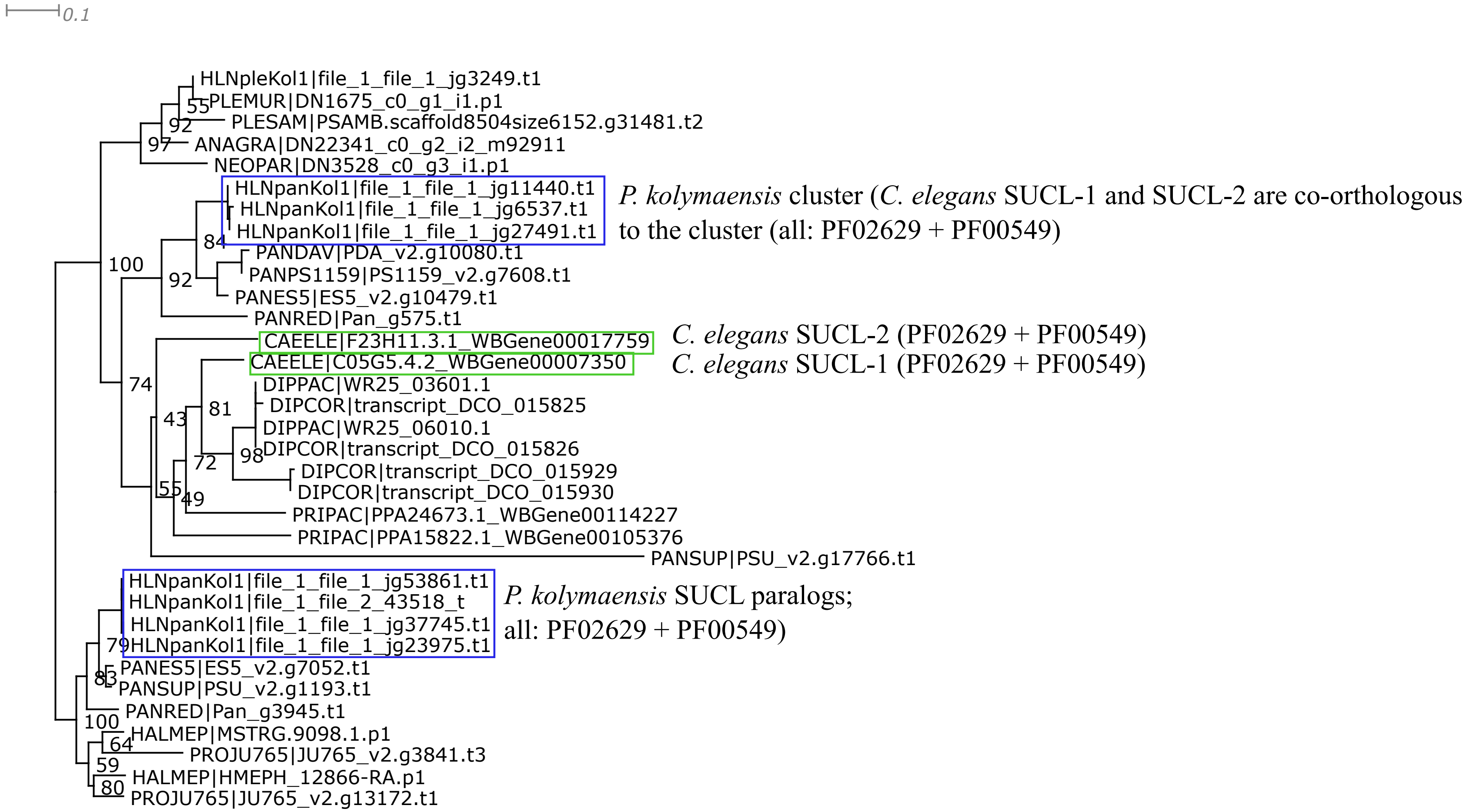


Trimal -automated1 function; short or spurious sequences manually removed afterwards;
IQtree2 ML phylogeny best-fit model according to BIC: LG+G4

SDHA-1 / SDHA-2

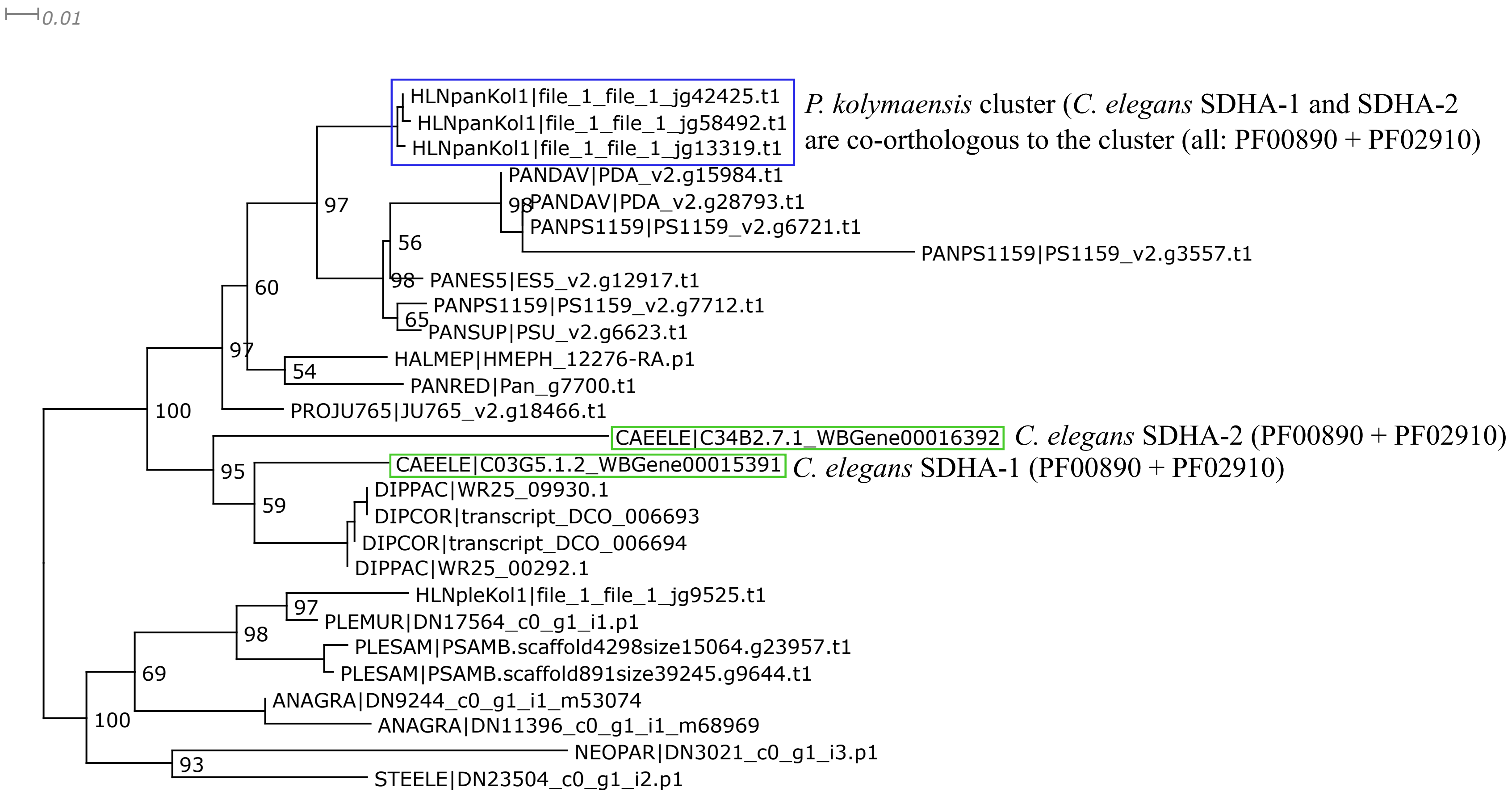


Trimal: 1. -resoverlap 0.75 -seqoverlap 75 functions; 2. -automated1 function; 3. short or spurious sequences manually removed afterwards; IQtree2 ML phylogeny best-fit model according to BIC: LG+I+G4

SDHB-1

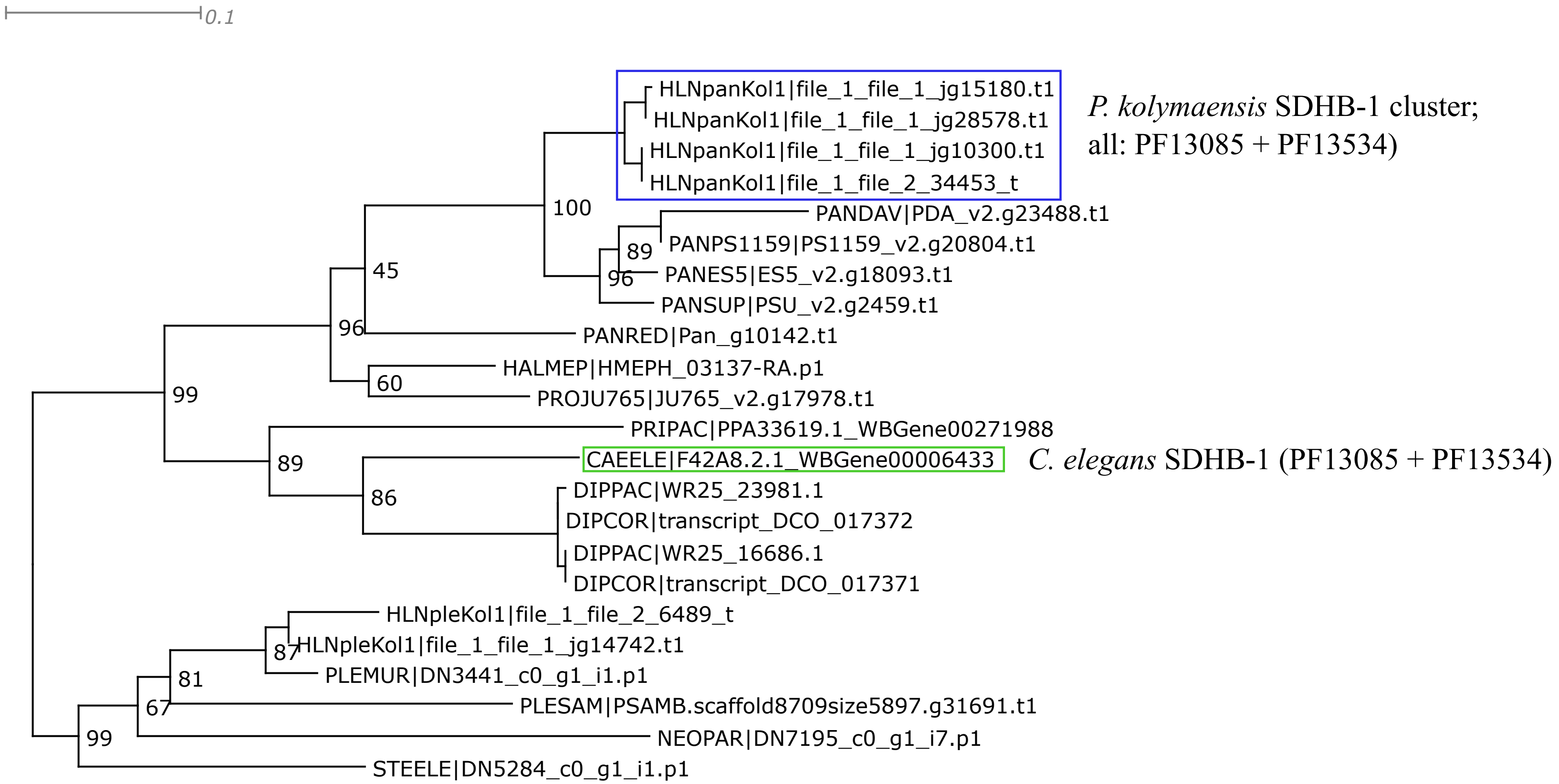


Trimal -automated1 function; short or spurious sequences manually removed afterwards;
IQtree2 ML phylogeny best-fit model according to BIC: WAG+I+G4

MEV-1

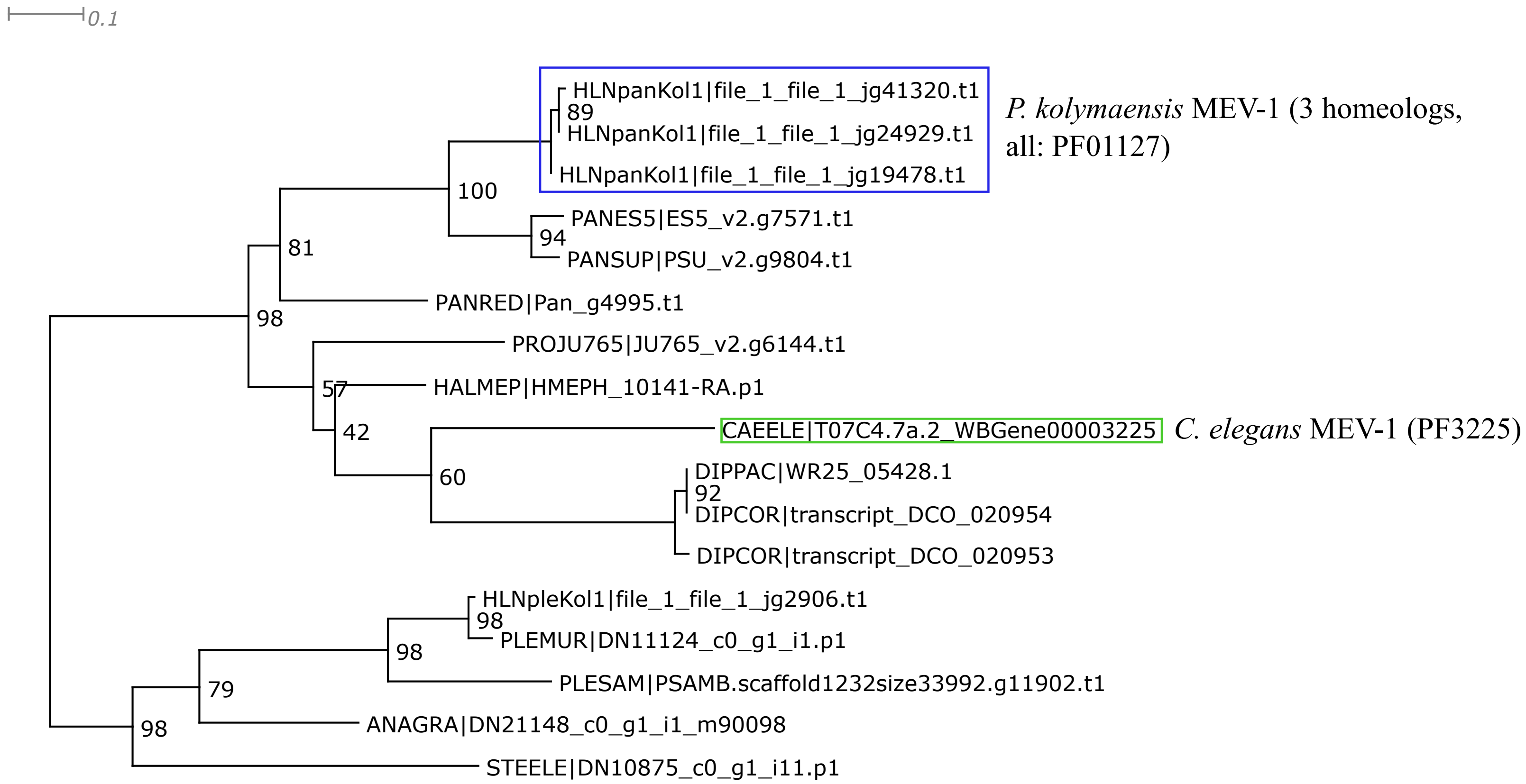


Trimal -automated1 function; short or spurious sequences manually removed afterwards;
IQtree2 ML phylogeny best-fit model according to BIC: LG+G4

FUM-1

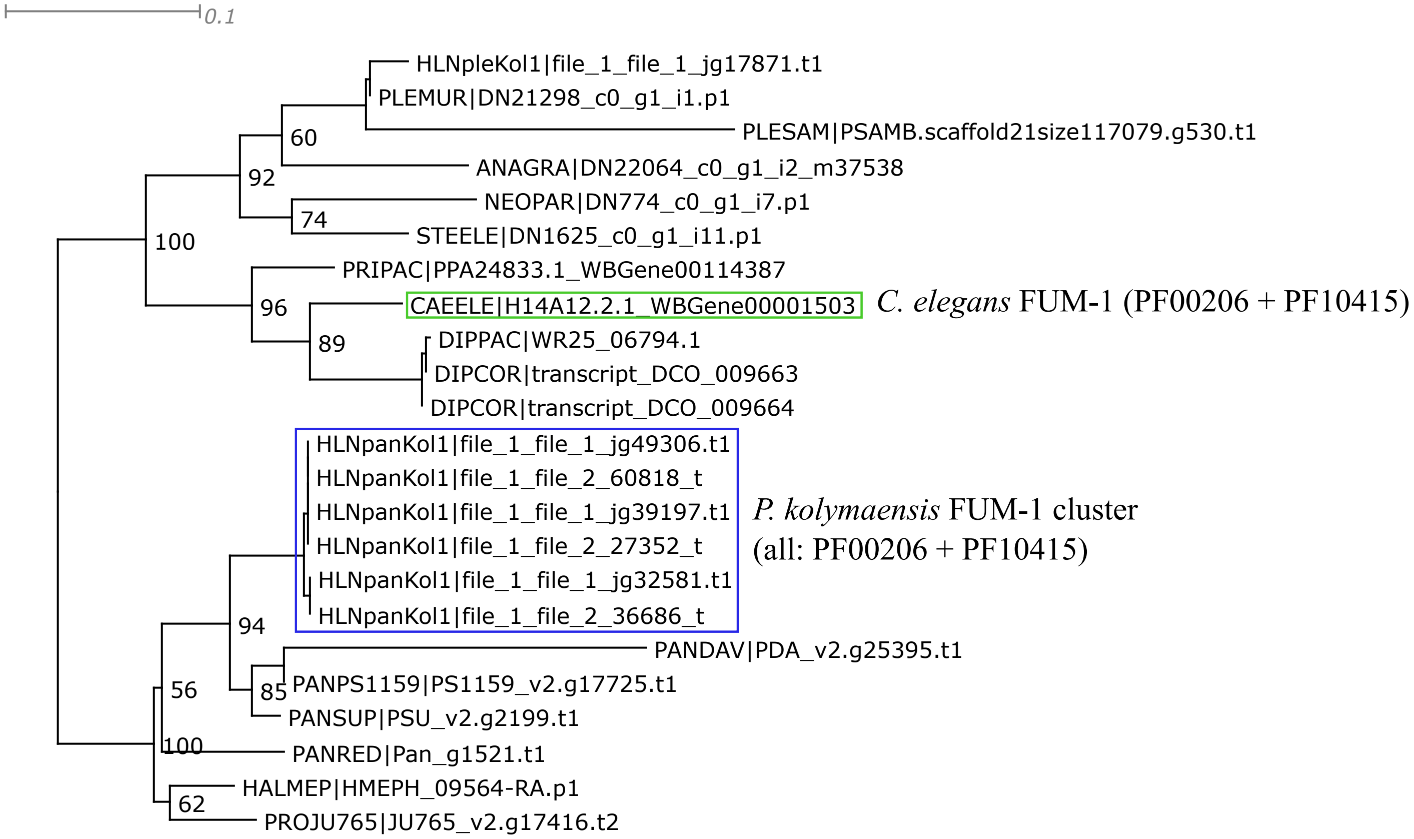


Trimal -automated1 function; short or spurious sequences manually removed afterwards;
IQtree2 ML phylogeny best-fit model according to BIC: WAG+G4

**Glyoxylate shunt**

ICL-1

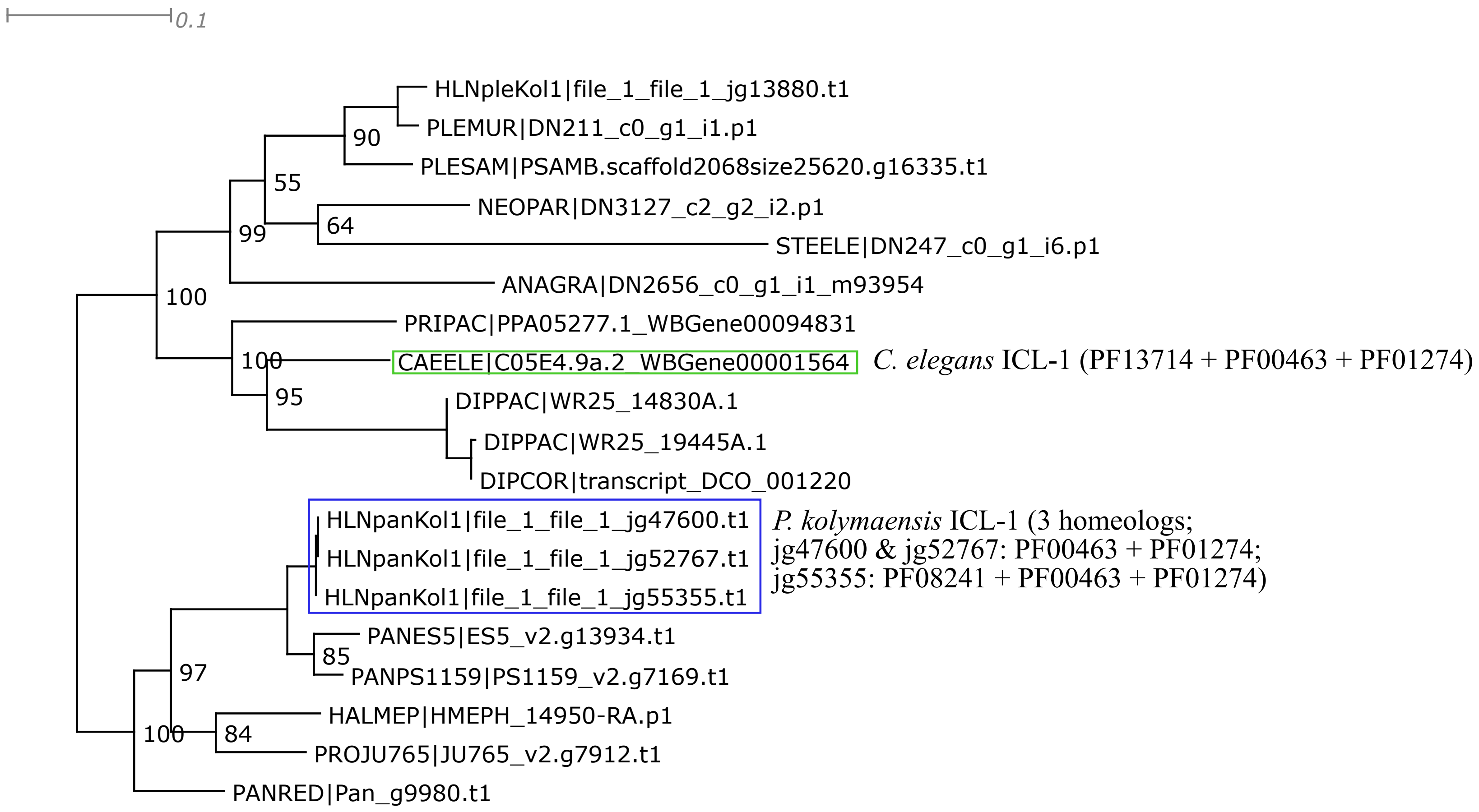


Trimal -automated1 function; short or spurious sequences manually removed afterwards;
IQtree2 ML phylogeny best-fit model according to BIC: LG+G4
Only sequences included that contain both Isocitrate lyase and Malate synthase combined.

**Glycolysis / Gluconeogenesis**

PDHB-1

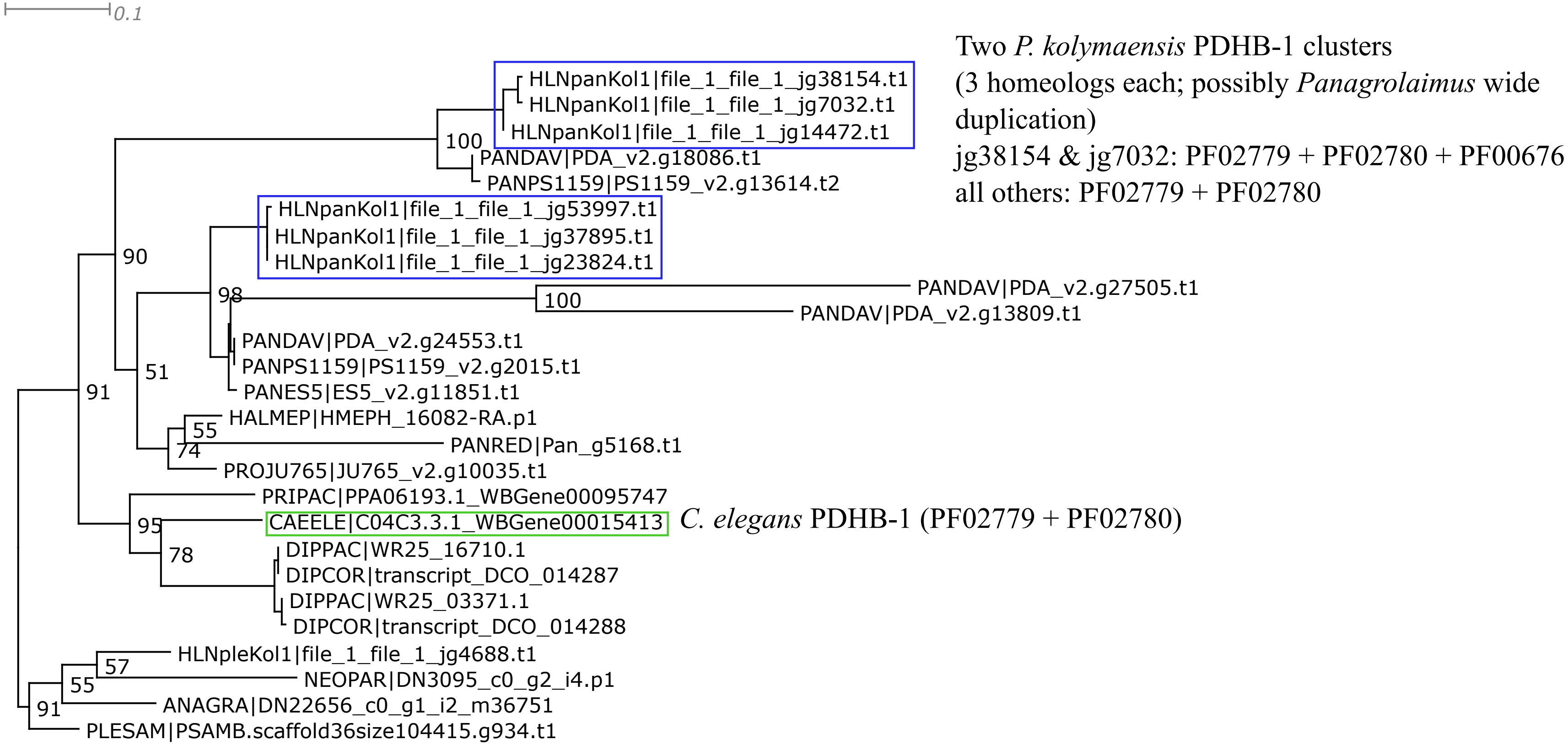


Trimal -automated1 function; short or spurious sequences manually removed afterwards;
IQtree2 ML phylogeny best-fit model according to BIC: LG+G4

PDHA-1

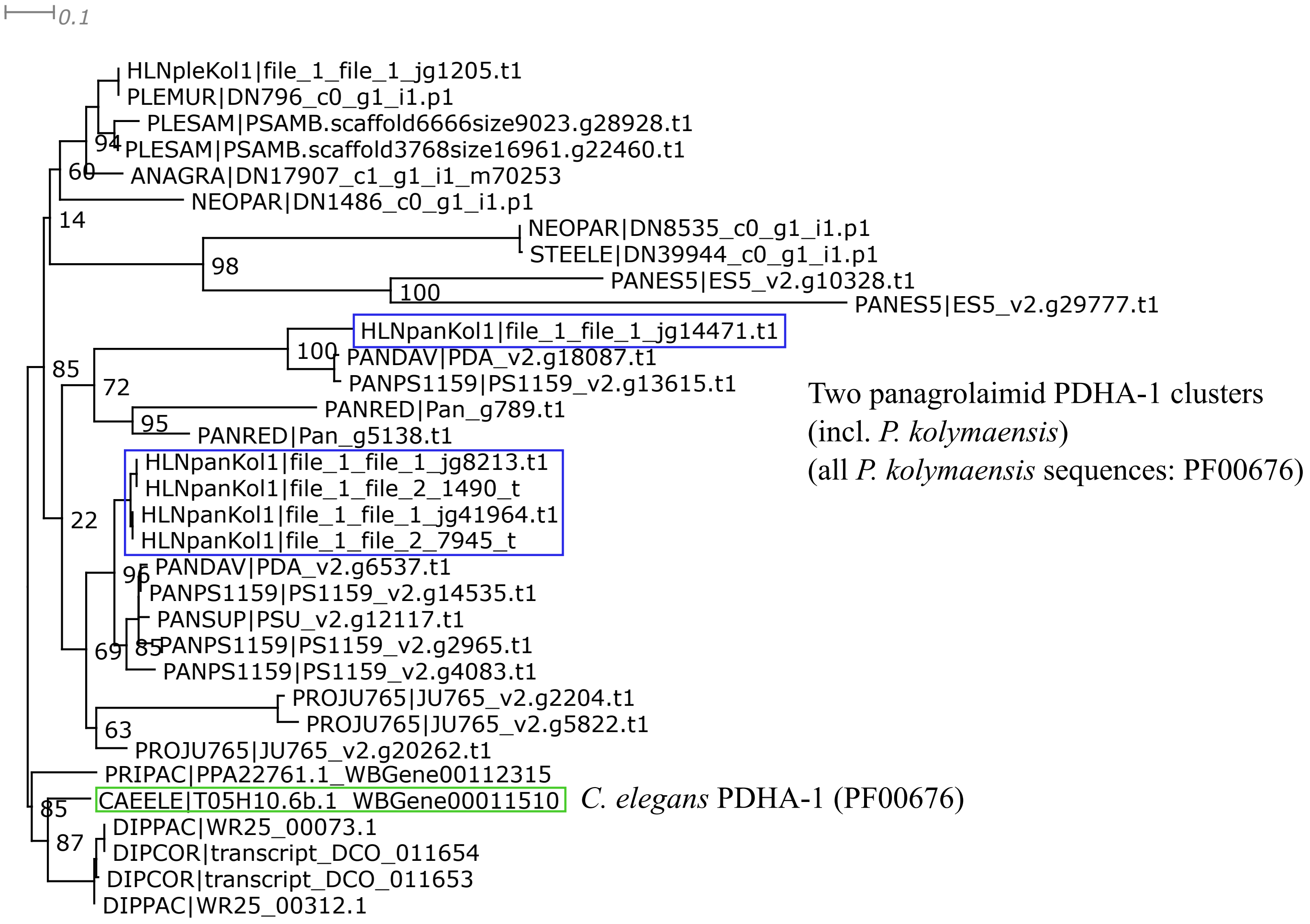


Trimal -automated1 function; short or spurious sequences manually removed afterwards;
IQtree2 ML phylogeny best-fit model according to BIC: LG+I+G4

DLAT-1 / DLAT-2

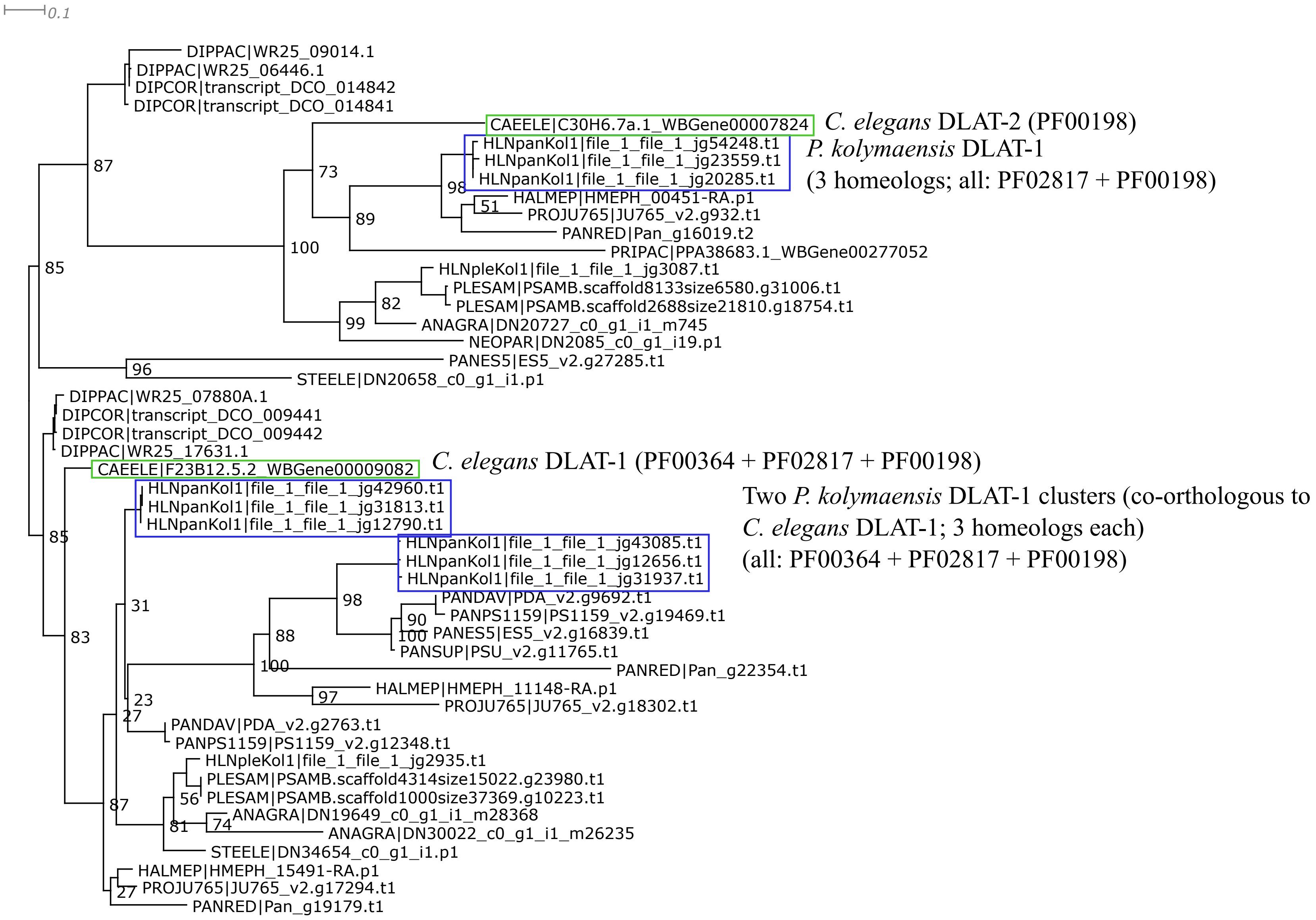


Trimal -automated1 function; short or spurious sequences manually removed afterwards;
IQtree2 ML phylogeny best-fit model according to BIC: LG+G4

DLD-1

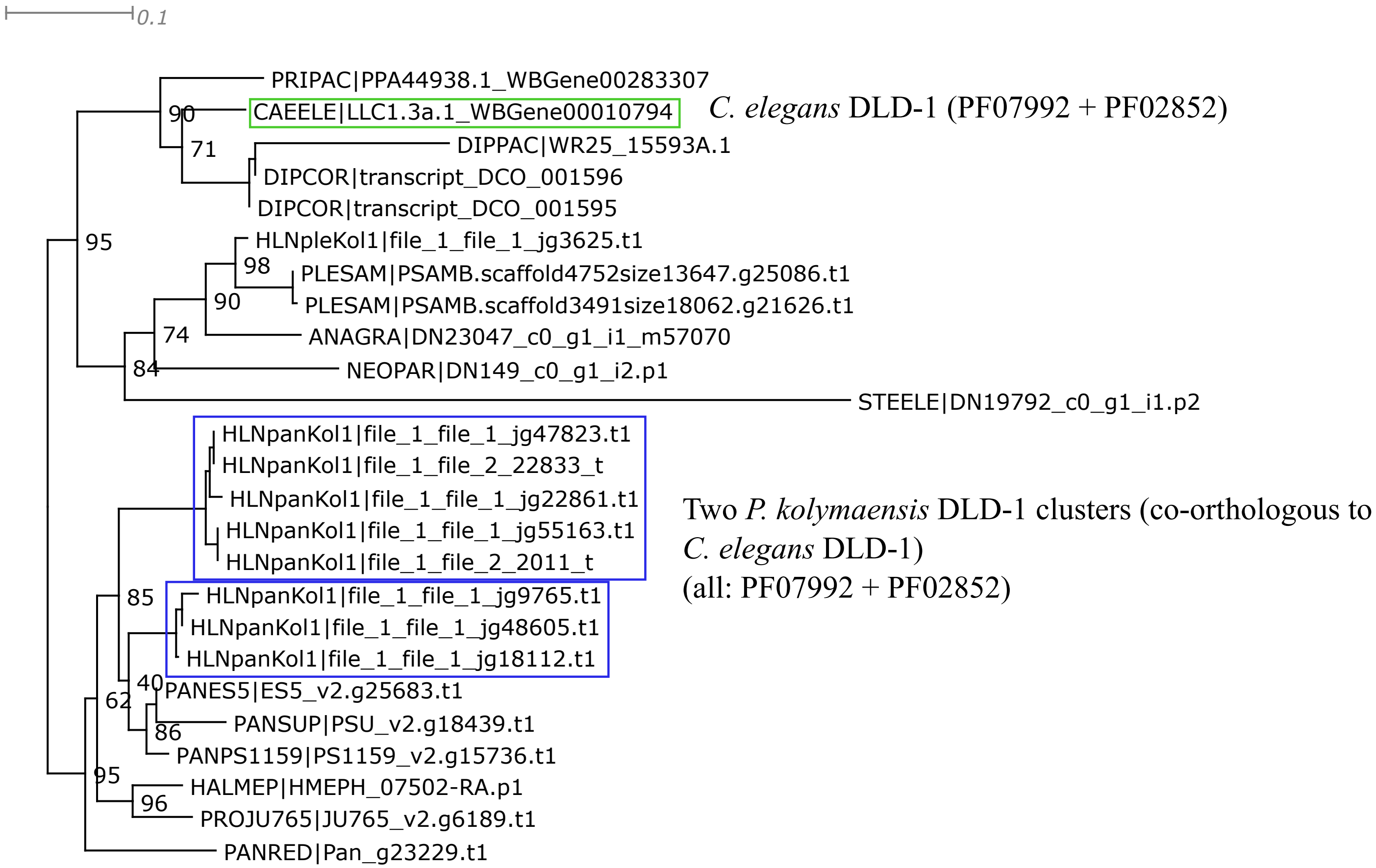


Trimal -automated1 function; short or spurious sequences manually removed afterwards;
IQtree2 ML phylogeny best-fit model according to BIC: LG+G4

PYC-1

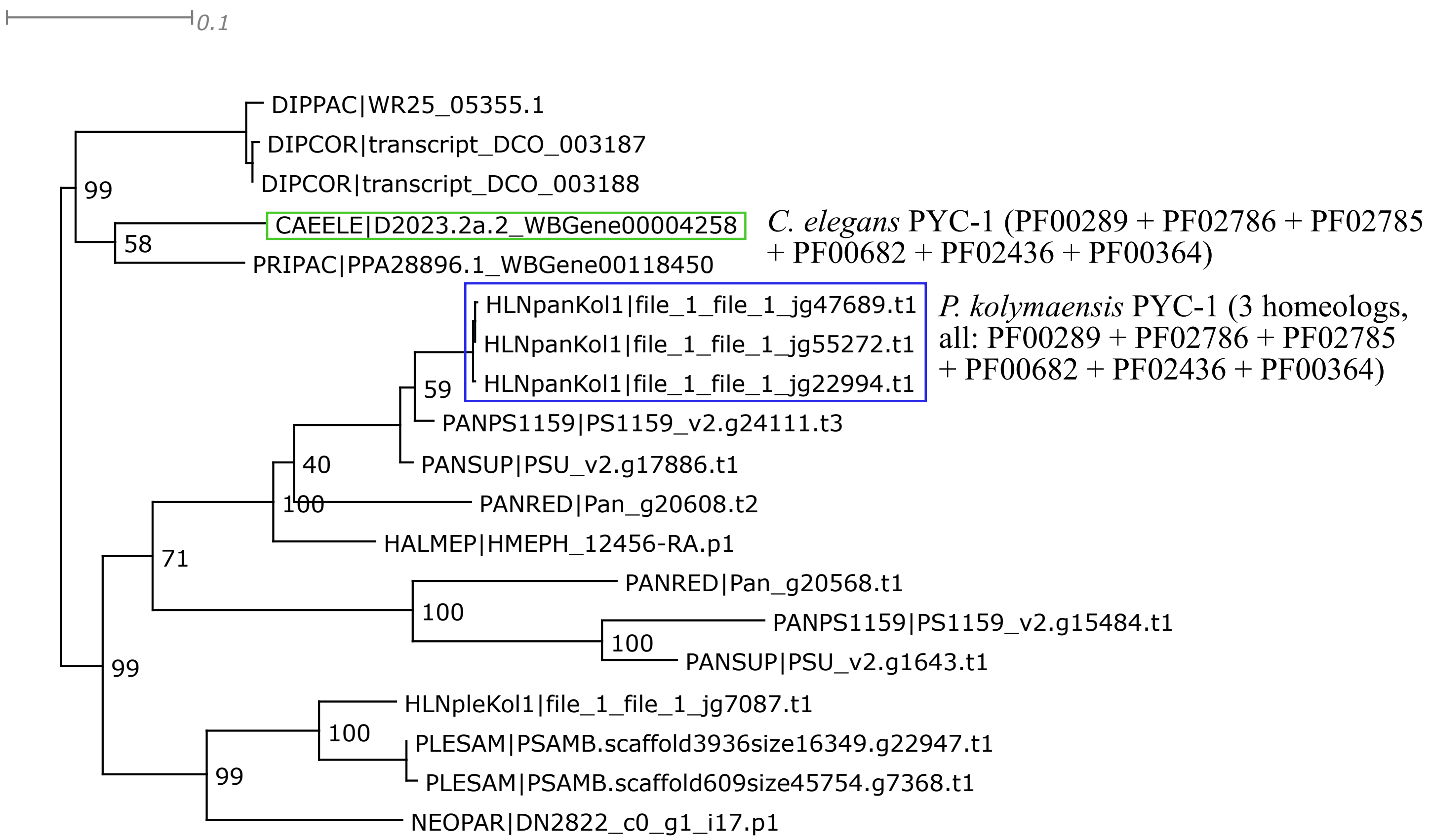


Trimal -automated1 function; short or spurious sequences manually removed afterwards;
IQtree2 ML phylogeny best-fit model according to BIC: LG+G4

PCK-1 / PCK-2

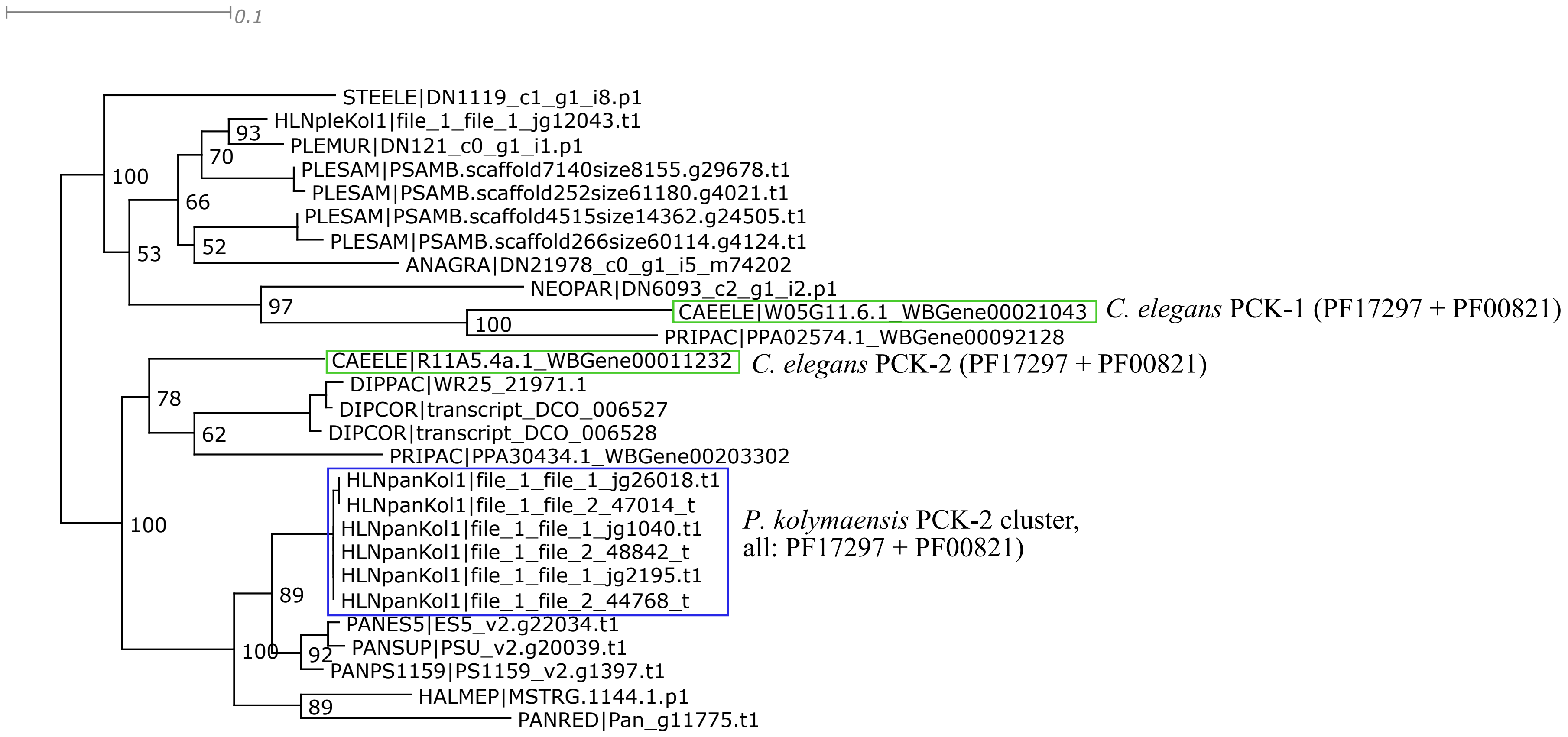


Trimal -automated1 function; short or spurious sequences manually removed afterwards;
IQtree2 ML phylogeny best-fit model according to BIC: LG+I+G4

PYK-1 / PYK-2

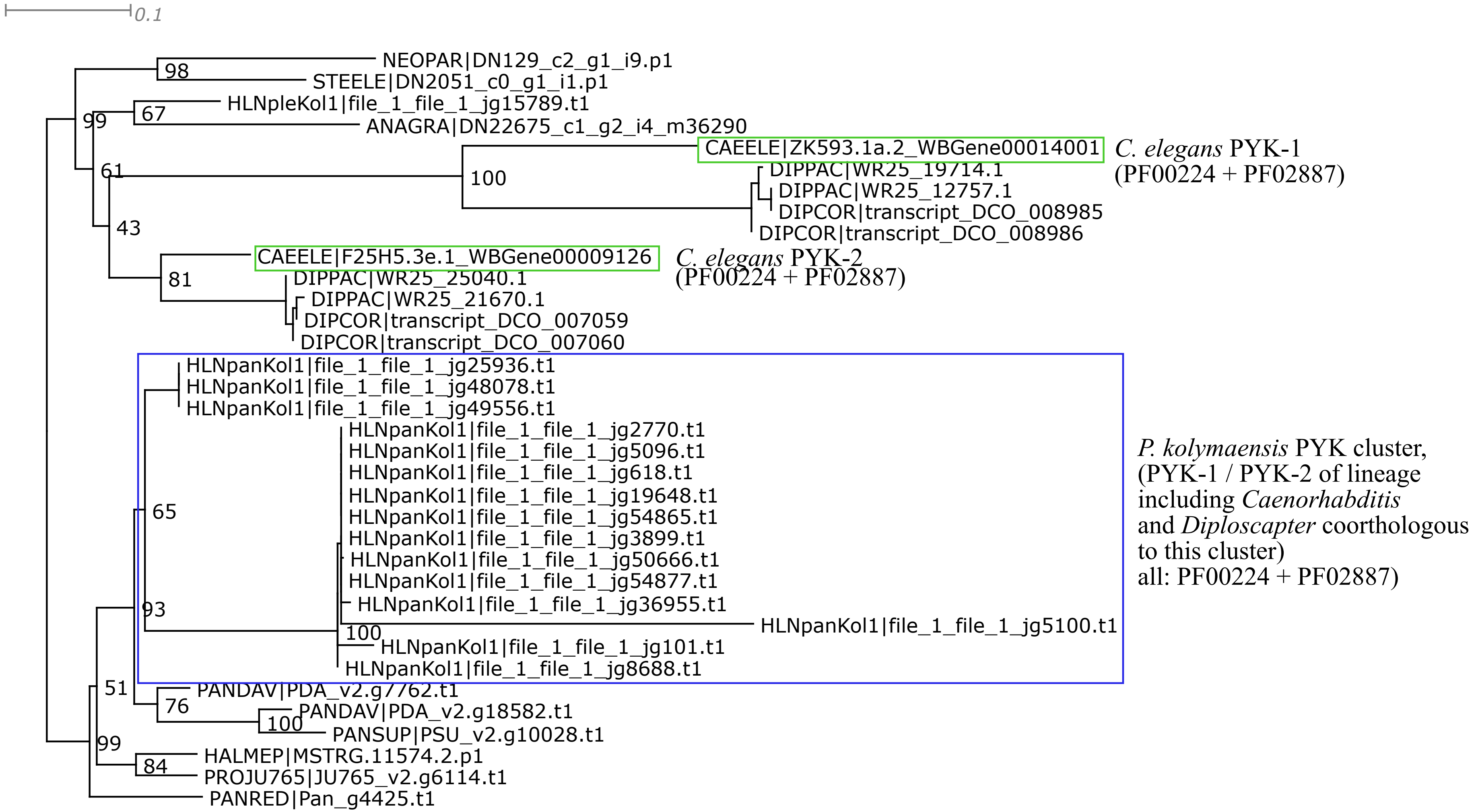


Trimal -automated1 function; short or spurious sequences manually removed afterwards;
IQtree2 ML phylogeny best-fit model according to BIC: LG+G4

ENOL-1

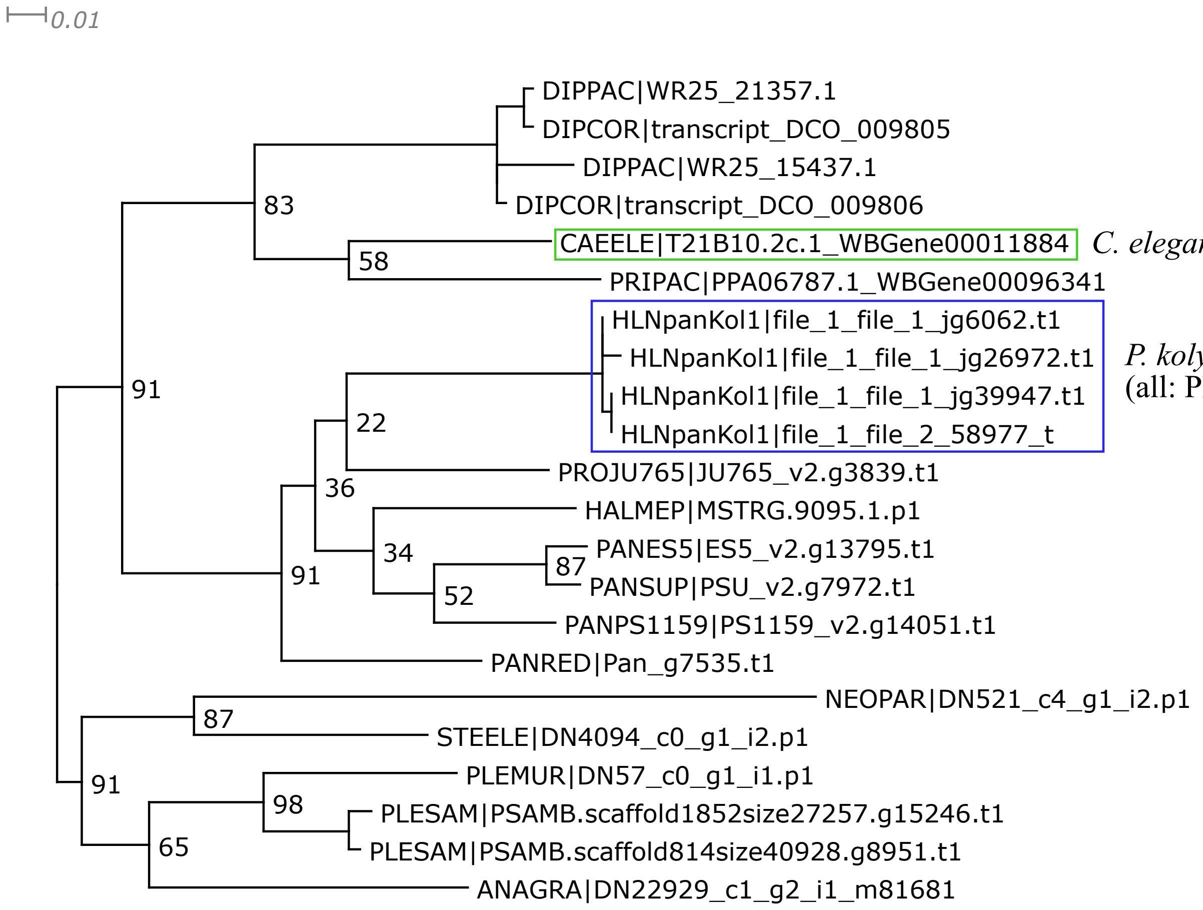


*C. elegans* ENOL-1 (PF03952 + PF00113)

*P. kolymaensis* ENOL-1 cluster,
(all: PF03952 + PF00113)

Trimal: 1. -resoverlap 0.75 -seqoverlap 80 functions; 2. -automated1 function; short or spurious sequences manually removed afterwards;
IQtree2 ML phylogeny best-fit model according to BIC: LG+G4

IPGM-1

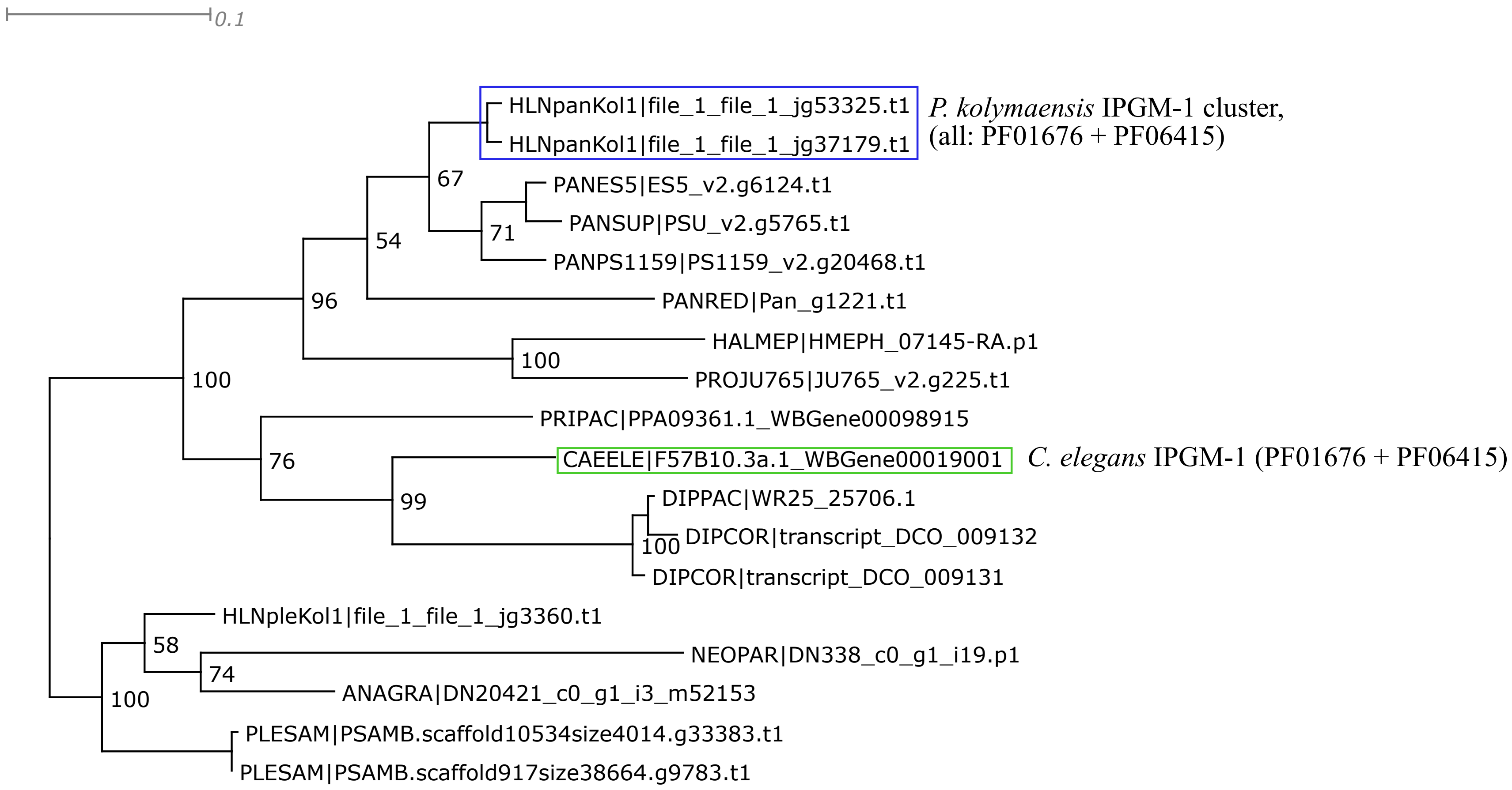


Trimal -automated1 function; short or spurious sequences manually removed afterwards;
IQtree2 ML phylogeny best-fit model according to BIC: LG+G4

PGK-1

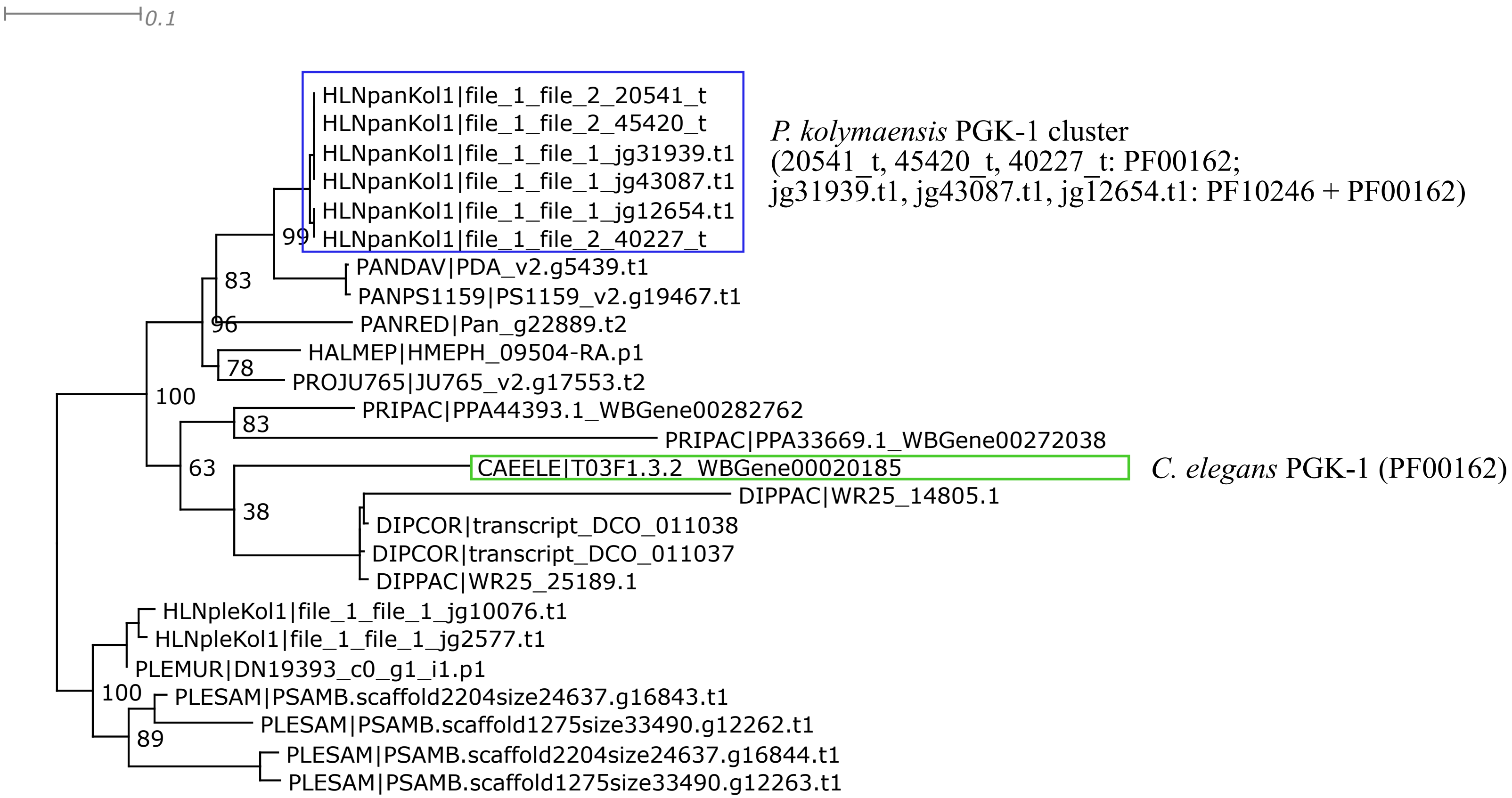


Trimal -automated1 function; short or spurious sequences manually removed afterwards;
IQtree2 ML phylogeny best-fit model according to BIC: WAG+G4

TPI-1

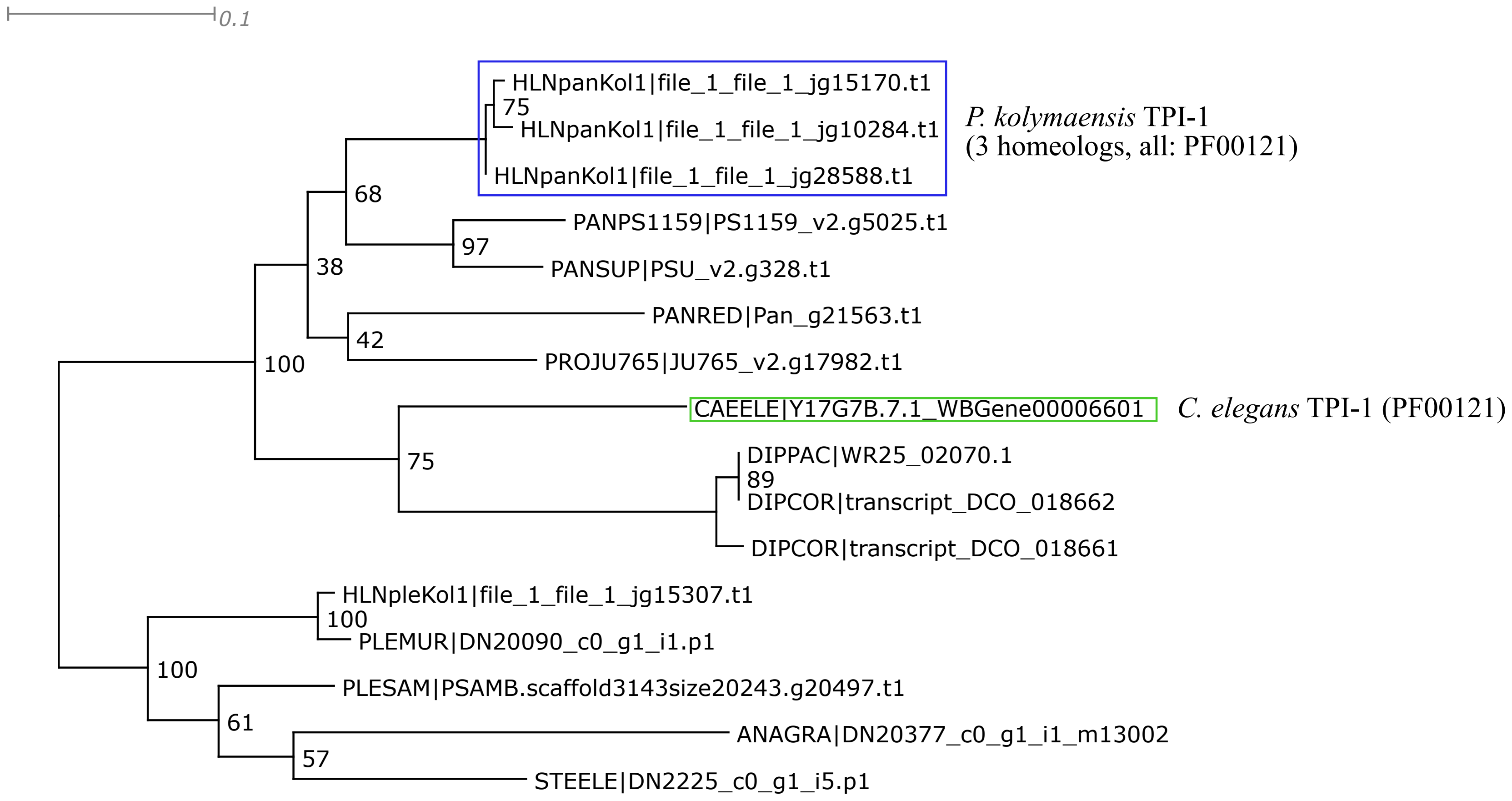


Trimal: 1. -resoverlap 0.75 -seqoverlap 80 functions; 2. -automated1 function; short or spurious sequences manually removed afterwards;
IQtree2 ML phylogeny best-fit model according to BIC: LG+G4

GPD-1, GPD-2, GPD-3, GPD-4

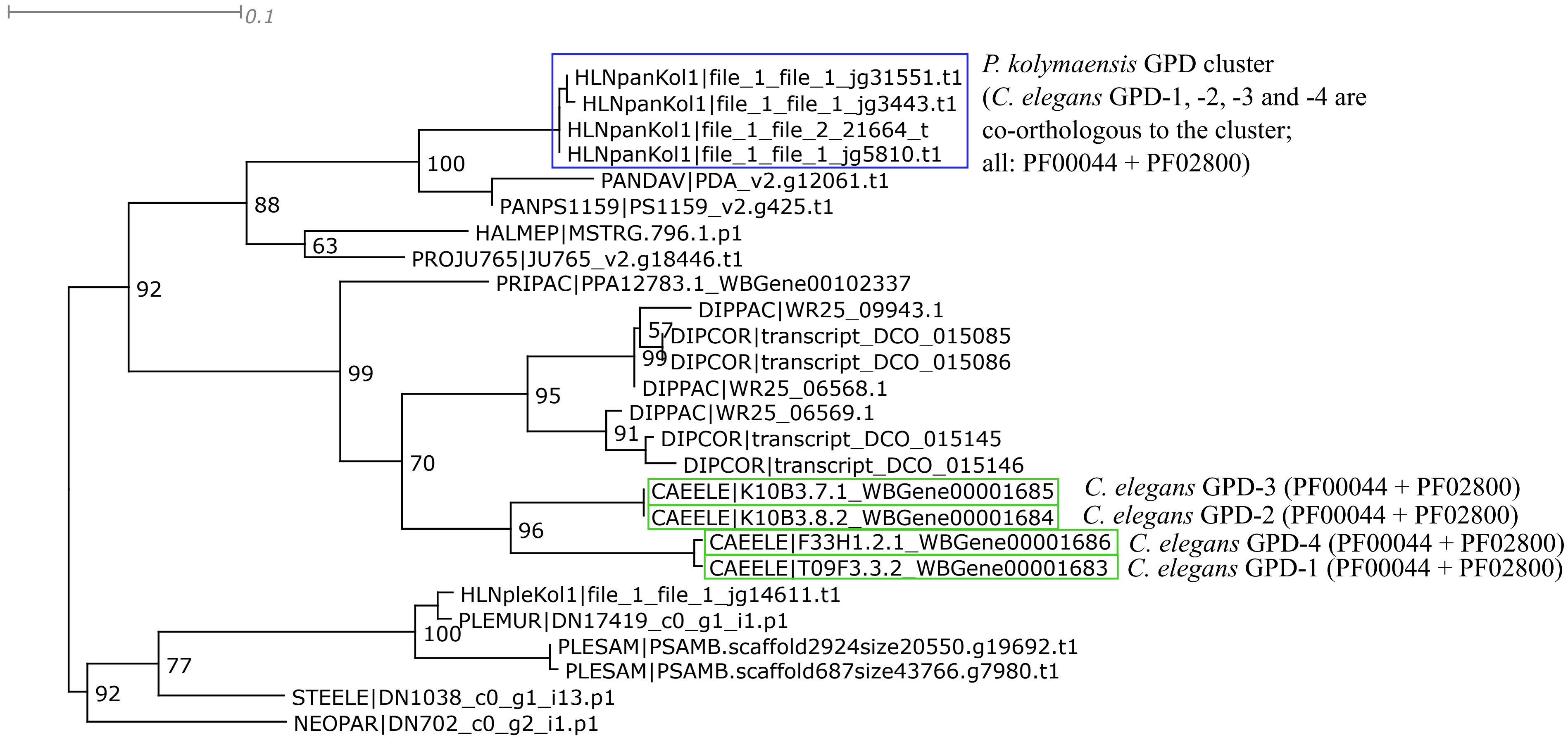


Trimal: 1. -resoverlap 0.75 -seqoverlap 80 functions; 2. -automated1 function; short or spurious sequences manually removed afterwards;
IQtree2 ML phylogeny best-fit model according to BIC: LG+G4

ALDO-1, ALDO-2

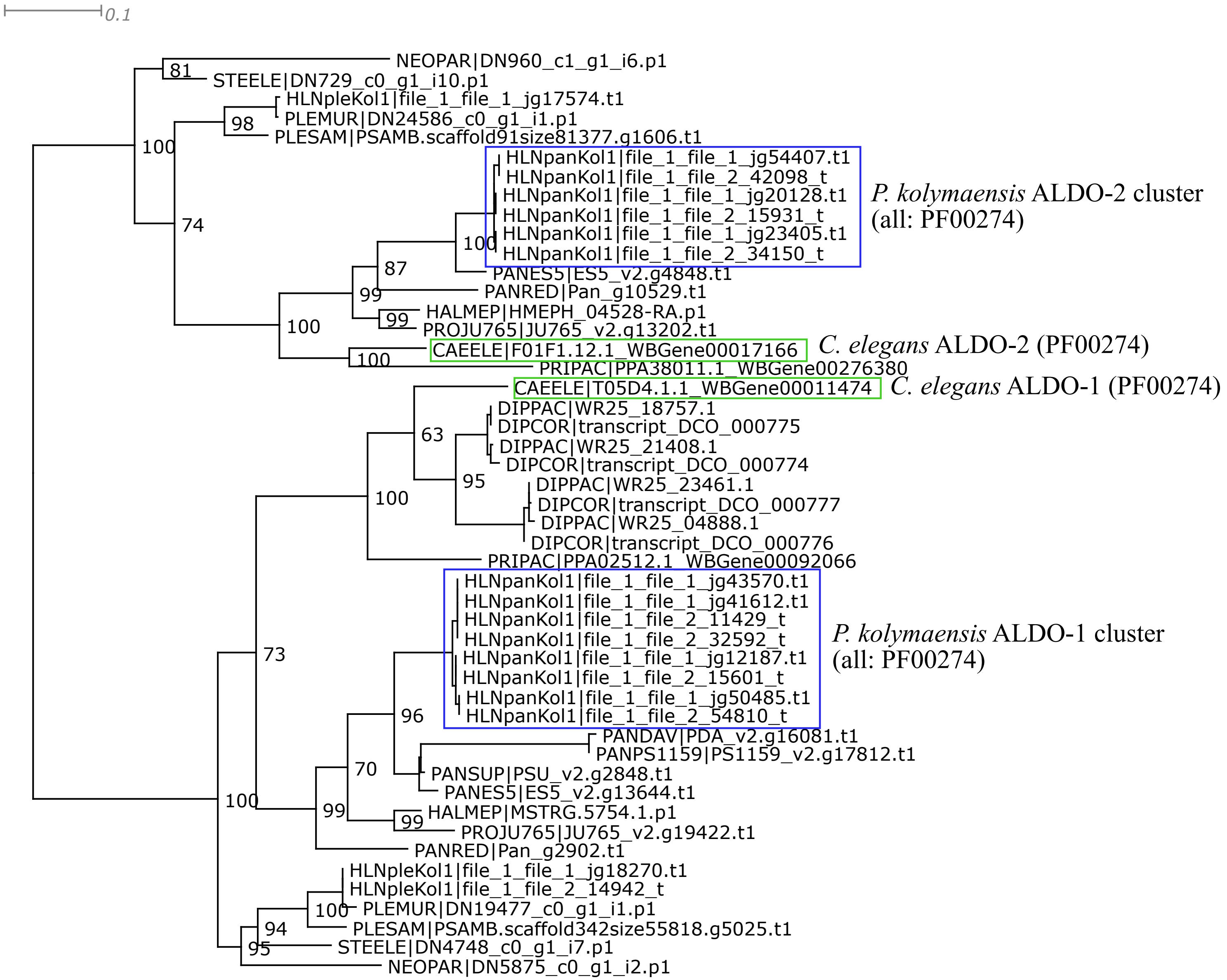


Trimal -automated1 function; short or spurious sequences manually removed afterwards;
IQtree2 ML phylogeny best-fit model according to BIC: LG+G4

FBP-1

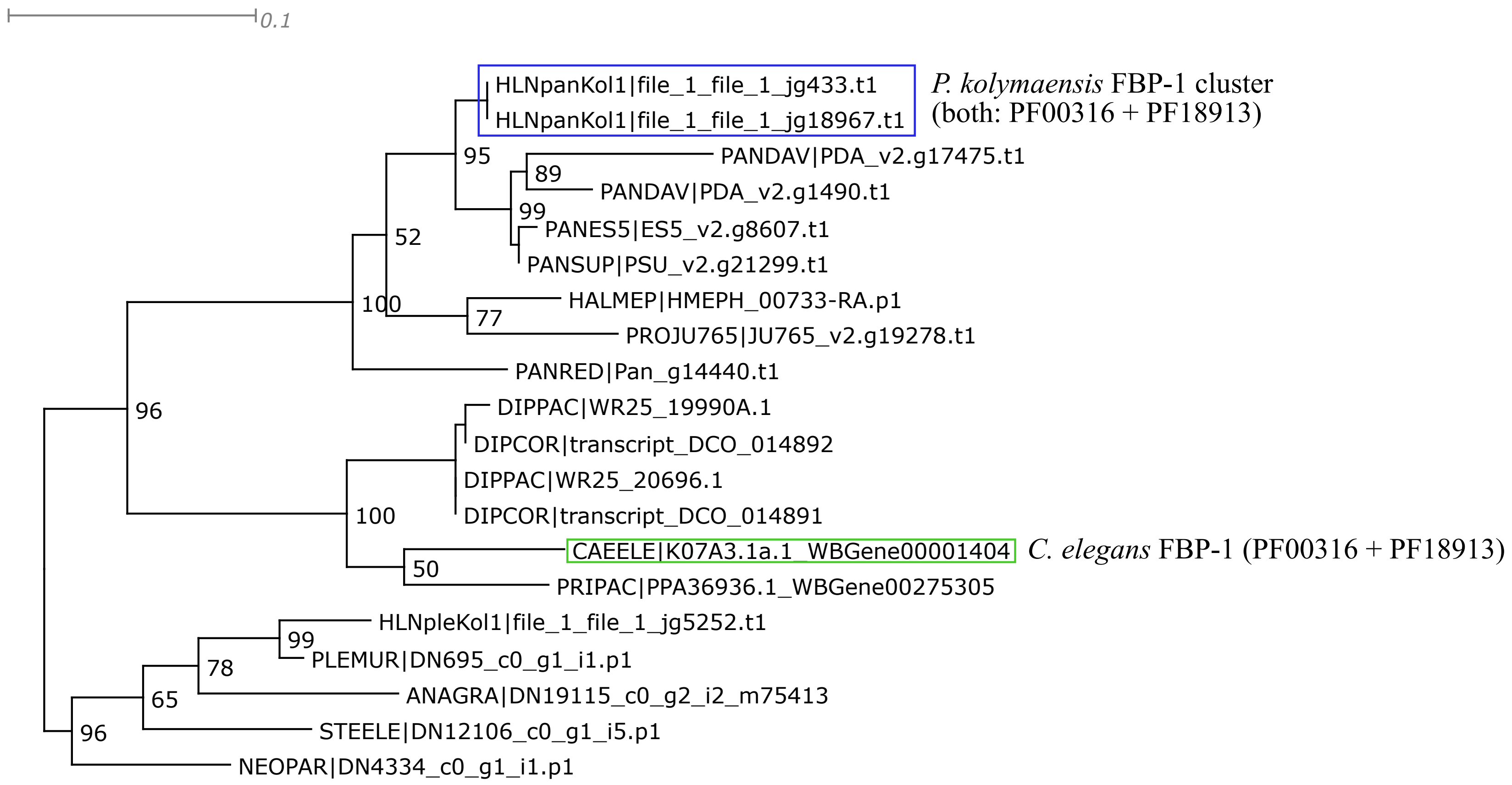


Trimal -automated1 function; short or spurious sequences manually removed afterwards;
IQtree2 ML phylogeny best-fit model according to BIC: WAG+G4

PFK-1

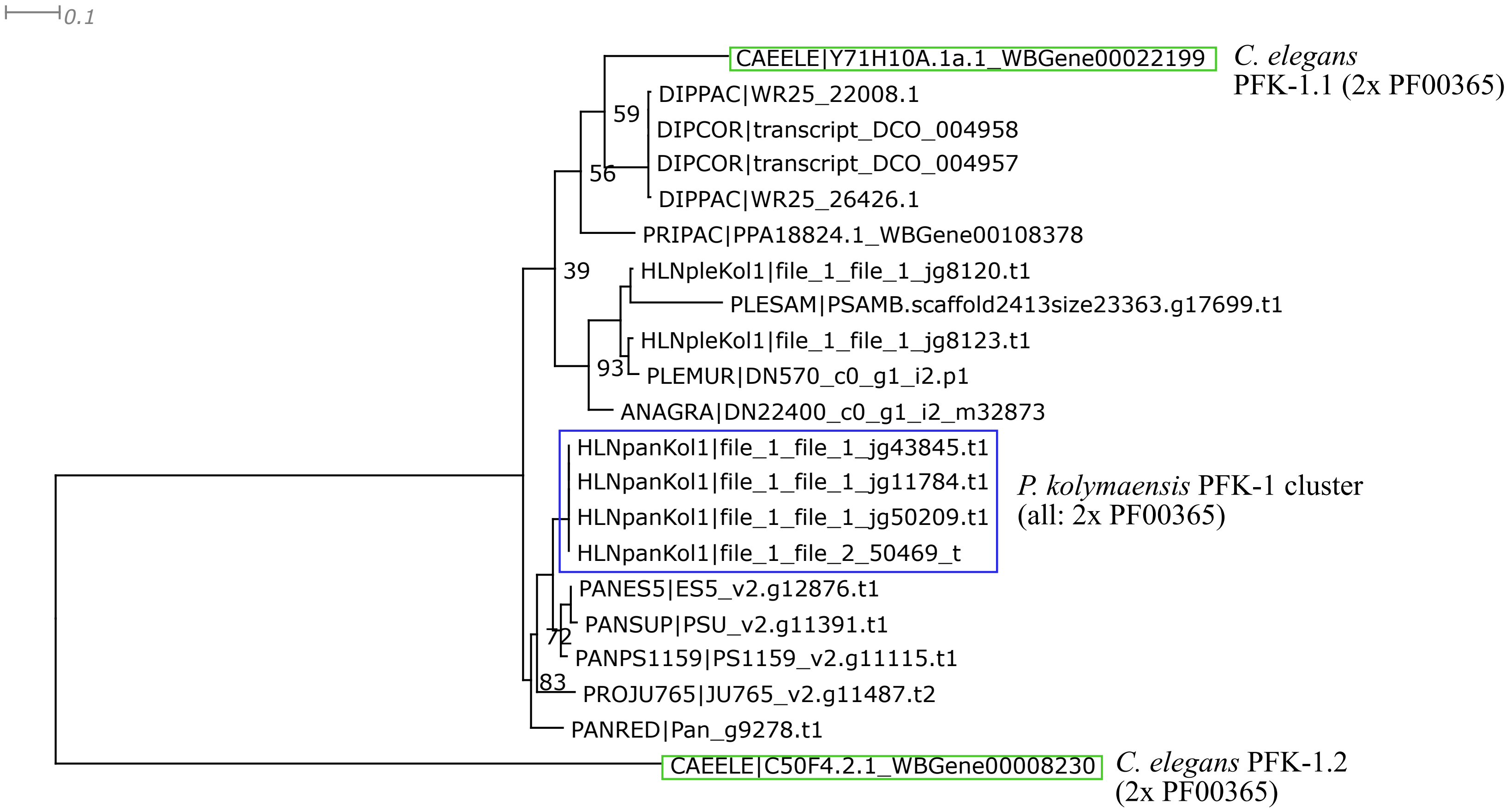


Trimal -automated1 function; short or spurious sequences (and sequences that disrupted phylogeny (N. parasiticus and H. mephisto) manually removed afterwards;
IQtree2 ML phylogeny best-fit model according to BIC: LG+G4

GPI-1

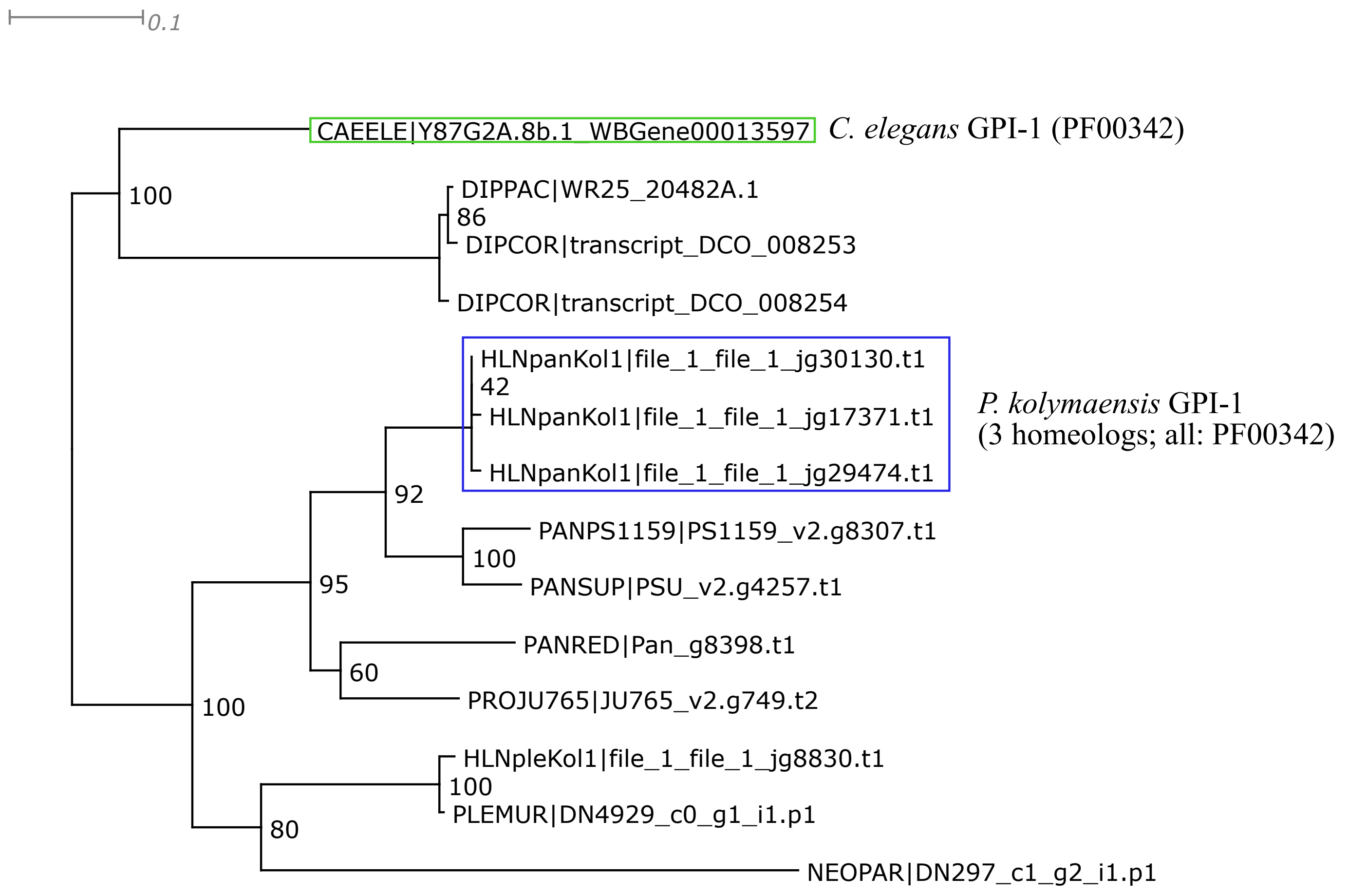


Trimal: 1. -resoverlap 0.75 -seqoverlap 80 functions; 2. -automated1 function; short or spurious sequences manually removed afterwards;
IQtree2 ML phylogeny best-fit model according to BIC: LG+G4

HXK-3

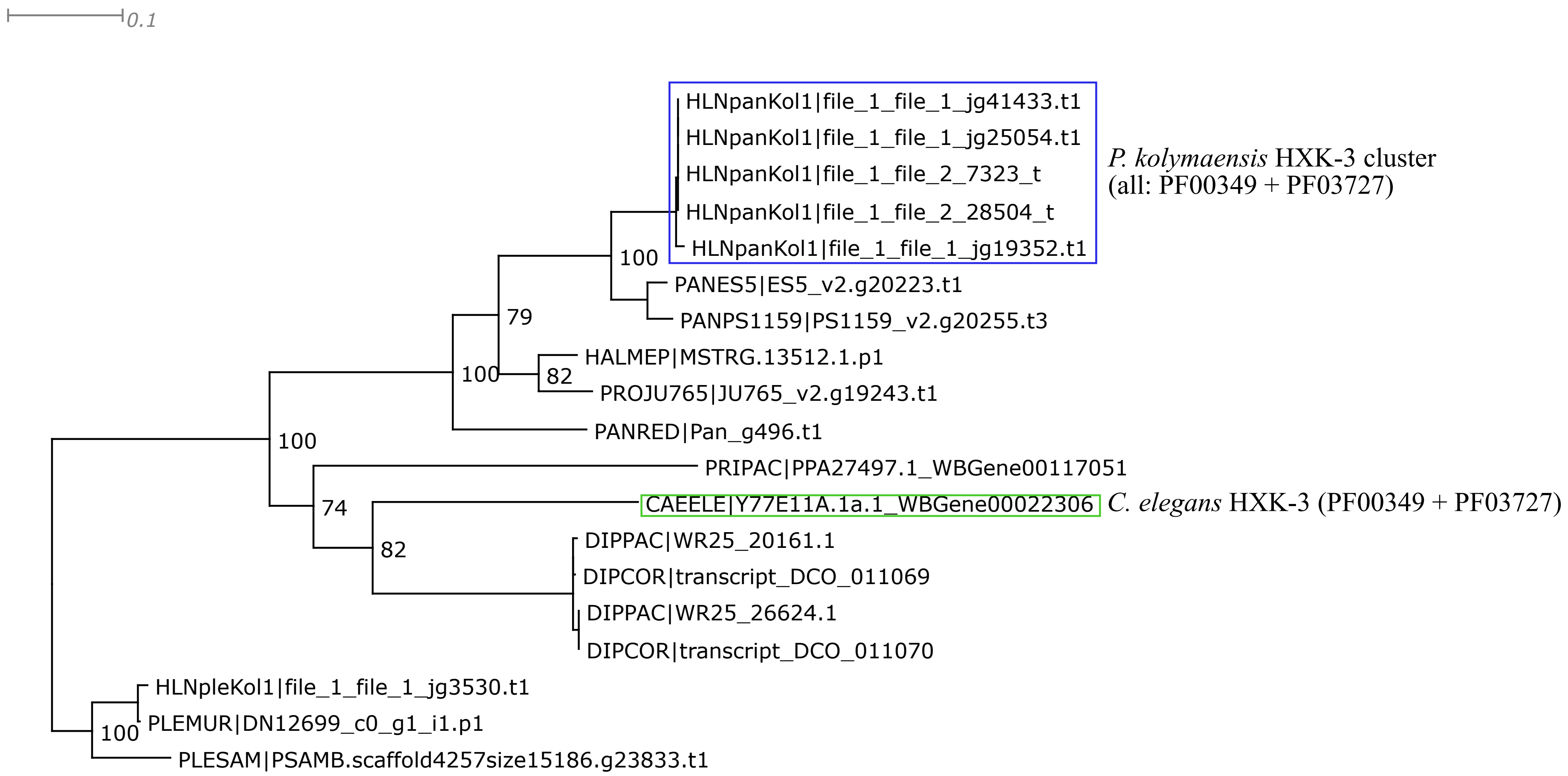


Trimal: 1. -resoverlap 0.75 -seqoverlap 80 functions; 2. -automated1 function; short or spurious sequences manually removed afterwards;
IQtree2 ML phylogeny best-fit model according to BIC: LG+G4

**Polyamine biosynthesis**

ODC-1

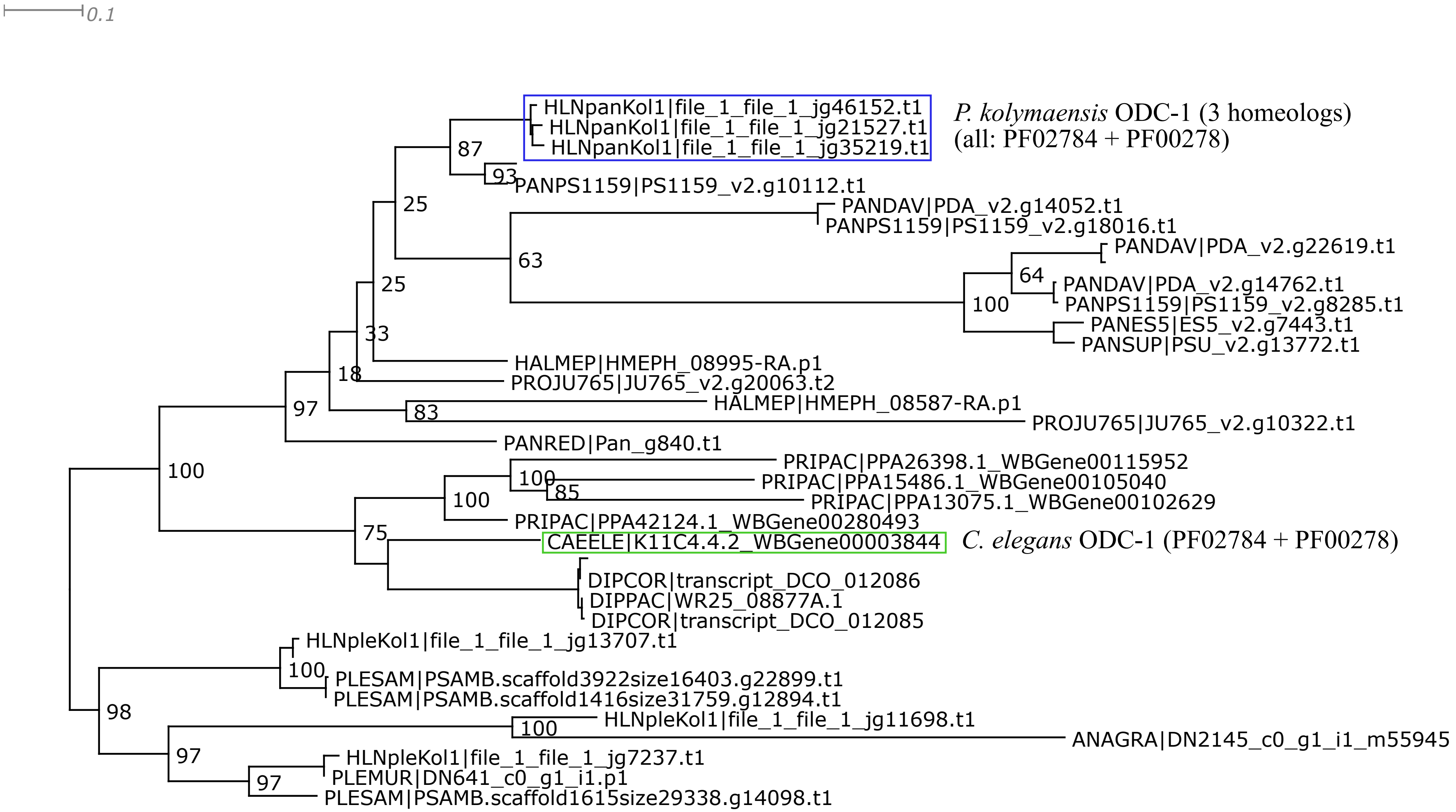


Trimal: 1. -resoverlap 0.75 -seqoverlap 80 functions; 2. -automated1 function; 3. short or spurious sequences manually removed afterwards; IQtree2 ML phylogeny best-fit model according to BIC: LG+I+G4

SPDS-1

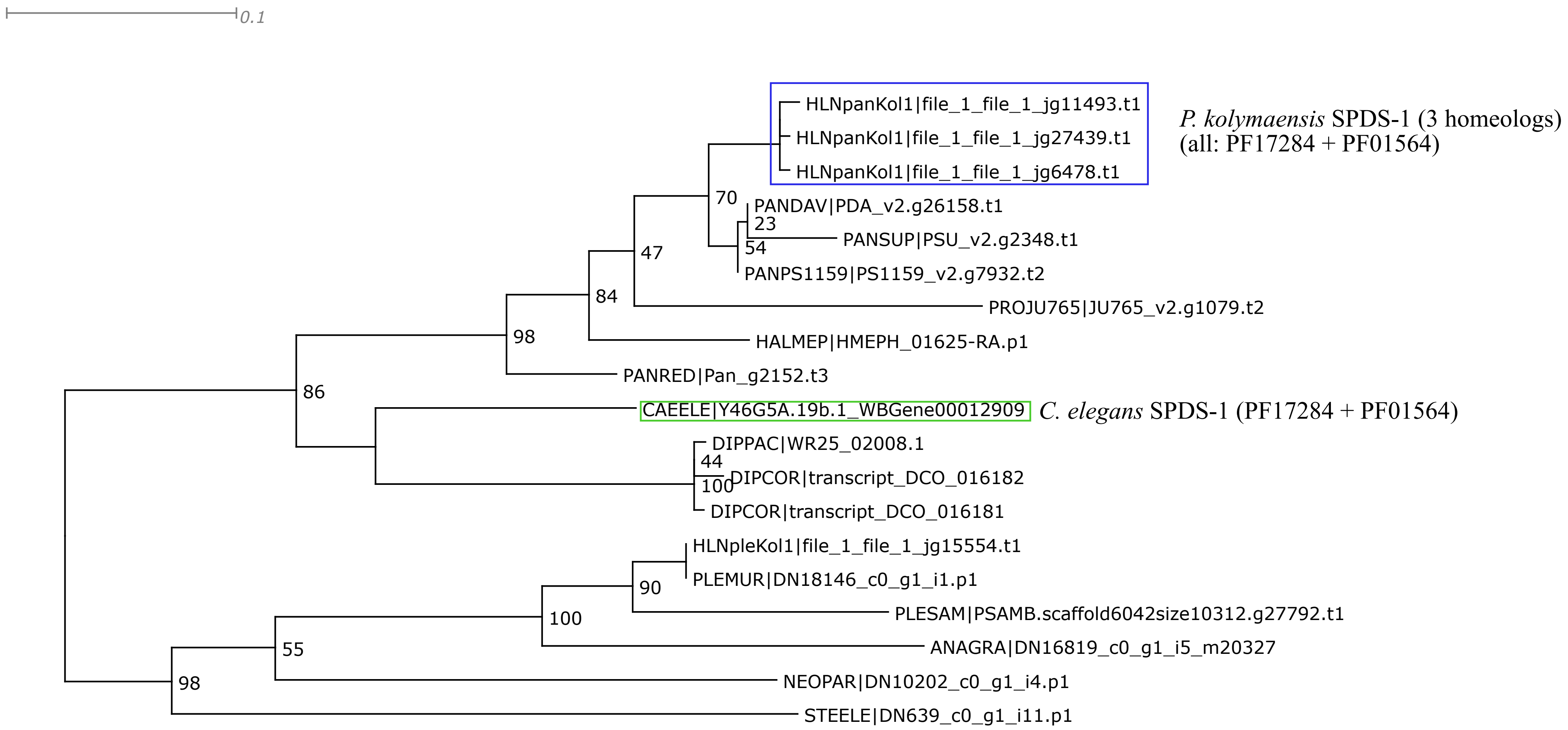


Trimal: 1. -resoverlap 0.75 -seqoverlap 80 functions; 2. -automated1 function; 3. short or spurious sequences manually removed afterwards; IQtree2 ML phylogeny best-fit model according to BIC: LG+G4

**Dauer genes**

DAF-1

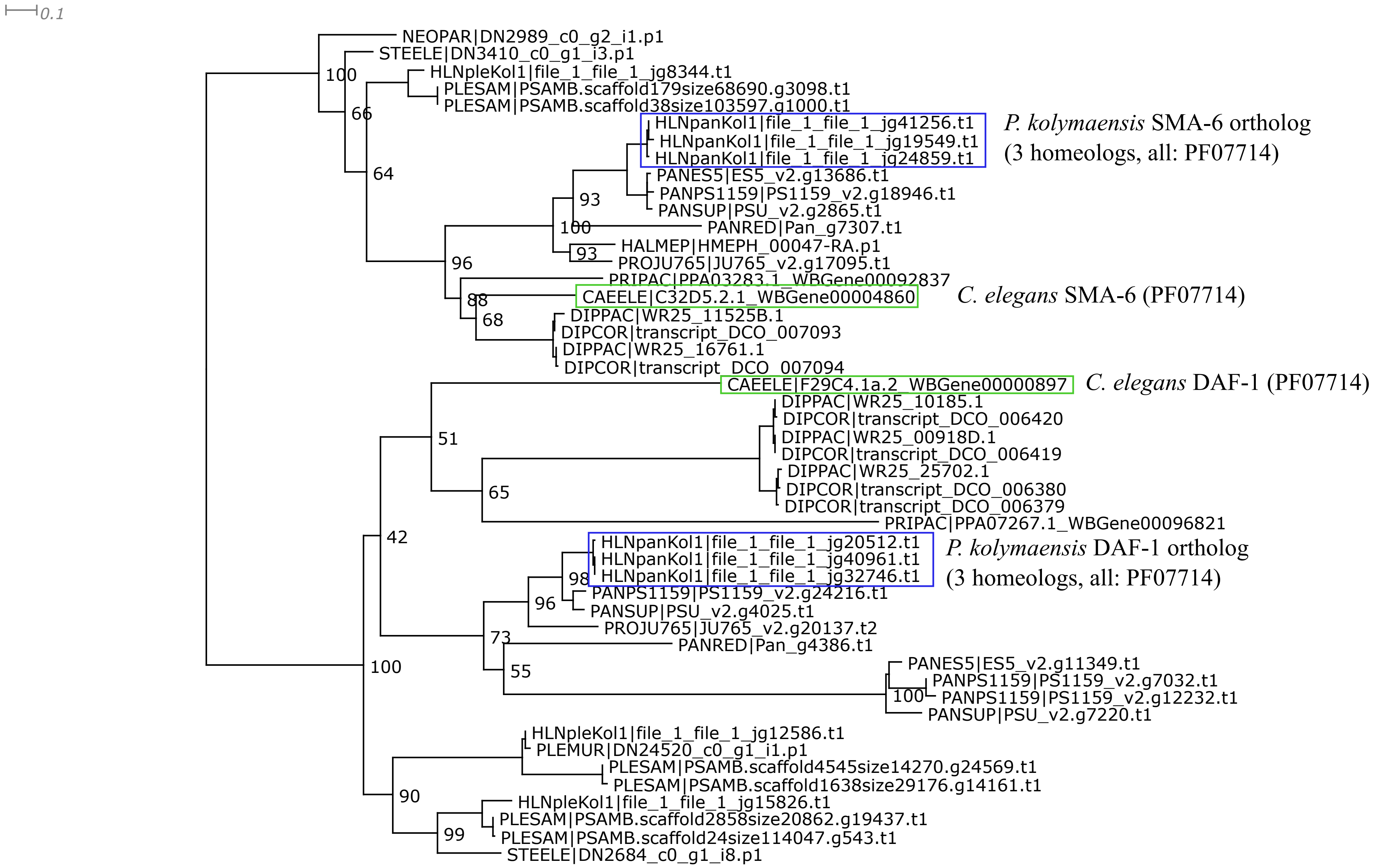


Trimal -automated1 function; short or spurious sequences manually removed afterwards;
IQtree2 ML phylogeny best-fit model according to BIC: LG+I+G4

DAF-2

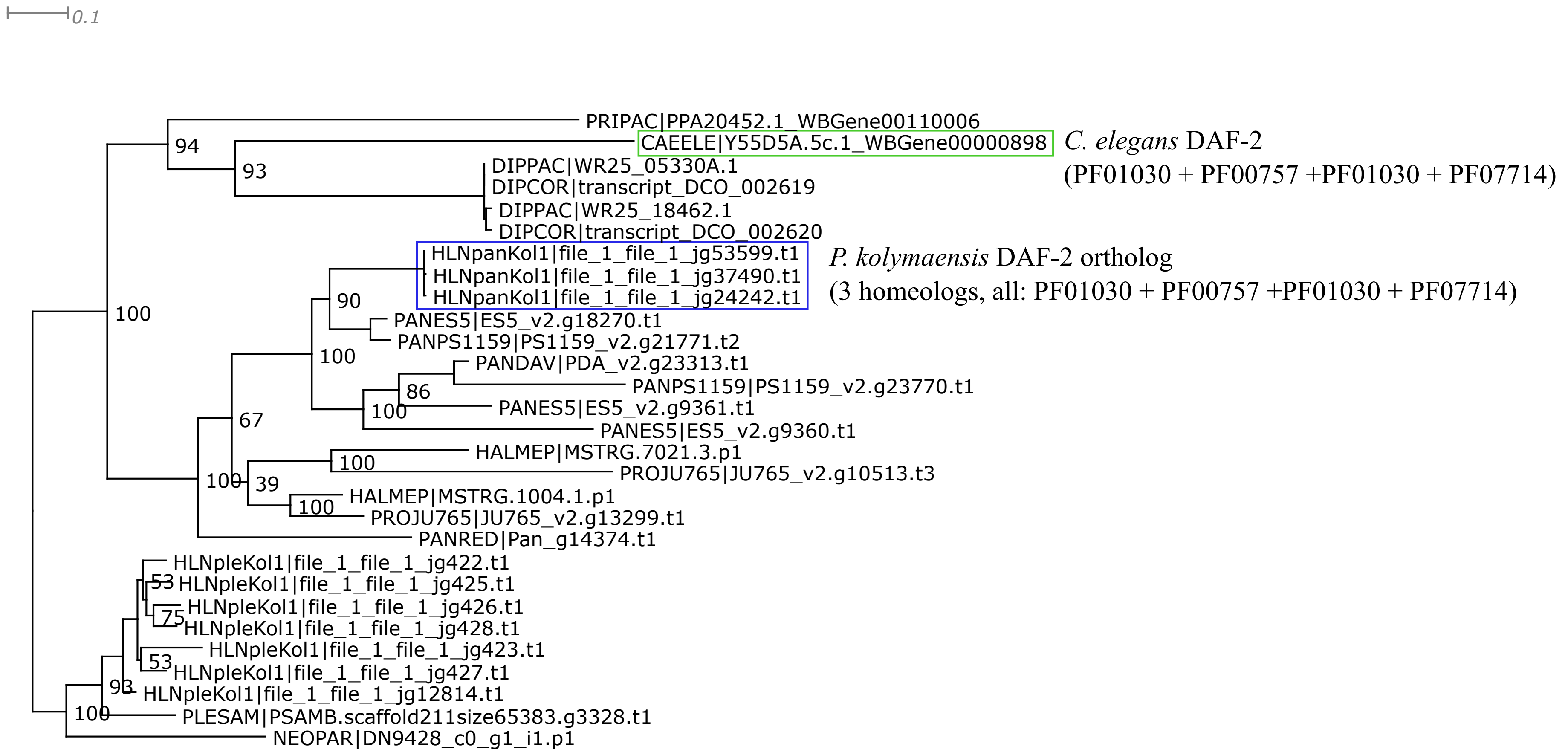


Trimal -automated1 function; Trimal functions -resoverlap 0.5 -seqoverlap 50;
IQtree2 ML phylogeny best-fit model according to BIC: LG+I+G4

DAF-4

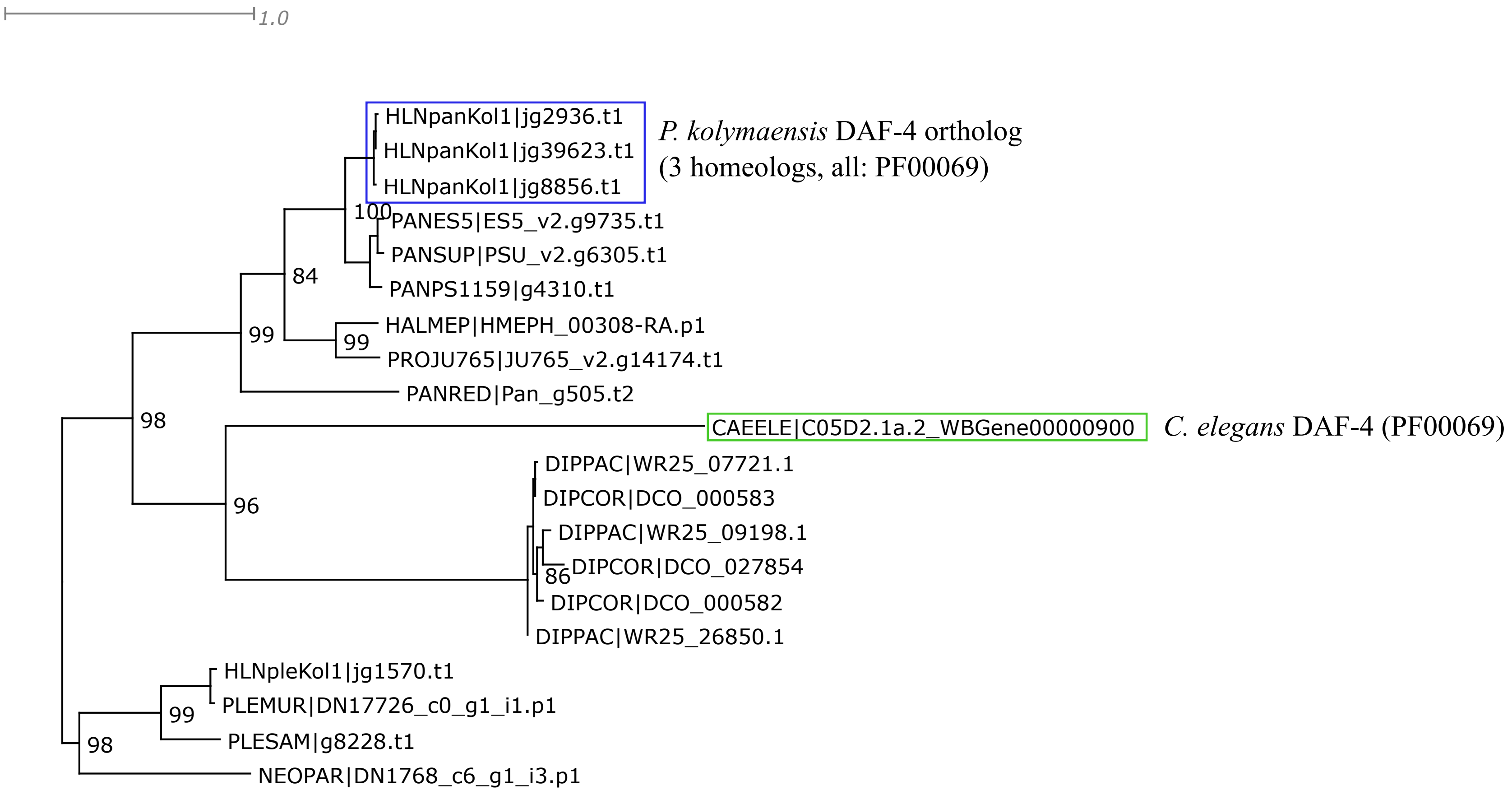


Trimal -automated1 function; short or spurious sequences manually removed afterwards;
IQtree2 ML phylogeny best-fit model according to BIC: LG+I+G4

DAF-6

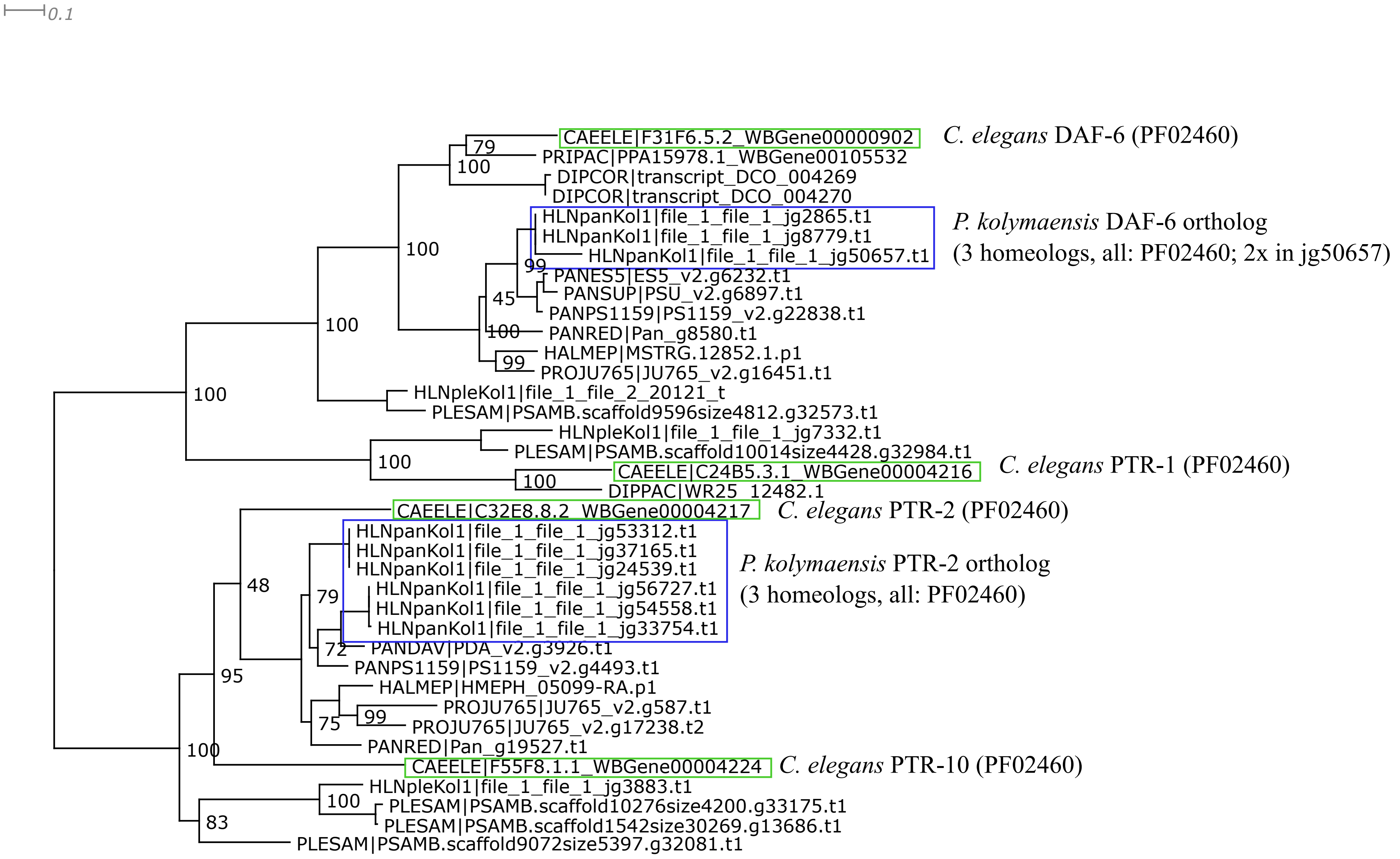


Trimal -automated1 function; Trimal function -resoverlap 0.75 -seqoverlap 75; more short or spurious sequences manually removed afterwards; IQtree2 ML phylogeny best-fit model according to BIC: LG+G4

DAF-7

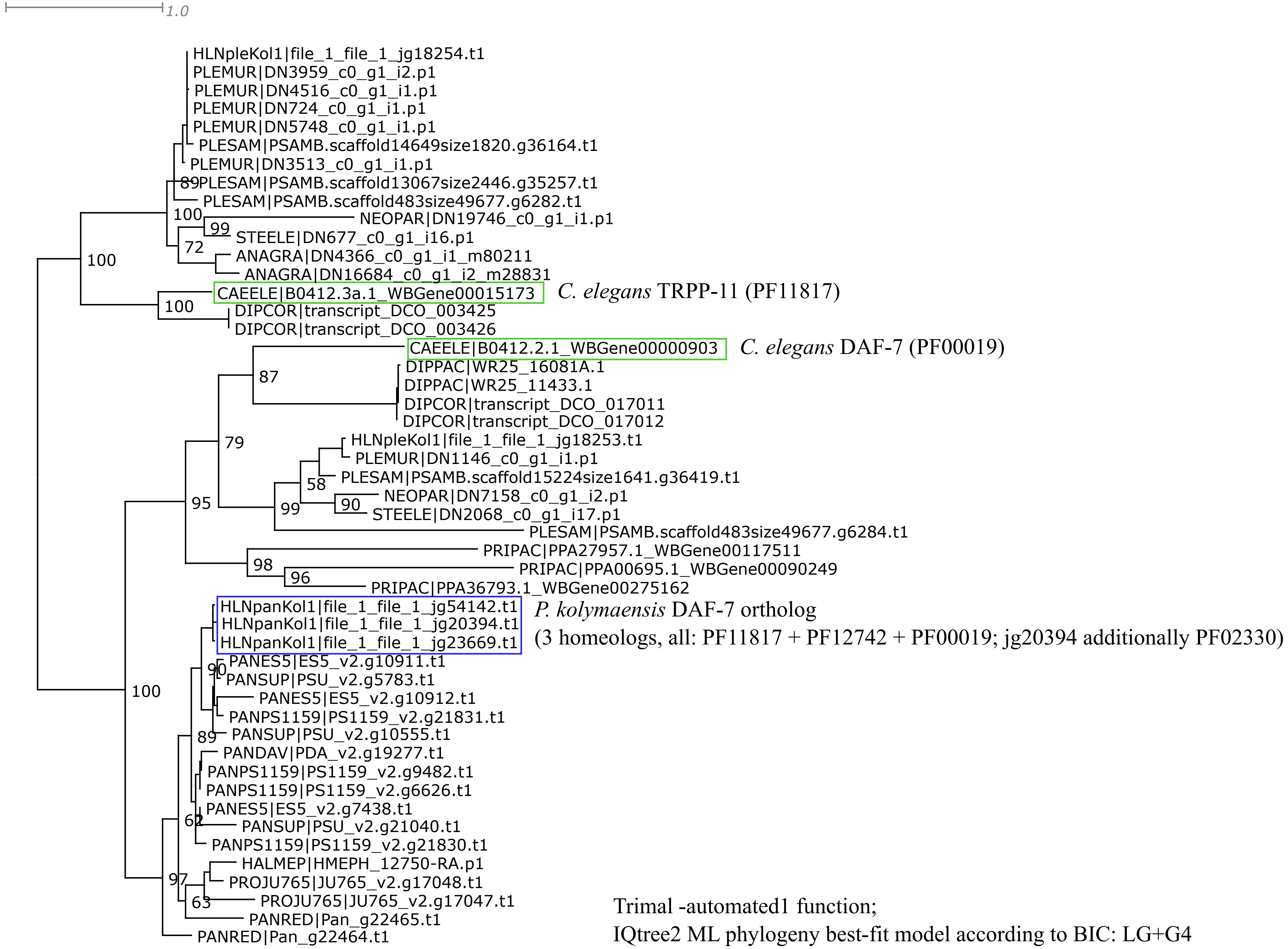

1.0
HLNpleKol1|file_1_file_1_jg18254.t1
PLEMUR|DN3959_c0_g1_i2.p1
PLEMUR|DN4516_c0_g1_i1.p1
PLEMUR|DN724_c0_g1_i1.p1
PLEMUR|DN5748_c0_g1_i1.p1
PLESAM|PSAMB.scaffold14649size1820.g36164.t1
PLEMUR|DN3513_c0_g1_i1.p1
89
PLESAM|PSAMB.scaffold13067size2446.g35257.t1
PLESAM|PSAMB.scaffold483size49677.g6282.t1
100
NEOPAR|DN19746_c0_g1_i1.p1
99
STEELE|DN677_c0_g1_i16.p1
72
ANAGRA|DN4366_c0_g1_i1_m80211
ANAGRA|DN16684_c0_g1_i2_m28831
100
CAEELE|B0412.3a.1_WBGene00015173
C. elegans TRPP-11 (PF11817)
100
DIPCOR|transcript_DCO_003425
DIPCOR|transcript_DCO_003426
CAEELE|B0412.2.1_WBGene00000903
C. elegans DAF-7 (PF00019)
87
DIPPAC|WR25_16081A.1
DIPPAC|WR25_11433.1
DIPCOR|transcript_DCO_017011
DIPCOR|transcript_DCO_017012
HLNpleKol1|file_1_file_1_jg18253.t1
79
PLEMUR|DN1146_c0_g1_i1.p1
PLESAM|PSAMB.scaffold15224size1641.g36419.t1
58
NEOPAR|DN7158_c0_g1_i2.p1
90
95
99
STEELE|DN2068_c0_g1_i17.p1
PLESAM|PSAMB.scaffold483size49677.g6284.t1
PRIPAC|PPA27957.1_WBGene00117511
98
PRIPAC|PPA00695.1_WBGene00090249
96
PRIPAC|PPA36793.1_WBGene00275162
HLNpanKol1|file_1_file_1_jg54142.t1
HLNpanKol1|file_1_file_1_jg20394.t1
HLNpanKol1|file_1_file_1_jg23669.t1
P. kolymaensis DAF-7 ortholog
(3 homeologs, all: PF11817 + PF12742 + PF00019; jg20394 additionally PF02330)
PANES5|ES5_v2.g10911.t1
90
PANSUP|PSU_v2.g5783.t1
100
PANES5|ES5_v2.g10912.t1
PANPS1159|PS1159_v2.g21831.t1
89
PANSUP|PSU_v2.g10555.t1
PANDAV|PDA_v2.g19277.t1
PANPS1159|PS1159_v2.g9482.t1
PANPS1159|PS1159_v2.g6626.t1
PANES5|ES5_v2.g7438.t1
62
PANSUP|PSU_v2.g21040.t1
PANPS1159|PS1159_v2.g21830.t1
HALMEP|HMEPH_12750-RA.p1
97
PROJU765|JU765_v2.g17048.t1
63
PROJU765|JU765_v2.g17047.t1
PANRED|Pan_g22465.t1
PANRED|Pan_g22464.t1
Trimal -automated1 function;
IQtree2 ML phylogeny best-fit model according to BIC: LG+G4

DAF-8 and DAF-14

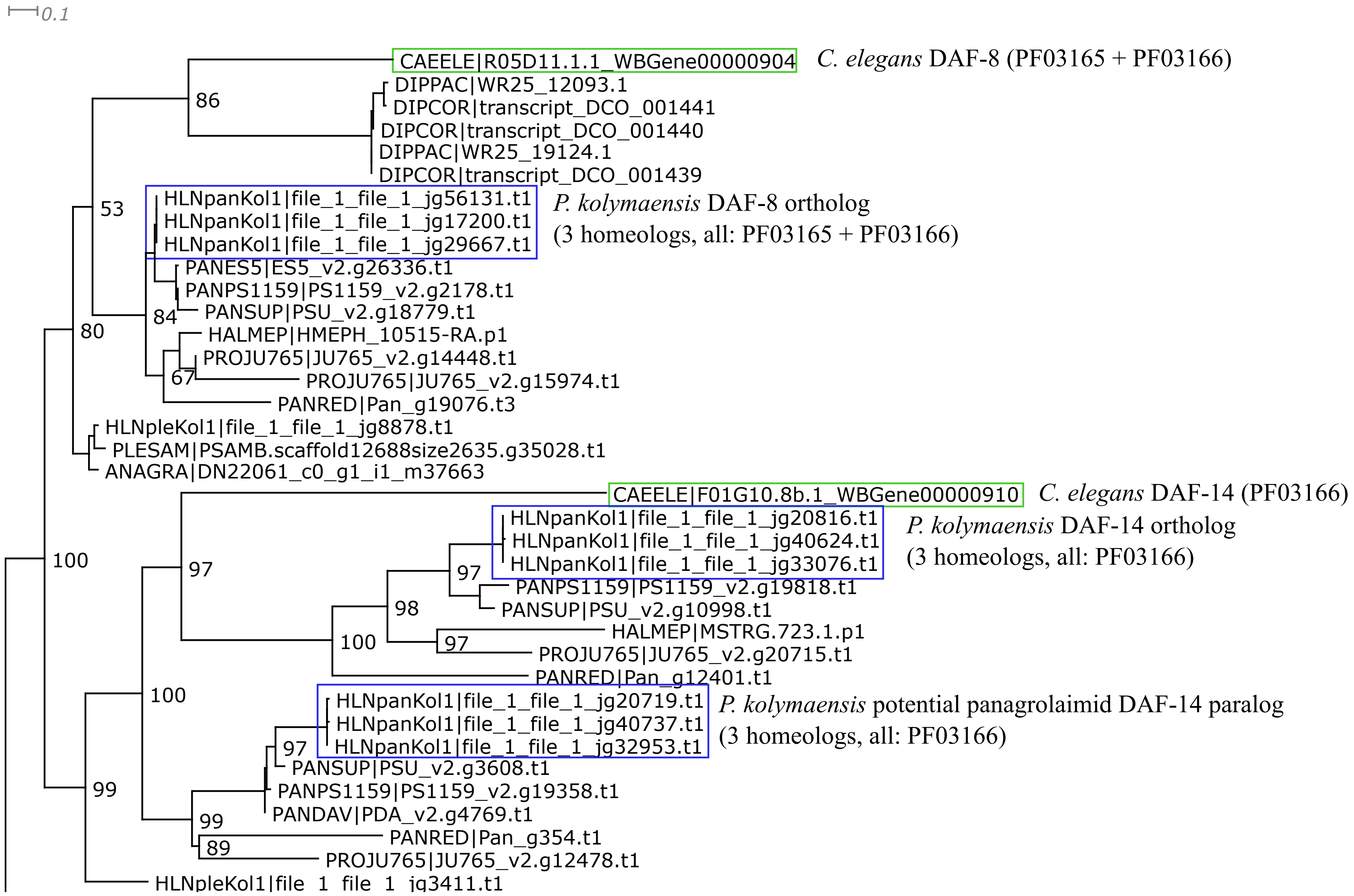


Partial phylogeny; clusters not shown here: SMA-2 orthologs, and SMA-3 orthologs;
Trimal -automated1 function; short and divergent sequences removed manually
IQtree2 ML phylogeny best-fit model according to BIC: LG+I+G4

DAF-9

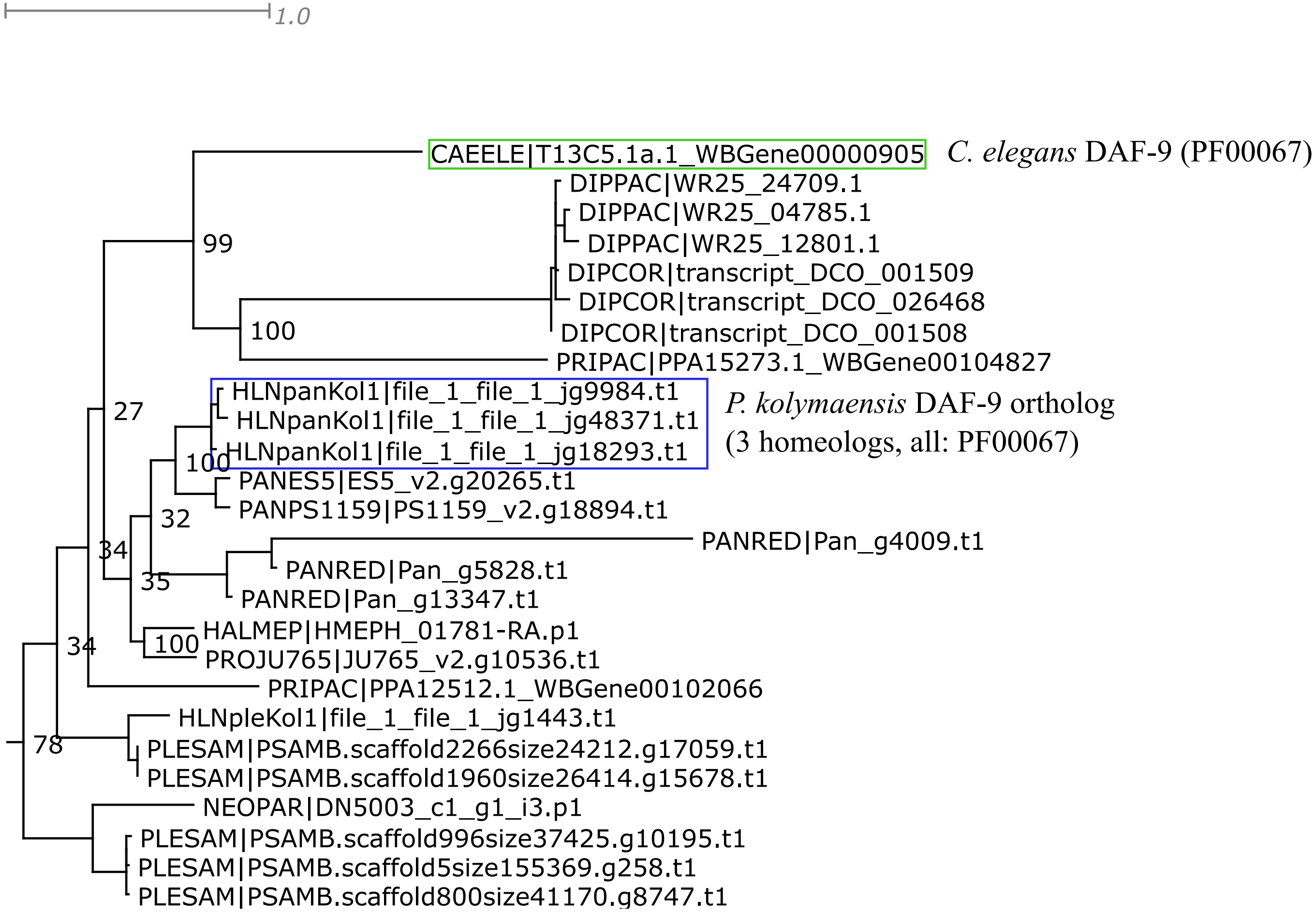


Partial phylogeny; the DAF-9 cluster is only one of many clusters in a large phylogeny of cytochrome P450 genes;
Trimal -automated1 function; short and divergent sequences removed manually
IQtree2 ML phylogeny best-fit model according to BIC: LG+G4

DAF-10

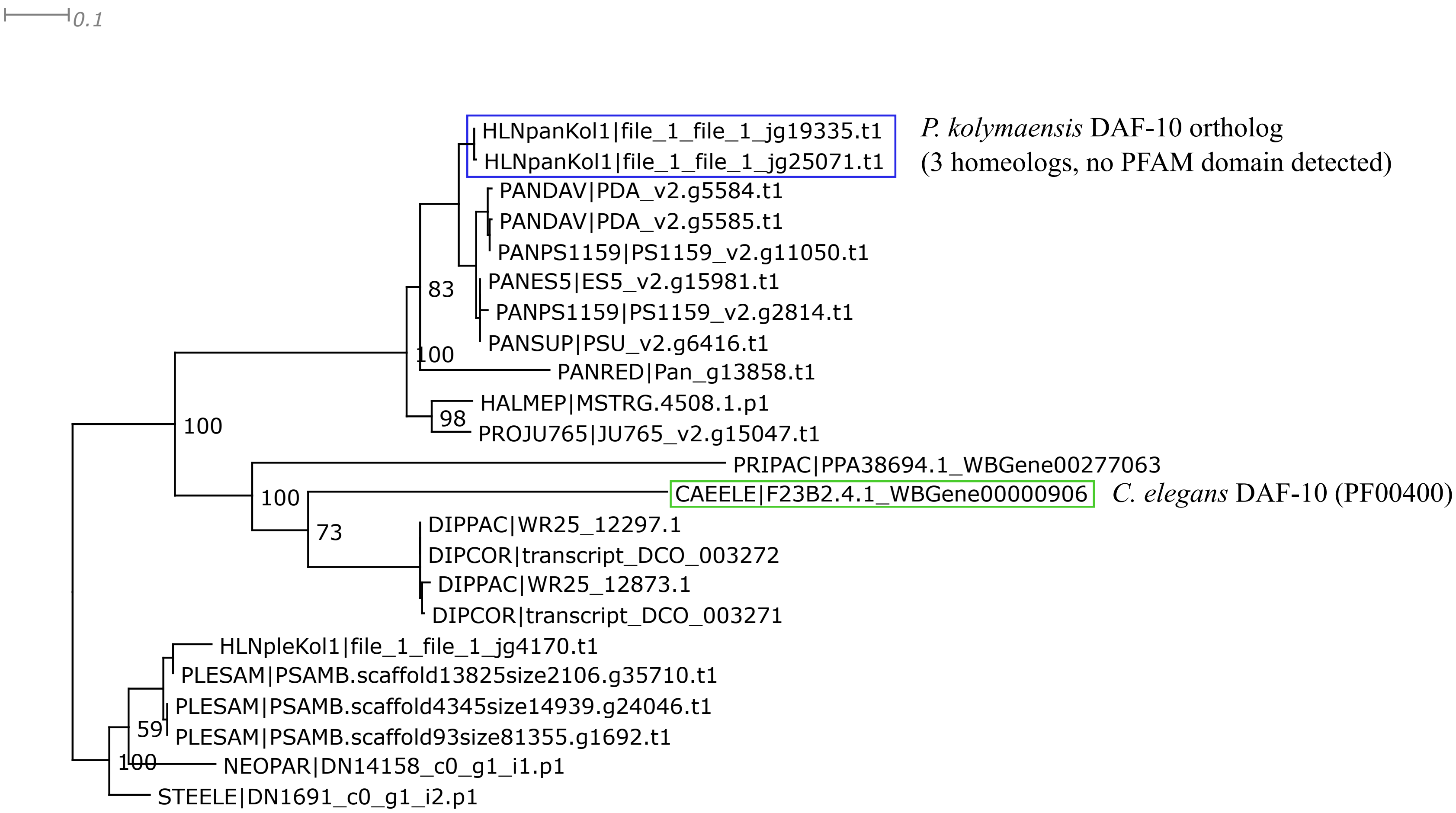


Trimal -automated1 function; .
IQtree2 ML phylogeny best-fit model according to BIC: LG+G4
The PFAM domain PF00400 found in the *C. elegans* DAF-10 protein is not detected in many panagrolaimids, but in some (*P. redivivus*, *Propanagrolaimus* sp. JU765, *H. mephisto*), it might be diverged and not recognised.

DAF-11

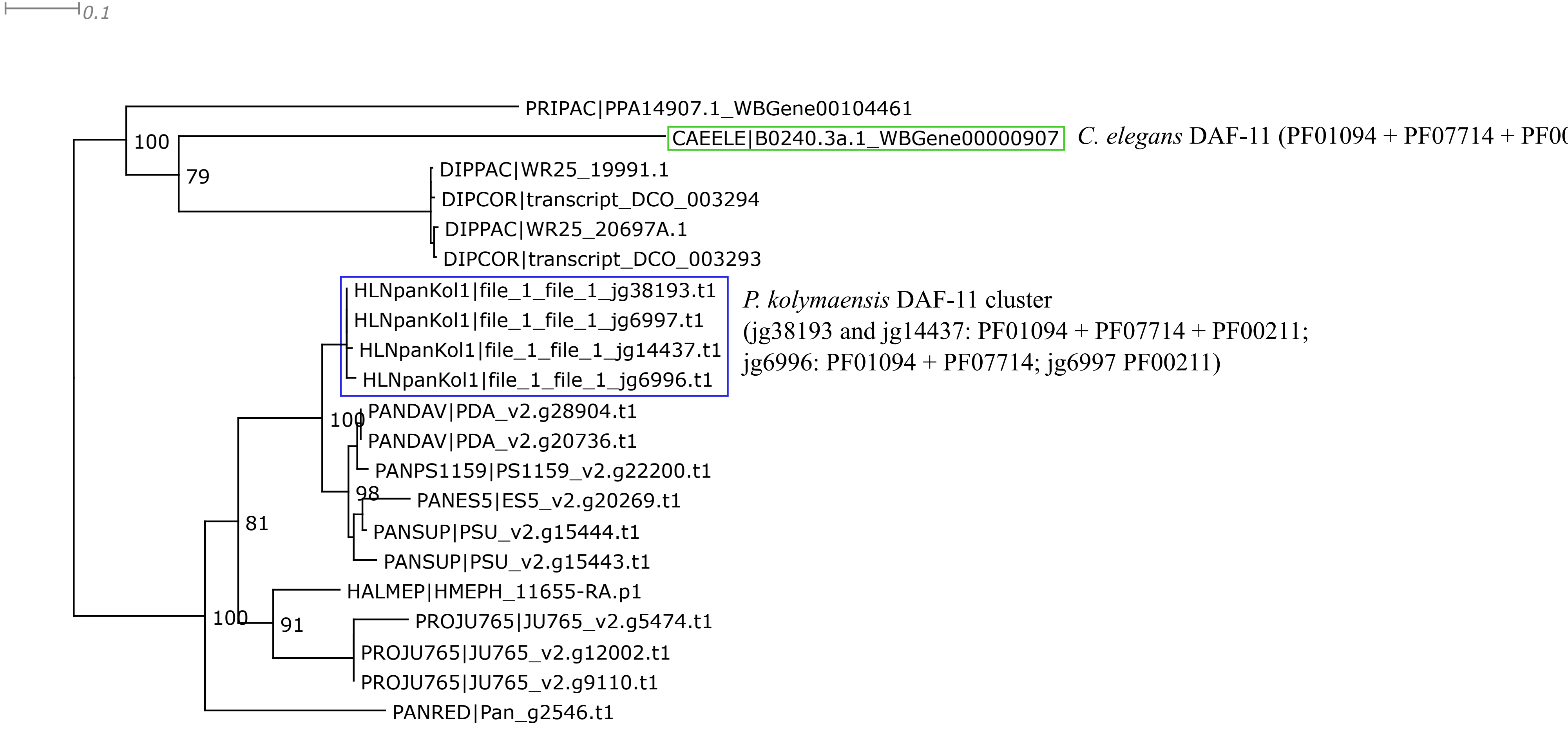


Trimal -automated1 function; .
IQtree2 ML phylogeny best-fit model according to BIC: LG+I+G4.

DAF-12

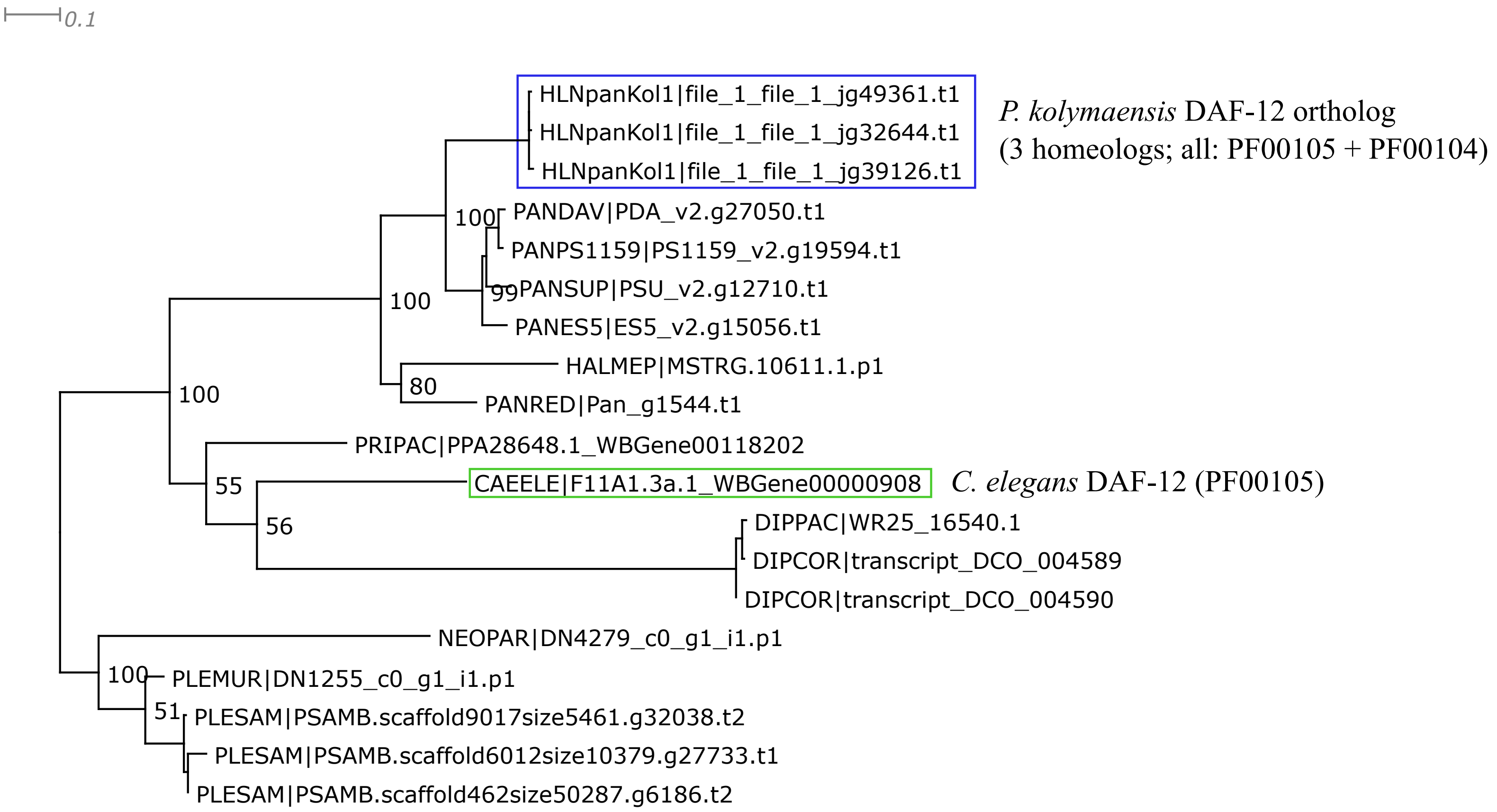


Trimal -automated1 function; short and spurious sequences removed manually.
IQtree2 ML phylogeny best-fit model according to BIC: LG+G4.

DAF-15

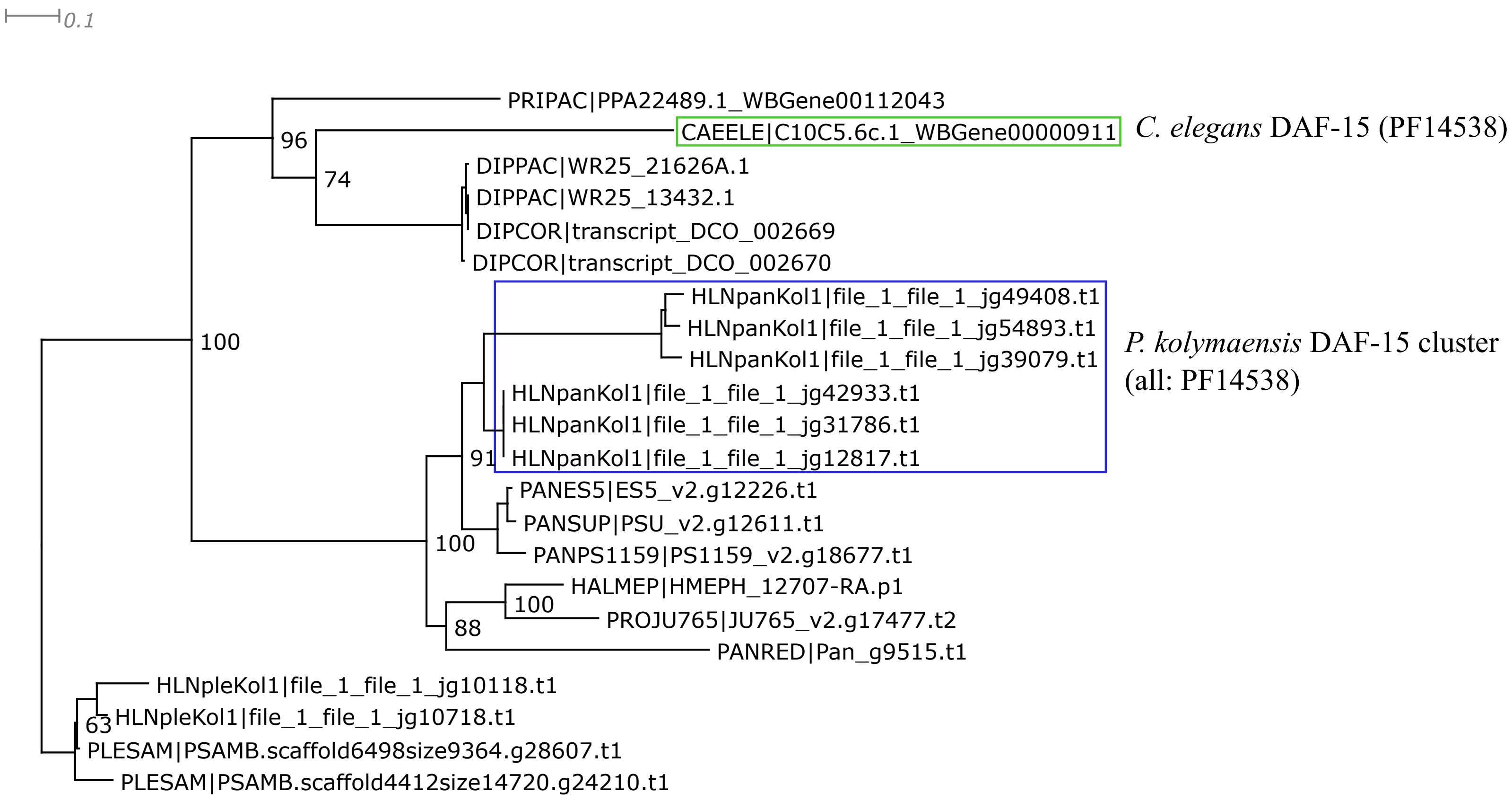


Trimal automated1 function; short and spurious sequences removed manually.
IQtree2 ML phylogeny best-fit model according to BIC: LG+G4.

DAF-16

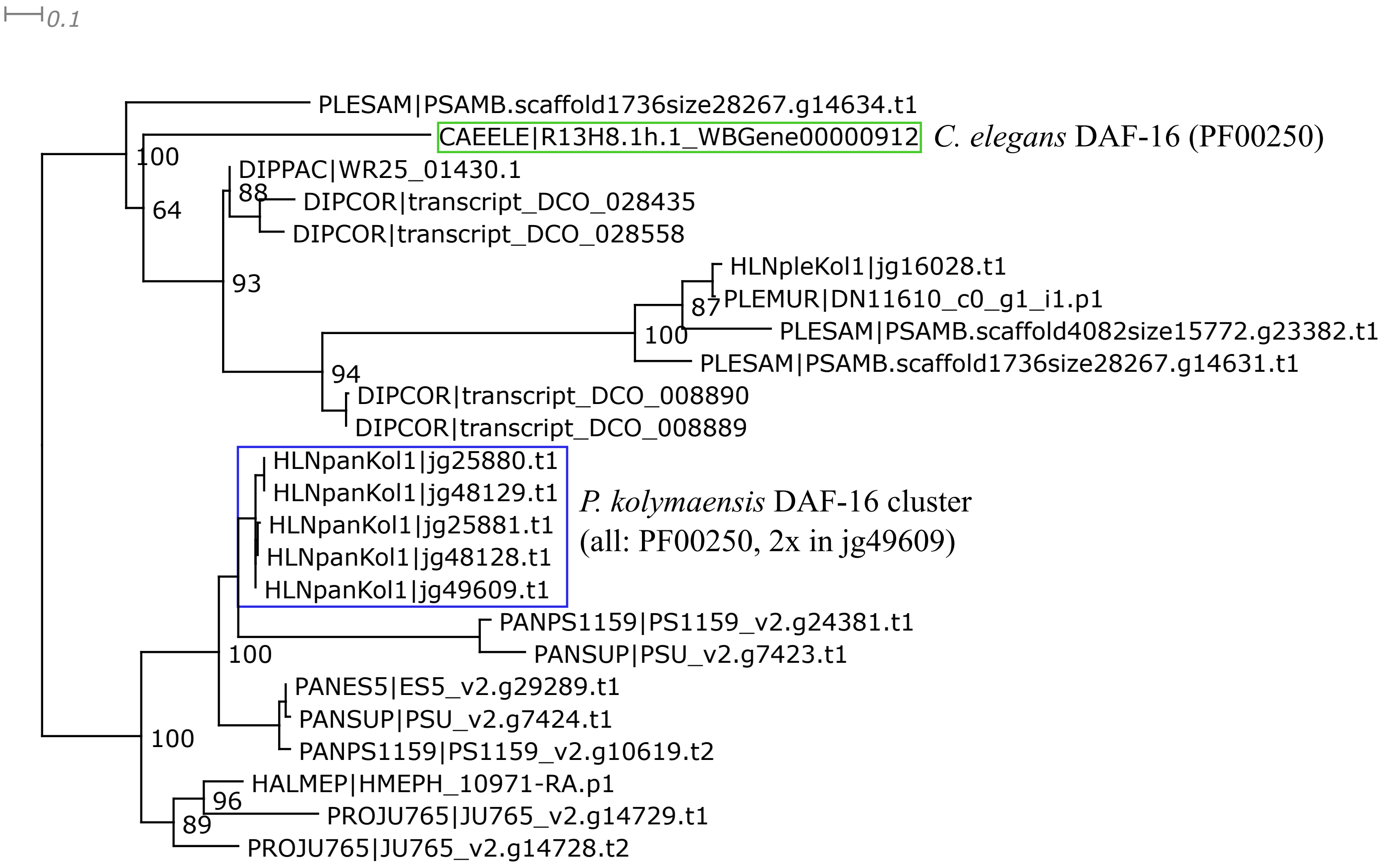


Short and spurious sequences removed manually.
IQtree2 ML phylogeny best-fit model according to BIC: VT+F+G4.

DAF-18

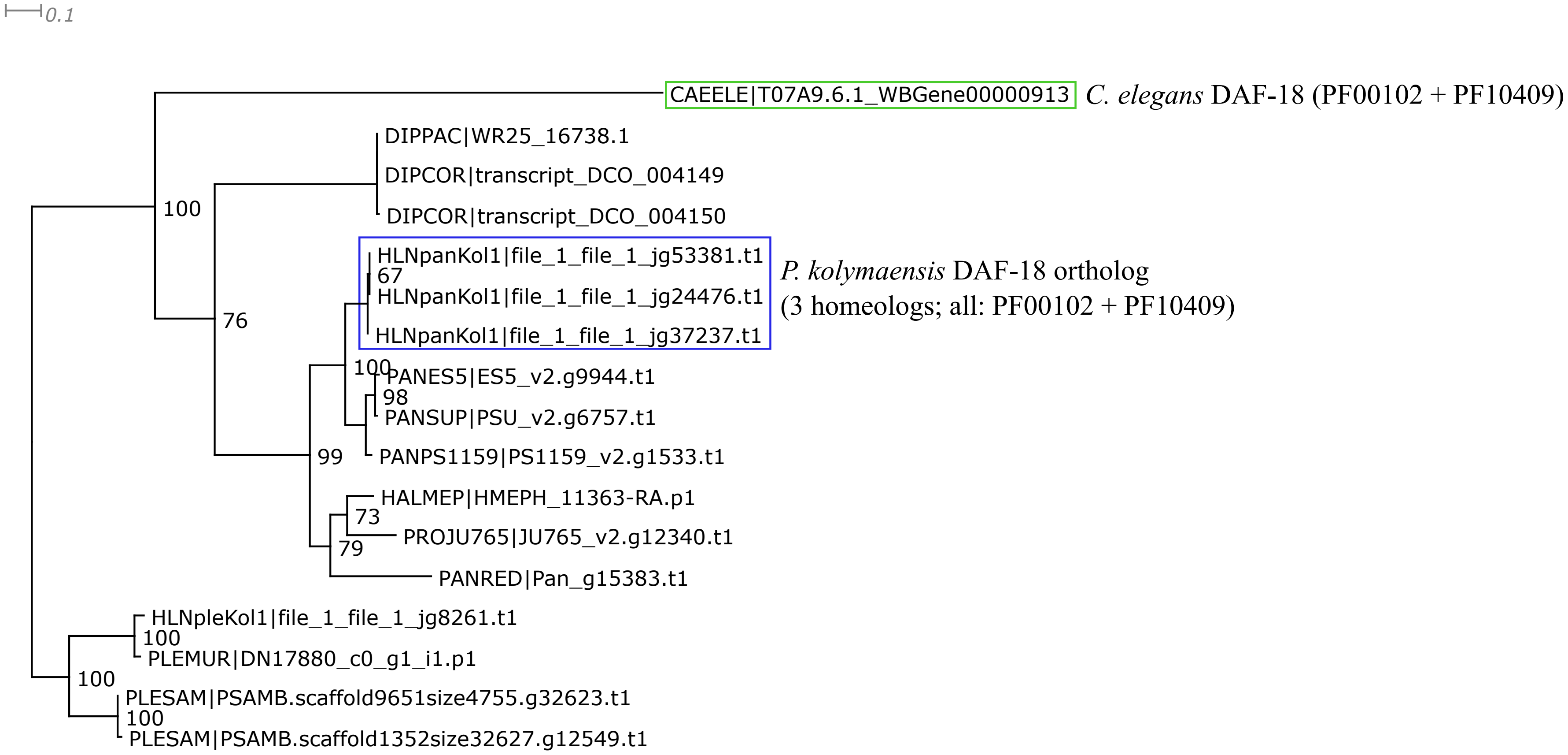


Trimal automated1 function: Trimal functions -resoverlap 0.75 -seqoverlap 75.
IQtree2 ML phylogeny best-fit model according to BIC: LG+I+G4.

DAF-19

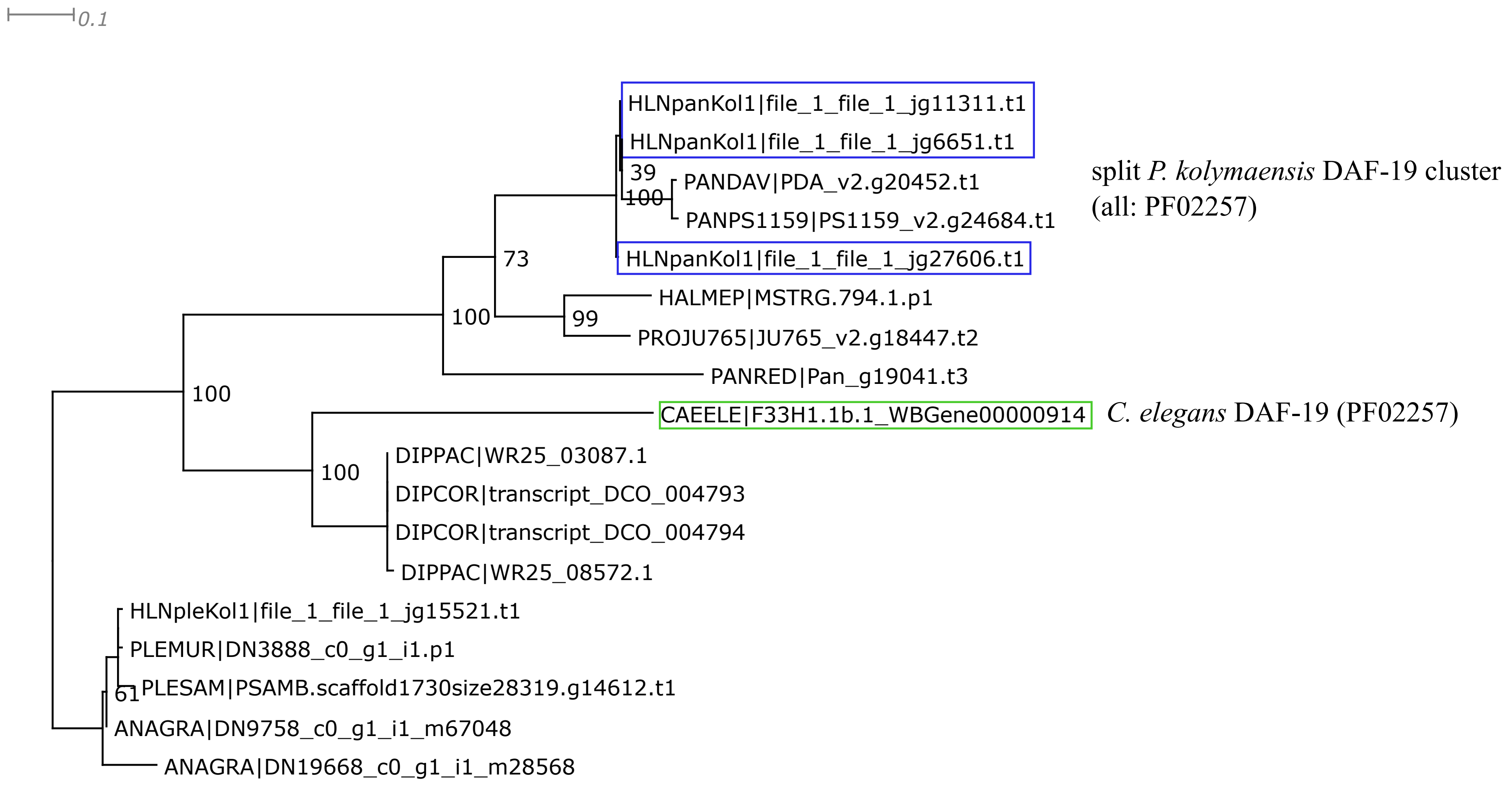


Trimal automated1 function; Short and spurious sequences removed manuallly.
IQtree2 ML phylogeny best-fit model according to BIC: LG+G4.

DAF-21

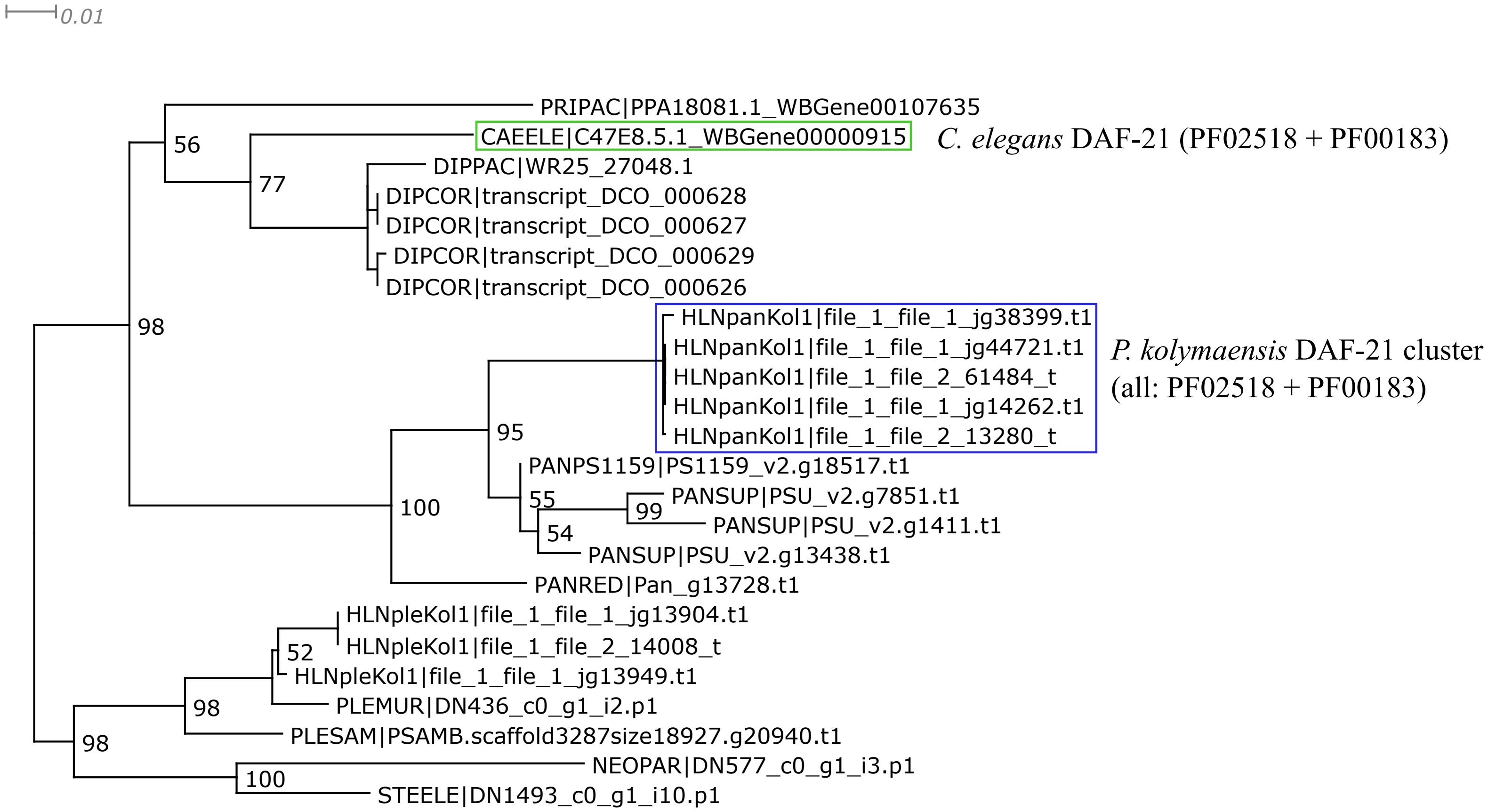


Trimal automated1 function; Short and spurious sequences removed manuallly.
IQtree2 ML phylogeny best-fit model according to BIC: LG+G4.

DAF-22

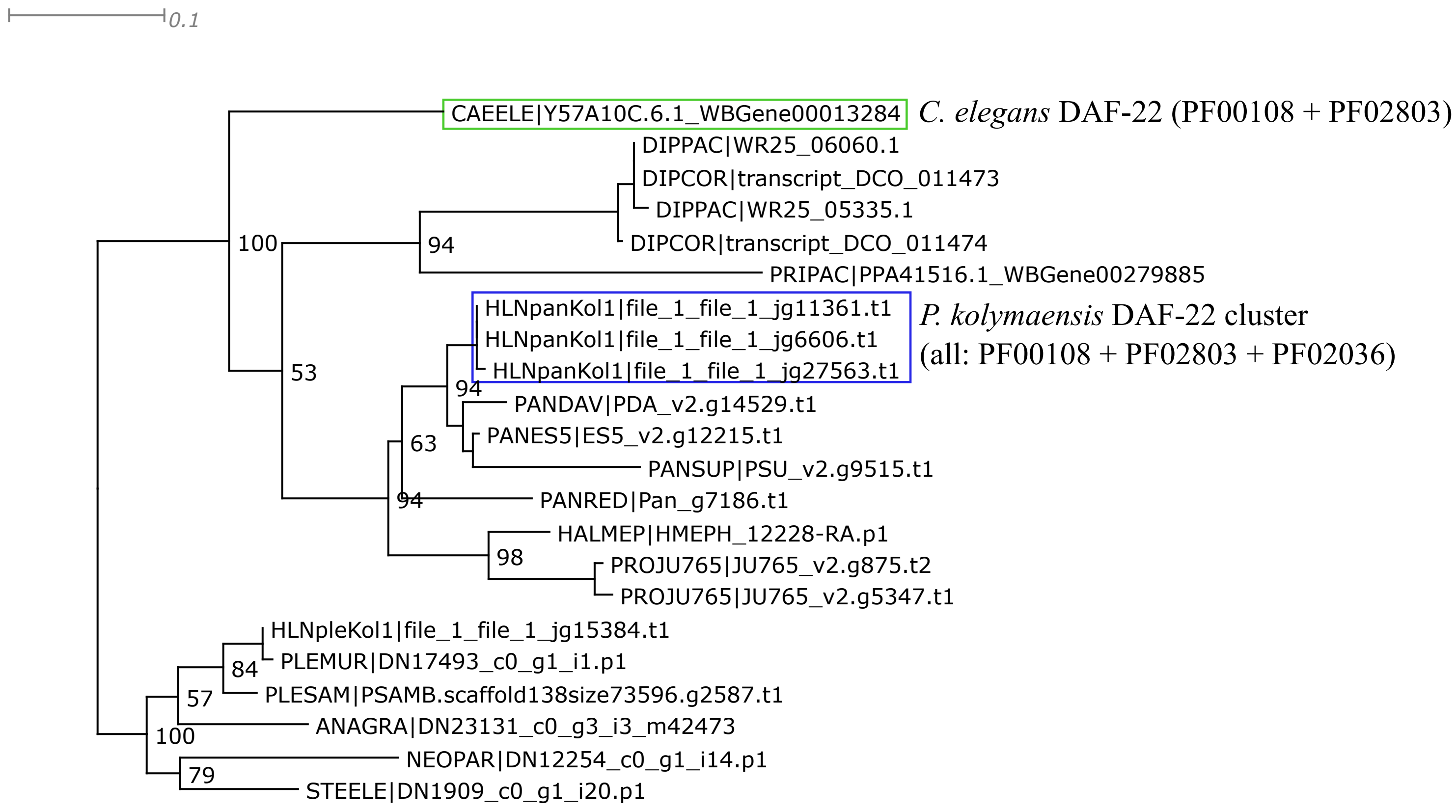


Trimal automated1 function; Short and spurious sequences removed manuallly.
IQtree2 ML phylogeny best-fit model according to BIC: LG+G4.

DAF-25

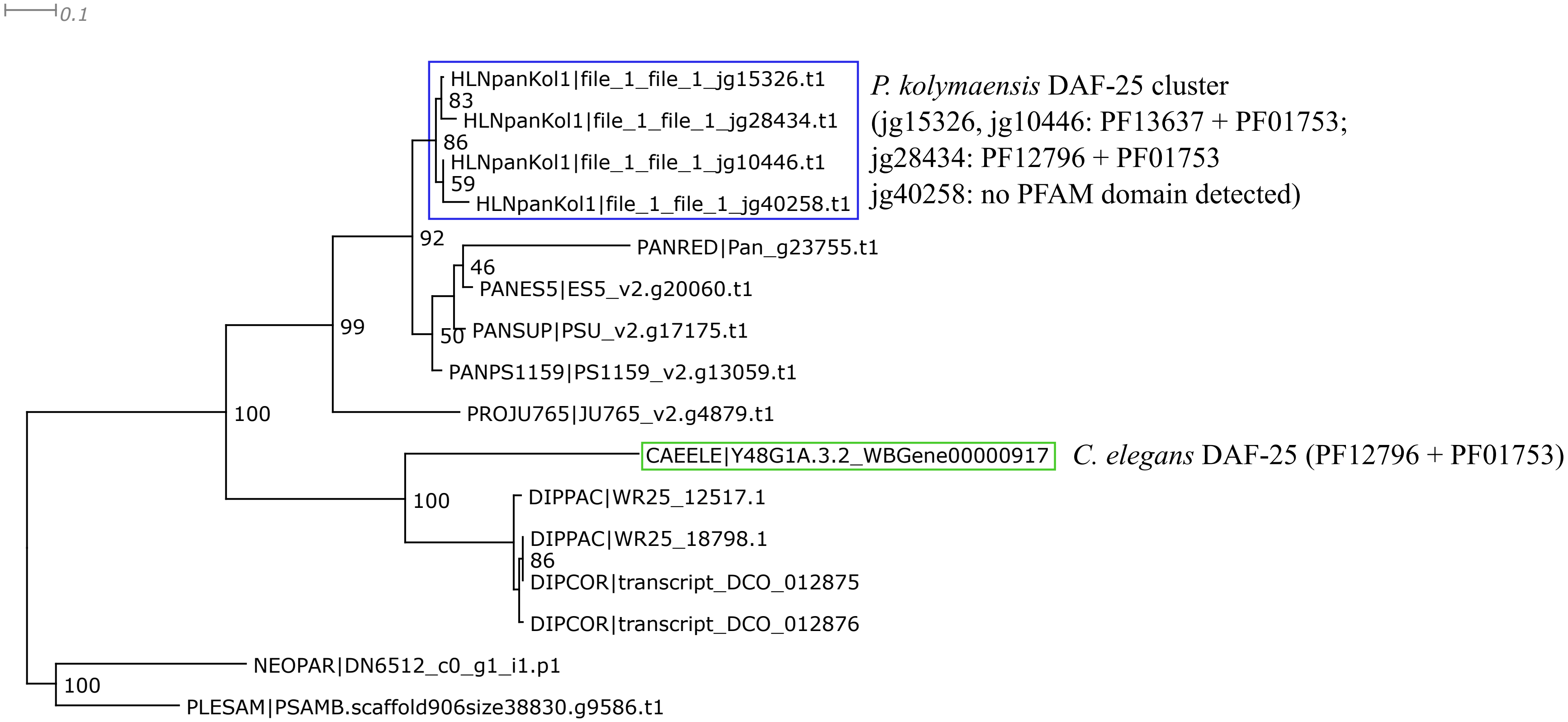


Trimal automated1 function; Short and spurious sequences removed manuallly.
IQtree2 ML phylogeny best-fit model according to BIC: LG+G4.

DAF-31

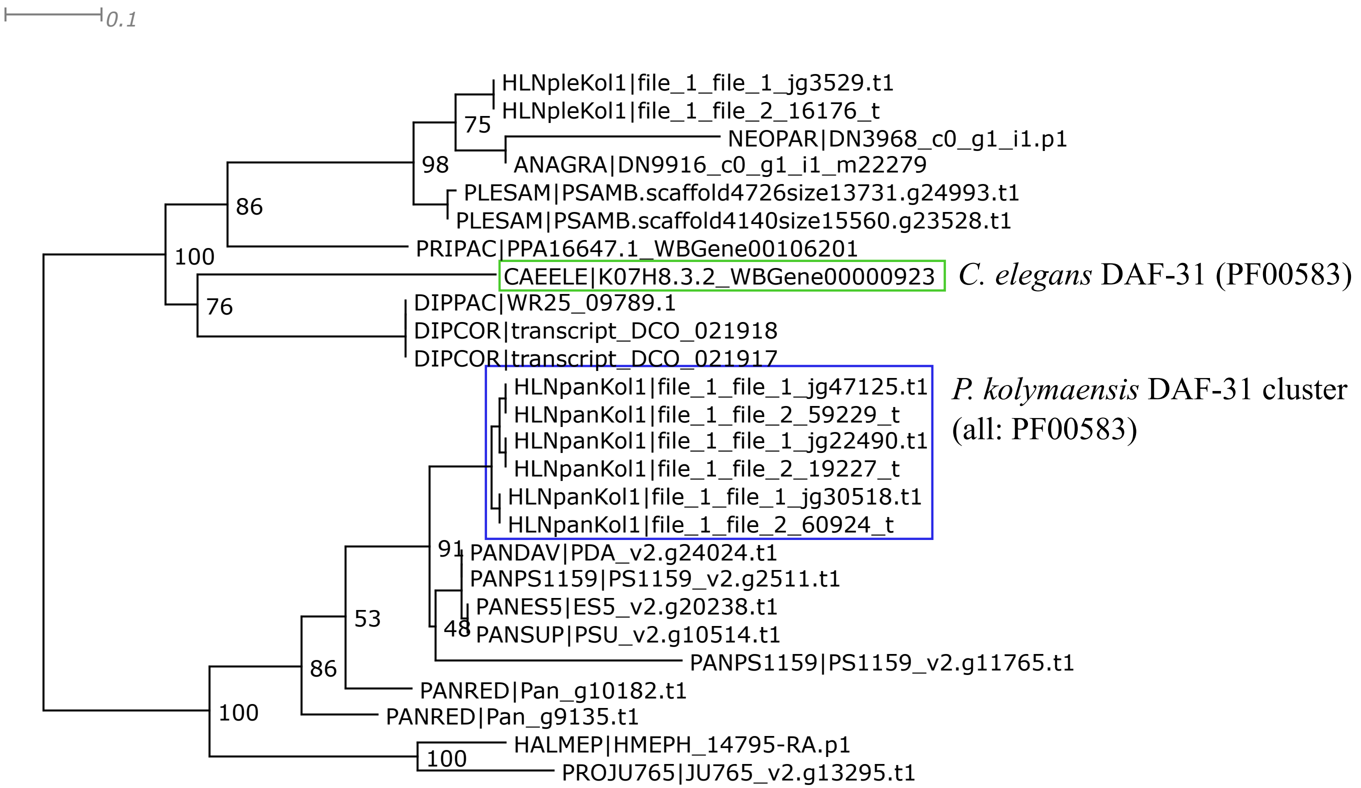


Trimal automated1 function; Short and spurious sequences removed manuallly.
IQtree2 ML phylogeny best-fit model according to BIC: LG+G4.

DAF-36

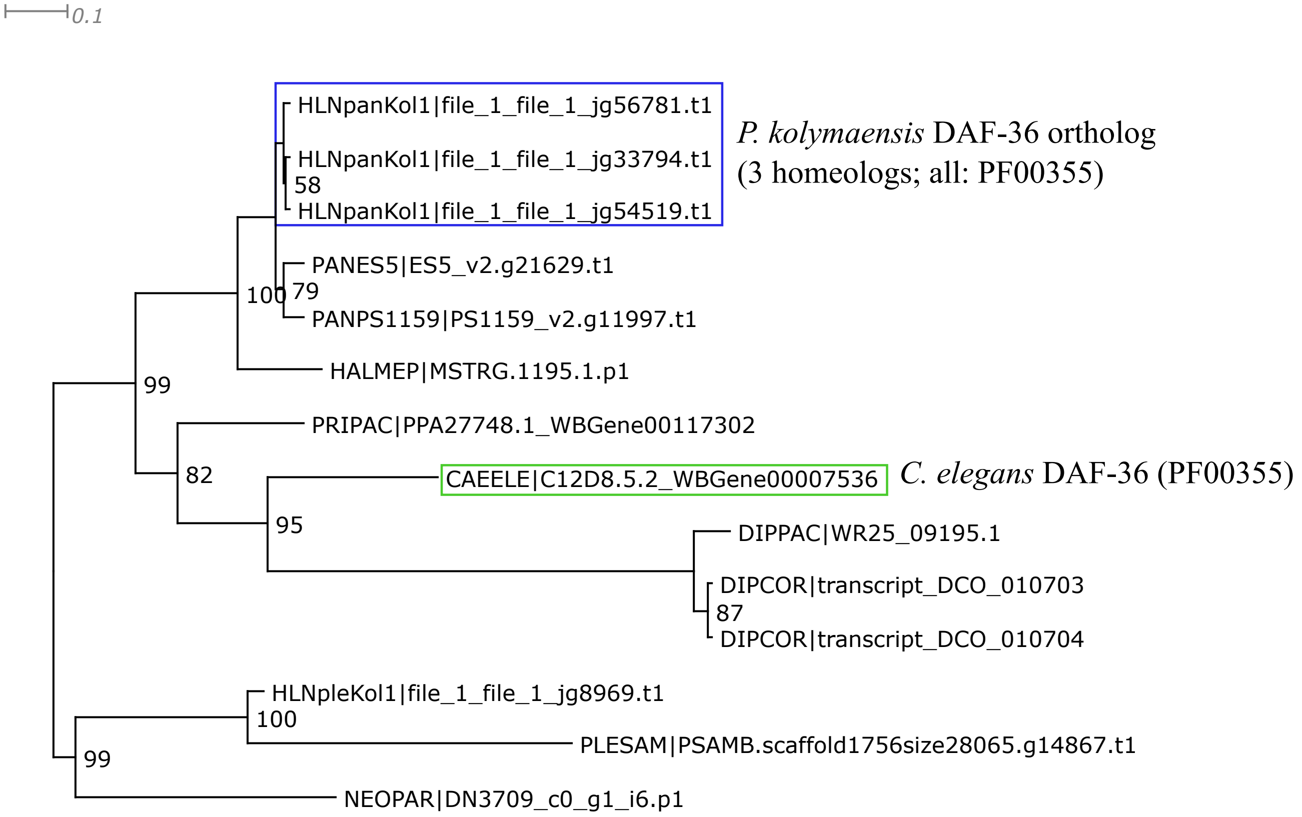


Trimal automated1 function; Short and spurious sequences removed manuallly.
IQtree2 ML phylogeny best-fit model according to BIC: LG+G4.

DAF-37

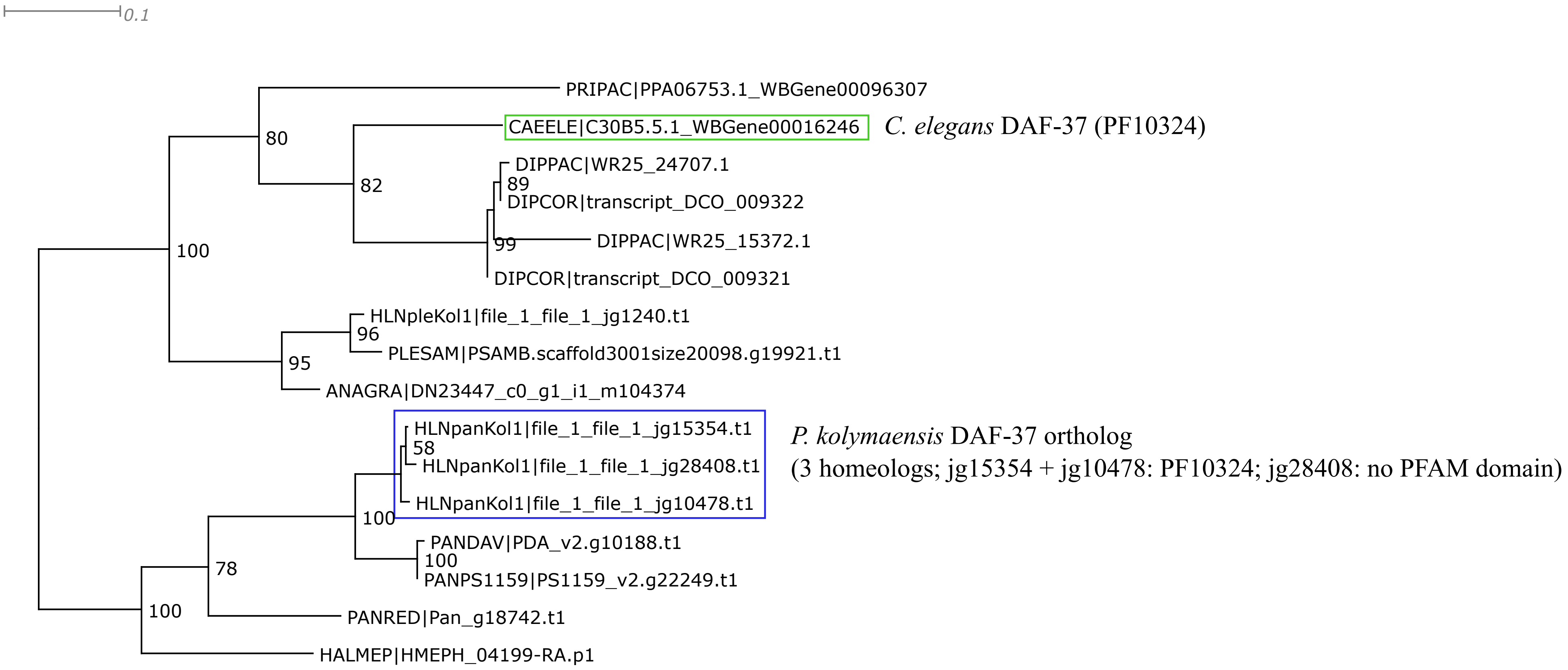


Trimal automated1 function; Short and spurious sequences removed manuallly.
IQtree2 ML phylogeny best-fit model according to BIC: LG+G4.

DAF-38

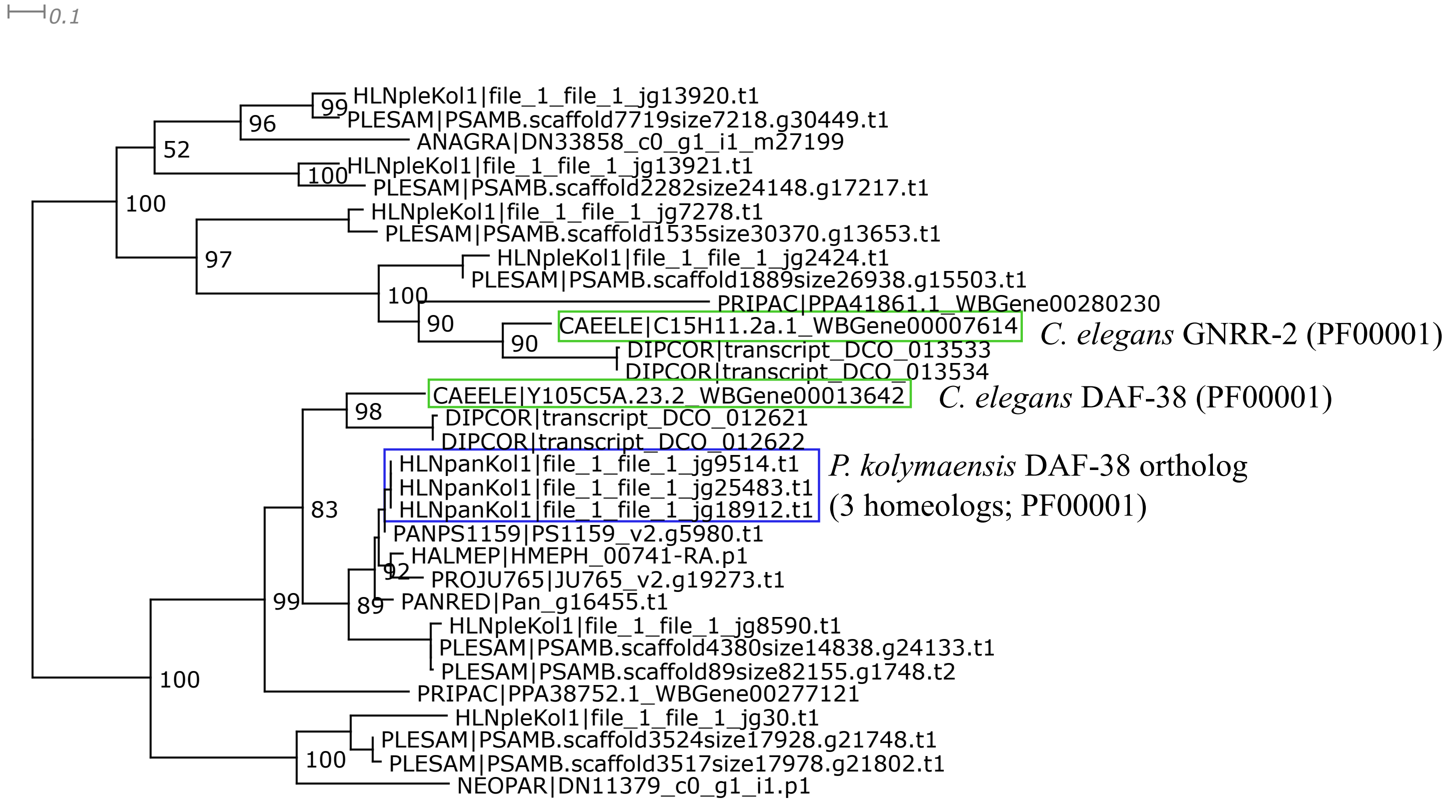


Trimal automated1 function; Short and spurious sequences removed manuallly.
IQtree2 ML phylogeny best-fit model according to BIC: mtInv+F+I+G4.

DAF-41

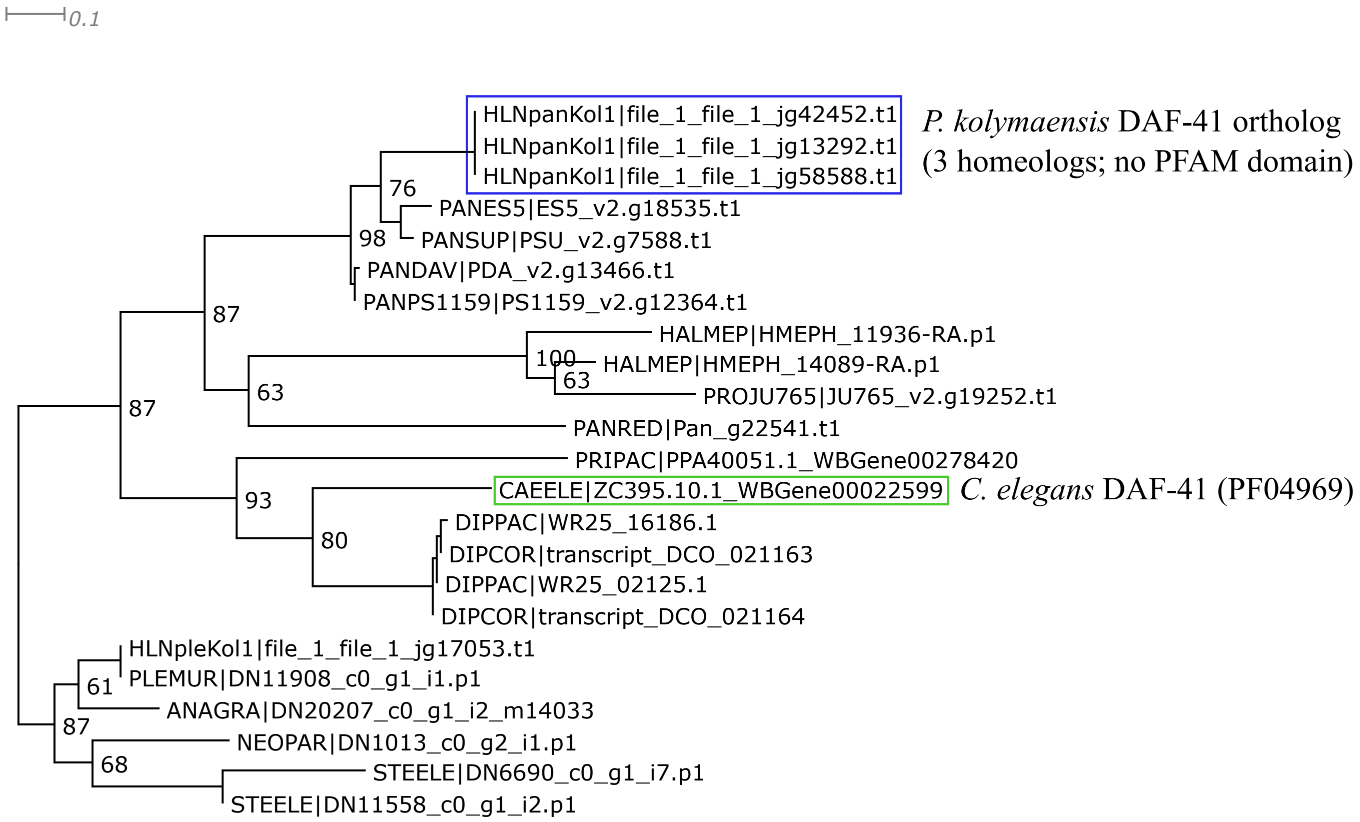


Trimal automated1 function; Trimal functions -resoverlap 0.8 -seqoverlap 80; Short and spurious sequences removed manuallly.
IQtree2 ML phylogeny best-fit model according to BIC: LG+G4.
Of all nematodes in this the phylogeny, the CS domain (PF04969) was only detected in *C. elegans* and some of the plectids.
